# Functional Right Ventricular and Atrial Cardiomyocytes Generated from Human Pluripotent Stem Cell-derived Second Heart Field Progenitors

**DOI:** 10.64898/2026.11.25.769301

**Authors:** Jianhua Zhang, Gina C. Kim, Vladislav Leonov, Hee Jae Jang, Jeanne Hsieh, Enson Zheng, Xuan Feng, Halle Fink, Amy Van Aartsen, Alexey V. Glukhov, Lee L. Eckhardt, Sean P. Palecek, Timothy J. Kamp

## Abstract

Second heart field (SHF) progenitors give rise to distinct populations of cardiomyocytes (CMs) in developing heart including right ventricular (rv) and atrial (a) CMs, but robust methods to generate these CM types from human pluripotent stem cells (hPSCs) are lacking. Here we differentiated hPSCs to SHF progenitors, rvCMs and aCMs identified by scRNA-seq transcriptome analysis in comparison to FHF-derived left ventricular (lv) CMs. The hPSC-CM single cell transcriptomes overlapped human fetal heart transcriptomes supporting the chamber-specific identity. The rvCMs strongly express the trabecular myocardium marker, *NPPA*, in contrast to the lvCMs which express *HEY2*, a compact myocardium marker. Functional studies demonstrated subtly different cardiac action potentials with a more prominent phase 1 repolarization in rvCMs and chamber-specific pharmacological responses. The gene expression pattern and functional properties suggest greater maturity of rvCMs relative to lvCMs. Overall, this study enables scalable access to chamber-specific hPSC-CMs for disease modeling and potential therapeutic applications.

## INTRODUCTION

Human pluripotent stem cells (hPSCs), including human embryonic stem cells (hESCs) and induced pluripotent stem cells (hiPSCs), have become an indispensable and powerful tool to understand human biology and disease ^1–3^. Growing emphasis on non-animal models to develop and test new cardiac therapies is increasing the use of hPSC-derived cardiomyocytes (hPSC-CMs) in pre-clinical model systems ^4,5^. In addition, clinical trials are beginning to test hPSC-CMs as a therapy for heart failure ^6,7^. Given the distinct embryonic, molecular and functional properties of cardiomyocytes present in different chambers of the heart, applications using hPSC-CMs require defined types of hPSC-CMs. For example, inherited heart disease can exhibit chamber-specific pathology: hypertrophic cardiomyopathy preferentially impacts the left ventricle (LV) ^8^, Brugada Syndrome pathology occurs in the right ventricle (RV) and right ventricular outflow tract (RVOT) ^9–12^, and arrhythmogenic cardiomyopathy (ACM) can predominantly impact the RV ^13^. Acquired heart disease such as atrial fibrillation, coronary artery disease and pulmonary hypertension can manifest chamber-specific pathophysiology. Congenital heart defects (CHDs) are specific to different regions of the developing heart and most commonly impact SHF-derived structures ^14,15^.

Heart development begins at gastrulation, during which cardiac progenitor cells (CPCs) leave the primitive streak (PS) and migrate anterior-laterally to form the cranial-cardiac mesoderm. Mesp1 is the earliest marker of CPCs. During mouse embryogenesis, Mesp1 expression starts at the onset of gastrulation (E6.5), continues until E7.5 and is then quickly down-regulated ^16–18^.

Studies using clonal analysis in mice identified two molecularly distinct CPC populations, the first heart field (FHF) and the second heart field (SHF) progenitors arising from two temporally distinct pools of Mesp1-expressing mesodermal progenitors at E6.5 and E7.25, respectively ^19,20^.

The FHF refers specifically to the first wave of mesodermal cells that differentiate to form the initial heart tube; The SHF consists of the progenitors that are continuously added to the formed heart tube at either the arterial or venous pole ^14^. FHF progenitors are unipotent and primarily give rise to CMs of LV, while SHF progenitors are multipotent and develop into RV and RVOT CMs, as well as non-myocytes including endothelial and smooth muscle cells ^19,21–27^. Both FHF and SHF progenitors contribute to atria. These differentiation trajectories are sculpted by varying gradients of morphogens including Wnts, FGFs, BMPs and retinoic acid (RA).

Identifying heart field progenitors and early progeny is handicapped by the lack of known and accessible markers. For example, molecular markers to define the FHF progenitors and distinguish the progenitors from the differentiated CMs are limited. Mutants of Nkx2-5 and Hand1 were found to primarily affect the LV structure during myocardial development and reflect the FHF lineage defects ^25,28–31^. Tbx5 mutant mice show severe defects in the atria and LV hypoplasia, indicating its critical roles in FHF progenitor proliferation and LV specification ^25,32,33^. The SHF is further defined as anterior SHF (aSHF) which develops into RV and outflow tract (OFT), and posterior SHF (pSHF) that contributes to atria. In contrast to the FHF, a number of molecular markers have been identified in developing SHF progenitors, including Islet-1 (Isl1), Tbx1, Foxh1, Mef2c, and Hand2, and fibroblast growth factors Fgf8 and Fgf10, as well as the most recently identified surface marker, CXCR4 ^21,25,34,35^. These genes are expressed in SHF progenitors, and most are downregulated upon differentiation to SHF lineages, suggesting progenitor-specific roles ^36^. RA is a crucial signaling morphogen in the pSHF region where Raldh2 is expressed to trigger atrial morphogenesis by activating the cardiac transcription factors Gata4 and Tbx5 ^37–39^.

Initial studies differentiating hPSCs to CMs did not identify or define the heart field- and chamber-specific hPSC-CMs ^40^. A recent study showed differentiation of heart field-specific lineages from mouse and human PSCs using reporter lines and identified the surface marker of CXCR4 to isolate SHF (CXCR4^+^) and FHF (CXCR4^-^) progenitors; however, differentiation of the heart field-specific progenitors to chamber-specific CMs was not described ^35^. Two other studies used the NKX2-5 and TBX5 double reporter lines to isolate hPSC-derived FHF and SHF lineages, and the differentiation protocols used in these studies were based on the biphasic Wnt modulation, the GiWi protocol, that mainly generated the FHF lineage; however, differentiation of the SHF progenitors to RV and OFT CMs, the major SHF lineages, was not elucidated ^41,42^. More recently Yang et al. ^43^ described heart field-specific lineage differentiation of hPSCs using suspended aggregates treated with different concentrations of Activin A and BMP4 to generate FHF (16B8A) and SHF (3B1.5A) mesodermal cell populations. However, this approach involved fluorescence-activated cell sorting (FACS) based on surface marker expression (CD235a/b, CXCR4, ALDH) to obtain the FHF, aSHF and pSHF progenitors, which is laborious, costly and not readily scalable. Another recent study from Mendjan’s group leveraged 2D and 3D hPSC differentiation approaches to generate FHF, aSHF and pSHF progenitors and integrated the progenitors to form multi-chamber cardioids to recapitulate early embryonic development. However, this approach for generating chamber-specific hPSC-CMs is also labor intensive and cannot readily generate large numbers of hPSC-CMs ^44^. Therefore, efficient and robust protocols to generate chamber-specific hPSC-CMs are needed to allow investigation of chamber-specific disease mechanisms and develop new therapies.

We recently published an efficient and scalable monolayer-based protocol to generate SHF progenitors by activating FGF signaling following mesoderm formation for differentiation of hPSCs to cardiac fibroblasts ^45^. Building on our SHF-specific differentiation protocol, in the present study we have developed methods to efficiently generate RV and atrial hPSC-CMs. We have performed thorough single-cell RNA sequencing and analyses to identify FHF and SHF progenitors as well as their differentiated chamber-specific hPSC-CMs from the newly developed SHF differentiation protocols as well as our previously developed GiWi protocol ^46,47^. Furthermore, we have performed comparative studies to compare our scRNA-seq dataset with published datasets of the hPSC-CMs and human fetal heart tissue. Functional analyses by single-cell patch-clamp, optical mapping and muscarinic cholinergic response confirmed the chamber-specificity of the hPSC-CMs derived from the SHF and the GiWi protocols. Moreover, the SHF-rvCM exhibited more mature cardiomyocyte phenotype at both molecular and functional levels.

## RESULTS

### Optimized GiWi protocol generates primarily FHF-derived ventricular hPSC-CMs

Biphasic modulation of Wnt signaling with small molecules (GiWi protocol) efficiently directs the differentiation of monolayer hPSCs to CMs ^46,47^. Recent studies using hPSC reporter lines for TBX5 and NKX2-5 suggested that biphasic Wnt modulation protocols produce both FHF (TBX5^+^/NKX2-5^+^) and SHF (TBX5^-^/NKX2-5^+^) lineages ^41,42^. Because the GiWi protocol is highly sensitive to the concentration of the GSK3β inhibitor, CHIR, in a cell line-dependent fashion, we systematically tested a range of 6 – 12 µM CHIR concentrations using the hES3-TBX5^TdTomato/W^/NKX2-5^eGFP/W^ reporter line ^42^. Flow cytometry analysis of the TBX5^TdTomato^, NKX2-5^eGFP^ and cTnT^+^ cells measured on days 0, 6, 10 and 15 of the GiWi protocol showed TBX5^+^ cells were present on day 6 with the highest percentage (46%) at 12 µM CHIR. In contrast, minimal NKX2-5^+^ cells were detected on day 6 (**Figure 1A,B**). The TBX5^+^ cell population continued to increase on day 10 (70%) when NKX2-5^+^ cells were also largely present (77%) along with the cTnT^+^ cells (80%) generated at the optimized 12 µM CHIR concentration (**Figure 1C,D**). The large population of NKX2-5^+^/TBX5^+^ cells (60%) indicated the generation of the FHF lineage ^41,42^. The TBX5 expression in the early differentiation of the GiWi protocol is consistent with mouse and chick heart development in which Tbx5 is first expressed throughout the entire cardiac crescent in early embryogenesis ^32^. However, the NKX2-5^+^/TBX5^-^cells defined as the SHF lineage in previous studies remained minimal ^41,42^ (**Figure 1D**). By day 15, the NKX2-5^+^/TBX5^+^ and cTnT^+^ cells remained similar to day 10, and the percentage of NKX2-5^+^/TBX5^+^cells varied in parallel with the cTnT^+^ cells with the highest average of ∼75% at the optimized 12 µM CHIR GiWi protocol (**Figure 1E,F**). The lower CHIR concentrations of 6-8 µM in the GiWi protocol generated significantly lower percentages of TBX5^+^/NKX2-5^+^ cells and cTnT^+^ cells (**Figure 1E,F**). The optimized CHIR concentration of 12 µM also showed the least batch-to-batch variation (**Supplementary Figure S1**). Overall, these results suggest that the monolayer differentiation protocol using biphasic Wnt modulation generates primarily FHF-derived hPSC-CMs at optimized CHIR concentration, consistent with another recent study ^48^.

**Figure 1.**
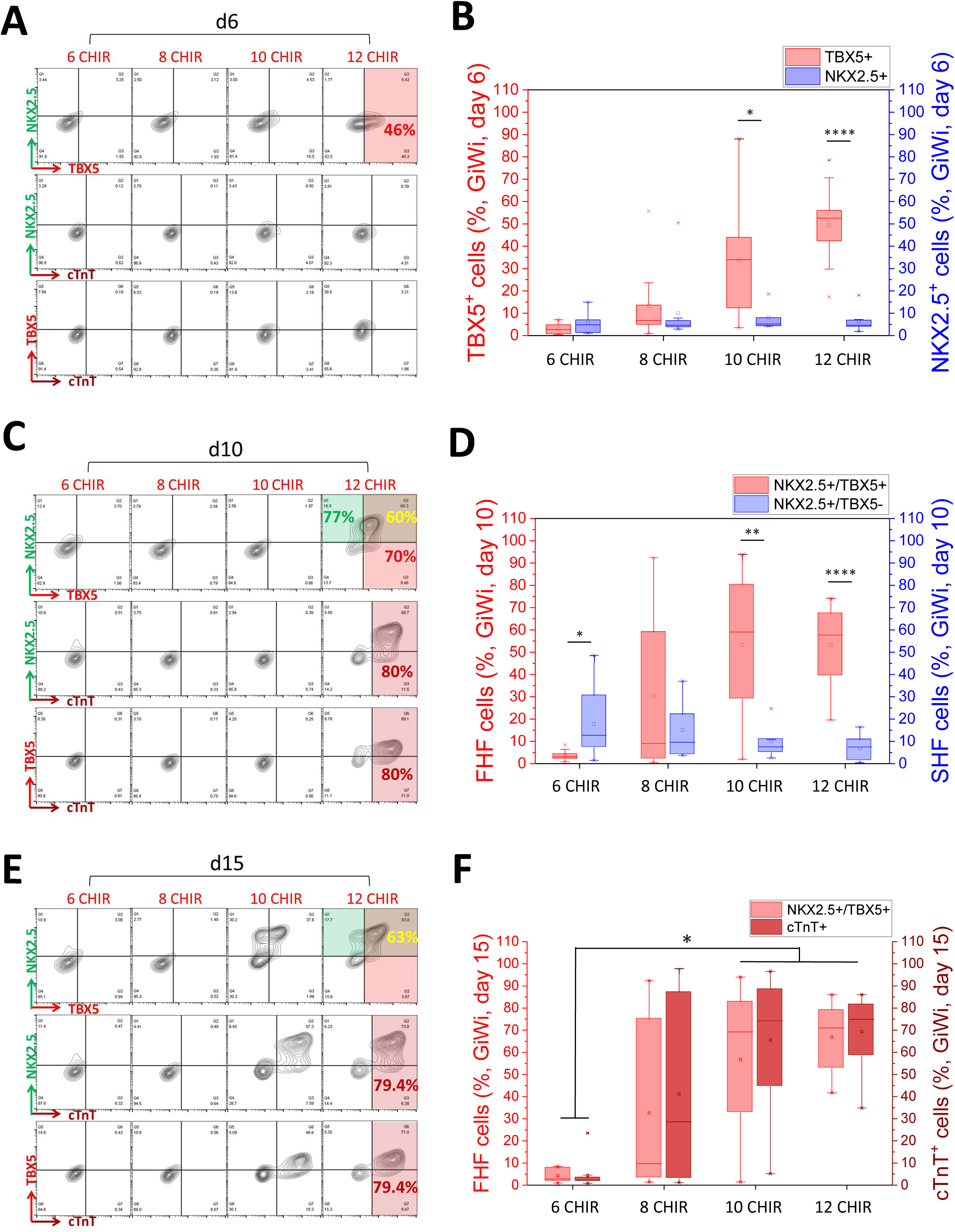
Optimized GiWi protocol using hES3-TBX5^TdTomato/W^/NKX2-5^eGFP/W^ reporter line generates primarily FHF-derived hPSC-CMs. (A) Flow cytometry analysis of TBX5, NKX2.5 and cTnT expressing cells on day 6 of the GiWi-protocol testing CHIR concentrations of 6-12 µM. (B) Average total TBX5^+^ or NKX2-5^+^ cells generated on day 6 of the GiWi protocol testing CHIR concentrations of 6-12 µM, measured by flow cytometry. N = 10 batches of differentiation as biological replicates. \**P*<0.05, **** *P*<0.0001, student t-test. (C) Flow cytometry analysis of TBX5, NKX2.5 and cTnT expressing cells on day 10 of the GiWi-protocol testing CHIR concentrations of 6-12 µM (continued differentiation trajectory from (A). (D) Average of total NKX2-5^+^/TBX5^+^ (FHF) and NKX2-5^+^/TBX5^-^(SHF) cells generated on day 10 of the GiWi protocol at CHIR concentrations of 6-12 µM, measured by flow cytometry. N = 10 batches of differentiation as biological replicates. \**P*<0.05, \*\**P*<0.005, **** *P*<0.0001, student t-test. (E) Flow cytometry analysis of TBX5, NKX2.5 and cTnT expressing cells on day 15 of the GiWi protocol testing CHIR concentrations of 6-12 µM (continued differentiation trajectory from (A) and (C)). (F) Average of total NKX2-5^+^/TBX5^+^ (FHF) and cTnT^+^ cells generated on day 15 of the GiWi protocol at CHIR concentrations of 6-12 µM, measured by flow cytometry. N = 10 batches of differentiation as biological replicates. \**P*<0.05, One-way ANOVA with Tukey post-hoc test. The box plots summarize the biological replicates with the box enclosing from first to third quartile, middle square indicating mean, line in box indicating median, and whiskers indicating outliers.

### Generation of SHF progenitors and derived hPSC-CMs

In contrast to FHF progenitors, SHF progenitors have distinct molecular features and properties manifested by continued proliferation and delayed differentiation. SHF development progresses from pharyngeal mesoderm where the cells are exposed to Wnt and FGF signaling to promote proliferation and maintain multipotency ^27^. As the SHF progenitors migrate to approach the developing heart tube, they are exposed to BMP and noncanonical Wnt to drive differentiation. The balance of proliferation and differentiation are regulated by a network of signaling pathways including canonical Wnt, FGF, BMP, noncanonical Wnt and retinoic acid signaling ^27^. We previously developed the GiFGF protocol to differentiate hPSCs to SHF lineage by activation of FGF signaling following mesoderm formation ^45^. Here, we use the hES3-TBX5^TdTomato/W^/NKX2-5^eGFP/W^ reporter line to verify the SHF progenitors generated in the GiFGF protocol by identifying the relative abundance of NKX2-5^+^/TBX5^-^cells ^41,42^. Flow cytometry showed the percentage of NKX2-5^+^ cells continued to increase from day 6 to day 10 and reached a peak of 52% of NKX2-5^+^/TBX5^-^cells at day 10 (**Supplementary Figure S2A**). However, the NKX2-5^+^/TBX5^+^ double positive cells, which represent the FHF lineage, exhibited the smallest population at day 9 (6%) (**Supplementary Figure S2A**). Given the sensitivity of the GiWi protocol to the CHIR concentrations, we also tested 6 - 12 µM CHIR in the GiFGF protocol.

Flow cytometry analysis for NKX2-5 and TBX5 positive cells showed 8 - 12 µM CHIR gave rise to similar percentage of NKX2-5^+^/TBX5^-^cells at day 9 with minor NKX2-5^+^/TBX5^+^ cells generated (**Supplementary Figure S2B,C**). Guided by developmental studies of SHF in mice ^27^, we next tested BMP and noncanonical Wnt signaling to drive the NKX2-5^+^/TBX5^-^ SHF progenitors to differentiate to CMs. The cells were treated with BMP4 (2.5-10 ng/ml) + Wnt 5a (100 ng/ml) for 4-10 days starting at day 9 in the CFBM+ medium (**Supplementary Table S1**), and the cells were differentiated for a total of 20 days. Flow cytometry analysis showed TBX5^-^/cTnT^+^ cells emerged as a small population in the 8 µM CHIR protocol on day 20, which indicates the developing SHF-derived CMs (**Supplementary Figure S2D**). However, treating the day 9 cells with BMP4 + Wnt 5a to promote differentiation of hPSC-CMs was problematic in our subsequent experiments because significant cell death occurred during the treatment. These results suggest that we might have missed the optimal timing for switching from the early phase proliferative and multipotent SHF progenitors to specification of cardiac fate by activating BMP and noncanonical Wnt signaling.

Using NKX2-5^+^/TBX5^-^ cells to identify early SHF progenitors is limiting because NKX2-5^+^ cells were not present until after day 4 (**Supplementary Figure S3**), after the initial phase of multipotent SHF progenitors. Early studies showed Nkx2.5 has multiple cis-regulatory elements that respond to different transcription factors to control gene expression in different regions of the developing heart ^49^. The failure of differentiating the SHF progenitors by BMP4 + Wnt 5a signaling when NKX2-5 is robustly expressed may be due to the previously described NKX2-5/BMP negative feedback loop ^50^. Thus, we hypothesized that the induction of CM fate in SHF occurs early, before NKX2-5 is significantly expressed. The SHF markers CXCR4 and ISL1 express earlier than NKX2-5, as shown at the mRNA level in our previous study ^45^ as well as by flow cytometry analysis of a time course of the GiFGF protocol (**Supplementary Figure S3**).

We then measured CXCR4^+^ cells in response to varying CHIR concentrations in the early progenitors of the GiFGF protocol (day 3.75 – 5.75) in comparison with the GiWi protocol, given the sensitivity of the monolayer-based differentiation protocols to the CHIR concentration. Flow cytometry analysis of the CXCR4^+^ cells showed a large population of CXCR4^+^ cells (67.4%) generated at day 3.75 with the optimized CHIR concentration of 8 µM, and the population of CXCR4^+^ cells gradually decreased by day 5.75 (**Figure 2A**). In contrast, only minimal CXCR4^+^ cells (9.40%) were generated at day 3.75 in the GiWi protocol with the optimized CHIR concentration of 12 µM. (**Figure 2A**).

**Figure 2.**
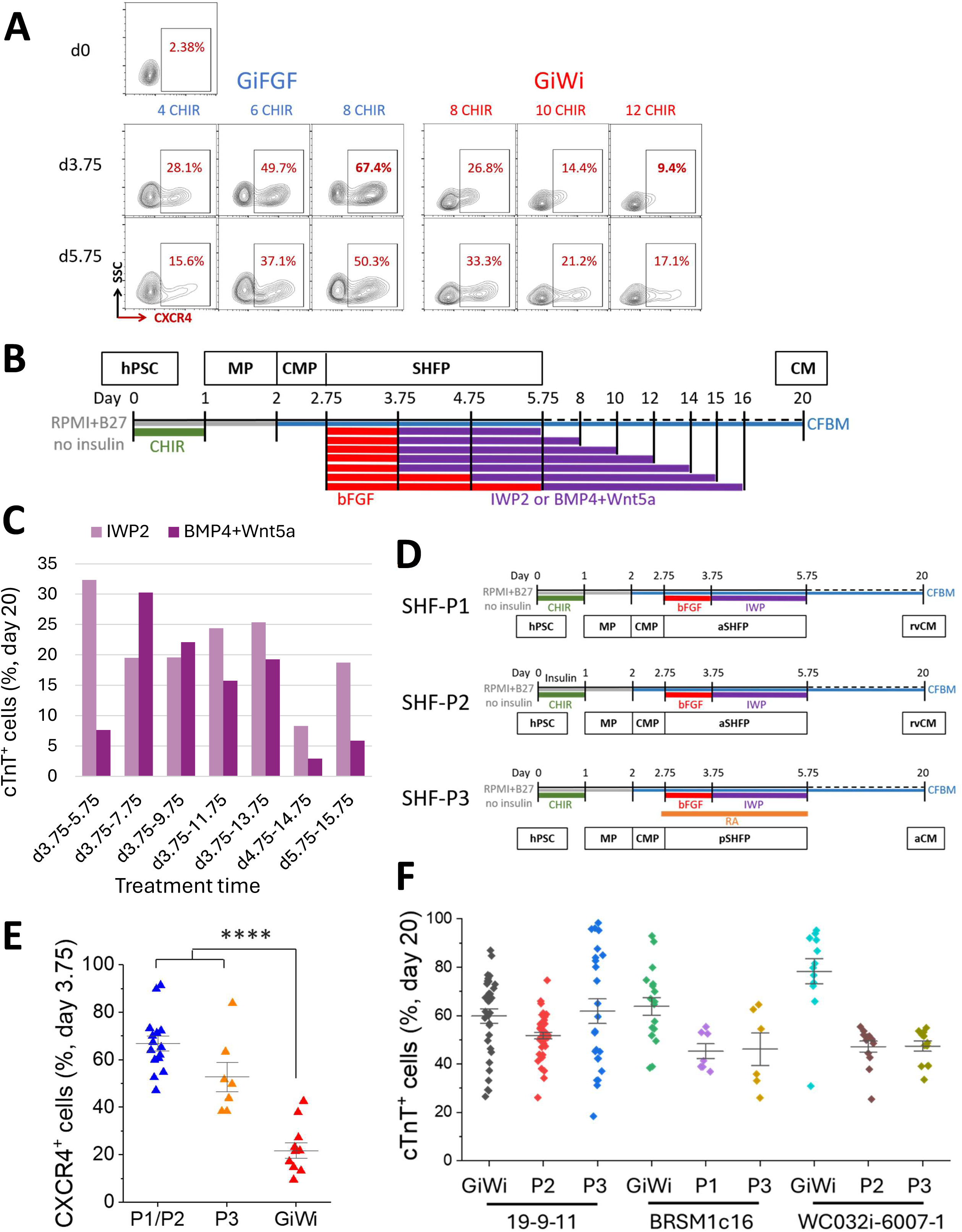
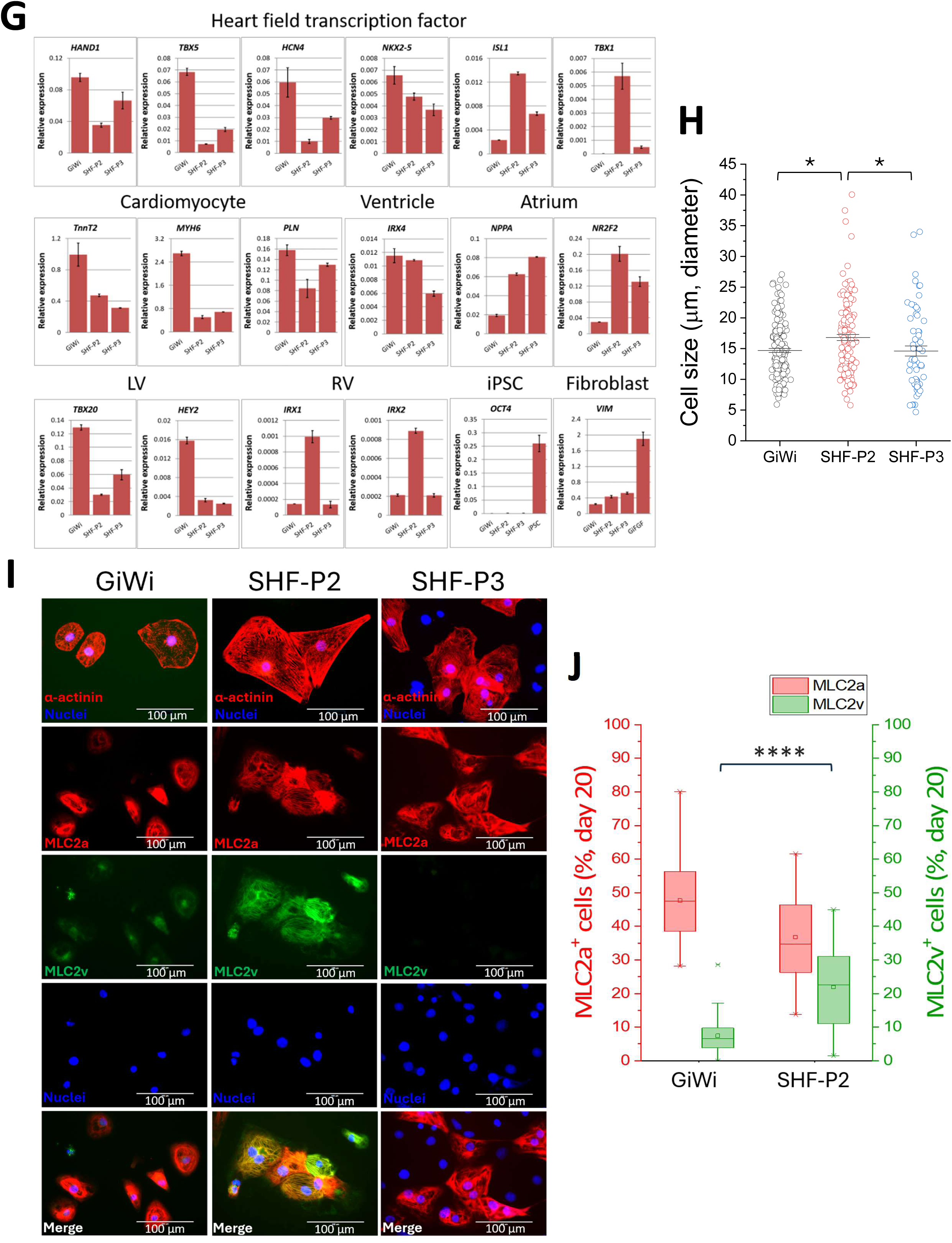
Generation of CXCR4^+^ SHF progenitors on day 3.75 following bFGF signaling and differentiation of SHF-derived hPSC-CMs. (A) Flow cytometry analysis of CXCR4^+^ cells on day 3.75 and 5.75 in GiFGF and GiWi protocols with varied CHIR concentrations for each protocol. (B) Schematic showing strategy to test inhibition of canonical Wnt with small molecule IWP2 or BMP4+Wnt5a (noncanonical Wnt) signaling to drive differentiation of SHF progenitors to hPSC-CMs. MP: mesodermal progenitor; CMP: cardiac mesodermal progenitor; SHFP: SHF progenitor. (C) Flow cytometry analysis of cTnT^+^ cells differentiated by the treatment of IWP2 or BMP4+Wnt5a in different time windows as shown in (B), following the generation of CXCR4^+^ SHFPs. (D) Schematics of the protocols to differentiate hPSC-rvCM and hPSC-aCM. (E) Average of CXCR4^+^ cells on day 3.75 in SHF-P1/P2, P3 and GiWi protocols, measured by flow cytometry. SHF-P1/P2, N = 15, SHF-P3, N=7 and GiWi protocol N = 11 biological replicates from DF19-9-11, BRSM1c16 and hES3-TBX5^TdTomato/W^/NKX2-5^eGFP/W^ lines. Scattered data plots with bars represent mean ± SEM. \*\*\*\**P*<0.0001, One-way ANOVA with Tukey post-hoc test. (F) Average of cTnT^+^ cells at 20 days differentiation using the GiWi, SHF-P1/P2 and P3 protocols measured by flow cytometry in multiple iPSC lines. DF19-9-11T, N = 32 (GiWi), 43 (P2) and 24 (P3). BRSM1c16, N = 18 (GiWi), 7 (P1) and 6 (P3). WC032i-6007-1, N =12 (GiWi), 13 (P2) and 11 (P3). Biological replicates. Scattered data plots with bars represent mean ± SEM. (G) Expression of heart field- and chamber-specific marker genes measured by qRT-PCR on the day 20 differentiated cells from DF-19-9-11T line using GiWi (73.2% cTnT^+^ CMs), SHF-P2 (32.3% cTnT^+^ CMs) and P3 (41.0% cTnT^+^ CMs) protocols. N=3 technical replicates. (H) Cell size measured on day 20 differentiated cells using GiWi, SHF-P2 and SHF-P3 protocols. N=182, 152 and 65 differentiated cells from DF19-9-11T line using GiWi, SHF-P2 and SHF-P3 protocols, respectively. \**P*<0.05, one-way ANOVA with Tukey post-hoc test. (I) Representative images of immunolabeling with sarcomeric α-actinin, myosin light chain 2a (MLC2a) and myosin light chain 2v (MLC2v) antibodies of the day 20 differentiated cells from GiWi, SHF-P2 and SHF-P3 protocols. (J) MLC2a^+^ and MLC2v^+^ cells (% of the total cells) counted on the MLC2a/MLC2v co-immunolabeling images as shown in (I). Total of 3766 and 2093 cells differentiated from GiWi and SHF-P2 protocols (7 differentiations in parallel) were analyzed, respectively. **** *P*<0.0001, student t-test.

To promote differentiation of SHF progenitors to right ventricular-like CMs (rvCMs) following the generation of the CXCR4^+^ cells at day 3.75, we tested adding BMP4 + Wnt5a to activate BMP and noncanonical Wnt signaling or IWP2 to inhibit canonical Wnt signaling. **Figure 2B** shows the schematic for an experiment testing BMP4 + Wnt5a treatment spanning different durations compared to the treatment with IWP2 alone. Wnt5a (100 ng/ml) + BMP4 (2.5 ng/ml) or IWP2 (5 µM) were added at day 3.75 and cells were treated for 2 to 10 days. Flow cytometry analysis of cTnT^+^ cells of the day 20 differentiated cells showed that treatment with BMP4 + Wnt5a for 4 days (day 3.75-7.75) or IWP2 (5 µM) for 2 days (day 3.75-5.75) resulted in a similar percentage of cTnT^+^ cells (**Figure 2C**). Longer treatment with either BMP4 + Wnt5a or IWP2 did not increase the cTnT^+^ cells (**Figure 2C**). Because the IWP alone or BMP4 + Wnt5a treatment yielded similar purity of the cTnT^+^ cells, we chose IWP treatment from day 3.75-5.75 for the SHF protocol to generate hPSC-rvCMs, given the lower cost of the small molecule than the growth factors. We also found insulin was necessary during day 0-1 for cell survival in some lines as in the GiWi protocol ^47^, thus the hPSC-rvCM differentiation protocols include SHF-P1 (no insulin) and SHF-P2 (with insulin) (**Figure 2D)**.

Generation of pSHF progenitors and atrial CM (aCM) requires retinoic acid (RA) signaling ^37–39,51^, so we tested adding RA (500 ng/ml) at day 2.75 when the SHF progenitors were generated, followed by IWP2 (5 µM) and RA (500 ng/ml) treatment for another 2 days to promote aCM differentiation, as shown in the SHF-P3 protocol (**Figure 2D).**

To assess the robustness and reproducibility of the SHF differentiation protocols, we performed series of differentiation using the SHF-P1, P2, P3 and optimized GiWi protocols running in parallel and in multiple cell lines. Flow cytometry analysis of the CXCR4^+^ cells (%) generated at day 3.75 in the SHF protocols with 8 µM CHIR including SHF-P1/P2 and P3 (with RA treatment at day 2.75-3.75) were comparable, but significantly higher than those in the GiWi protocol with 12 µM CHIR in multiple cell lines (**Figure 2E and Supplementary Figure S4**). Flow cytometry analysis of cTnT^+^ cells (%) at 20 days differentiation showed comparable hPSC-CM differentiation in SHF-P1, P2 and P3 protocols across the lines (**Figure 2F**).

To determine the heart field- and chamber-specificity of the hPSC-CMs differentiated using the SHF protocols, in comparison with the FHF-derived hPSC-CMs differentiated from the GiWi protocol, we evaluated the heart field- and chamber-specific gene expression by qRT-PCR of day 20 cells. **Figure 2G** shows *HAND1*, *TBX5* and *HCN4* for FHF were highly expressed in the GiWi-differentiated cells, while *ISL1* and *TBX1* for SHF and aSHF, respectively, were highly expressed in SHF-P2-differentiated cells. *NKX2*-5 expressed in both GiWi and SHF protocols-differentiated cells. The pan CM marker gene *TNNT2* expressed higher in GiWi than SHF protocols-differentiated cells likely due to the higher percentage of cTnT^+^ cells differentiated from the GiWi protocol relative to the SHF protocols (**Figure 2G)**. *IRX4*, a ventricular-specific transcription factor ^52,53^, was expressed comparably in GiWi and SHF-P2 but higher than the SHF-P3 differentiated cells, indicating the ventricular identity for both GiWi- and SHF-P2-CMs. Interestingly, both *NPPA* and *NR2F2* that are expressed in the atria and classified as atrial markers ^33,54–56^, expressed higher in both SHF-P2 and SHF-P3 than GiWi protocol differentiated cells, suggesting their expression may be upregulated during SHF development. *TBX20* and *HEY2* as FHF and ventricular, particularly LV markers ^54,57^ were highly expressed in the GiWi protocol-differentiated cells indicating the FHF lineage and LV identity. Furthermore, *NPPA* and *HEY2* as specific molecular markers for the trabecular and compact ventricular myocardium, respectively ^54,58,59^, were expressed differently in the GiWi and SHF-P2 protocols, may suggest a different subpopulation of ventricular CMs (compact vs. trabeculae) in the GiWi- and SHF-P2-differentiated CMs. *IRX1* and *IRX2* were highly expressed in the SHF-P2 protocol-differentiated cells suggesting the SHF lineage and RV identity. OCT4 expression was not observed in any of the protocol differentiated cells suggesting lack of undifferentiated iPSCs. Furthermore, *VIM* was expressed lower in all GiWi, SHF-P2 and P3 differentiated cells than the GiFGF protocol differentiated hPSC-cardiac fibroblasts ^45^, suggesting only minor fibroblast population present in these three hPSC-CM differentiation protocols (**Figure 2G**). We also evaluated the gene expression for the myofilament protein isoforms that reflect the maturity of the hPSC-CMs ^60^.

*TNNT2* expression was comparable when comparing GiWi and SHF-P2 differentiated ventricular hPSC-CMs, while expression of *TNNI3*, *MYL2* and *MYH7* were significantly higher in SHF-P2-than the GiWi-differentiated cells, suggesting a more mature phenotype of SHF-P2-rvCMs than GiWi-lvCMs (**Supplementary Figure S5**). Furthermore, we measured cell size of the day 20 differentiated cells from each protocol. Phase contrast images were taken when harvesting the cells, and cell diameters were measured using ImageJ. The average cell diameter of the suspended cells from the SHF-P2-differentiated cells on day 20 (16.8±0.49 µm) was significantly greater than from the GiWi-(14.7±0.49 µm) and SHF-P3-(14.6±0.82 µm) differentiated cells (**Figure 2H**).

Immunolabeling of day 20 differentiated hPSC-CM populations from GiWi, SHF-P2 and SHF-P3 protocols with antibodies to myofilament proteins including sarcomeric α-actinin, MLC2a and MLC2v was also performed to probe the cell identity and maturity. The hPSC-CMs differentiated from all three protocols showed strong expression of α-actinin, and the striated pattern of α-actinin was more evident in GiWi- and SHF-P2-CMs than the SHF-P3-CMs (**Figure 2I**). MLC2a^+^ cells were observed in all three protocol-CMs with more robust expression in SHF-P3-CMs, but variable expression from cell to cell in GiWi- and SHF-P2-CMs (**Figure 2I**). In contrast, MLC2v^+^ cells were observed in both GiWi- and SHF-P2-CMs but not in SHF-P3-CMs (**Figure 2I**). These results are consistent with the ventricular identity of GiWi and SHF-P2 CMs and atrial-like cells of SHF-P3-CMs. Moreover, quantitative measurement by counting the MLC2a^+^ and MLC2v^+^ of the immunolabeled cells showed significantly higher percentage of MLC2v^+^ cells in SHF-P2-CMs than GiWi-CMs (**Figure 2J**), consistent with the gene expression measured by qRT-PCR (**Supplementary Figure S5**). Overall, these results suggest the GiWi protocol generates primarily lvCMs, while the SHF-P2 and P3 protocols generate rvCMs and aCMs, respectively. Furthermore, the different expression of the myofilament protein isoforms and the increased cell size in the SHF-P2-rvCMs suggest a more mature phenotype than the GiWi-lvCMs.

### Heart field-specific lineages defined by single-cell transcriptomics

To better understand the heart field- and chamber-specific hPSC-CMs differentiated using the described protocols, we performed scRNA-seq of the DF19-9-11T hiPSC-differentiated day 3.75 progenitors and day 20 cells from the GiWi, SHF-P2 and SHF-P3 protocols. We first confirmed successful differentiation from all three protocols by flow cytometry to analyze the SHF progenitors (CXCR4^+^ cells) on day 3.75 and cTnT^+^ cells on day 20 (**Supplementary Figure S6**). A total of 20,000 cells from two progenitor samples (day 3.75) from the GiWi-12 µM CHIR and SHF-P2 protocols and four samples of day 20 differentiated cells of GiWi-12 µM CHIR, GiWi-8 µM CHIR, SHF-P2 and SHF-P3 protocols were included as a single batch experiment for the scRNA-seq. Transcriptomic analysis of all 6 samples revealed 19 clusters which were identified as 16 cell types by marker gene expression, differential gene expression, and Gene Ontology (GO) analysis (**Figure 3A**). The day 3.75 progenitors generated in the GiWi −12 µM CHIR protocol were identified as predominantly FHF progenitors by the marker genes *DCC*, *GYPB* and *HAND1* along with other highly expressed DEGs (**Figure 3B and Supplementary Figure S7**), and a small cell population of definitive endoderm progenitors (DE-1) that differentially express *FOXA2* and *SOX17* (**Figure 3B**). The day 3.75 progenitors generated in the SHF-P2 protocol were heterogenous, including SHF, definitive endoderm (DE-2) and primitive streak (PS) progenitors identified by the marker genes of *CXCR4*, *LHX1* and *FOXC1* for SHF, *FOXA2* and *SOX17* for DE, and *TBXT* and *POU5F1* for PS, respectively (**Figure 3B**), along with other highly expressed DEGs (**Supplementary Figure S7**). The day 20 GiWi-differentiated cells were identified primarily as lvCM of the FHF lineage by DEGs of *HAND1*, *IRX4*, *HEY2*, *MYL2*, *MYH6*, *MYH7* and *CACNA1C* (**Figure 3C)**. Furthermore, the GiWi-lvCM were further divided into immature (*MYL2*^low^/*MYH6*^high^/*MYH7*^low^), mature (*MYL2*^high^/*MYH6*^low^/*MYH7*^high^) and proliferating (*MKI67*^+^/*TOP2A*^+^) lvCM (**Figure 3A,D**). The day 20 SHF-P2 protocol-derived cells were identified mainly as rvCM by the DEGs of *NPPB*, *ABRA*, *MEF2C*, *RRAD*, *CMYA5*, *MYL2*, *MYH7*, *A2M* and *CACNA1D* (**Figure 3E**). The day 20 SHF-P3 protocol-derived cells were identified as two major hPSC-CM populations: aCM (*NR2F2* and *VWDE*) and sinoatrial nodal (SAN) CM (*SHOX2* and *TBX18*) (**Figure 3F**). Other cell types in the day 20 samples include epicardial cells, cardiac fibroblast (CF) and epithelial cells (**Figure 3A**). Moreover, the day 20 differentiated cells from GiWi-12 µM CHIR and GiWi-8 µM CHIR overlapped in the clustering, indicating that the different CHIR concentrations did not change the cell fate (**Figure 3A**).

**Figure 3.**
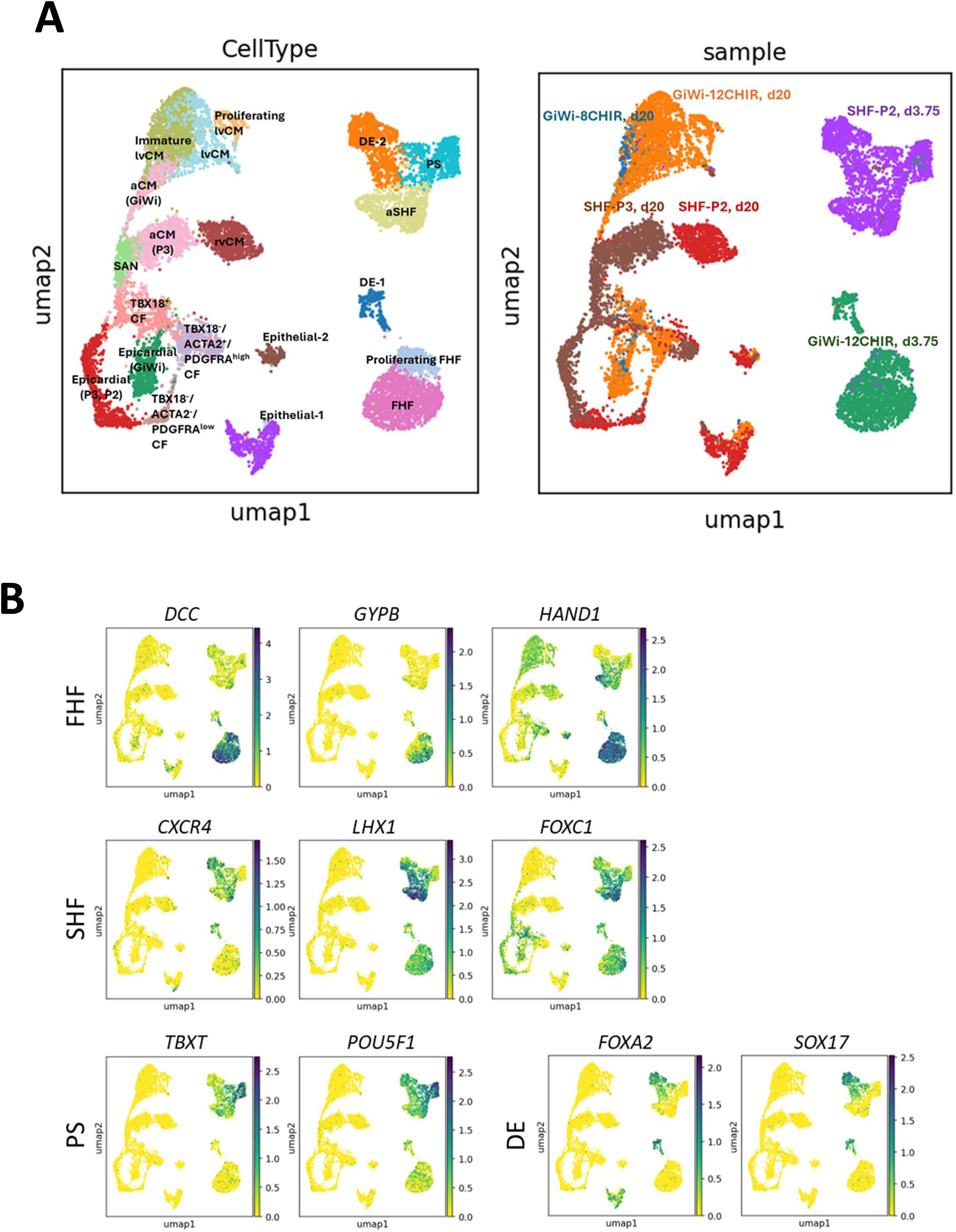
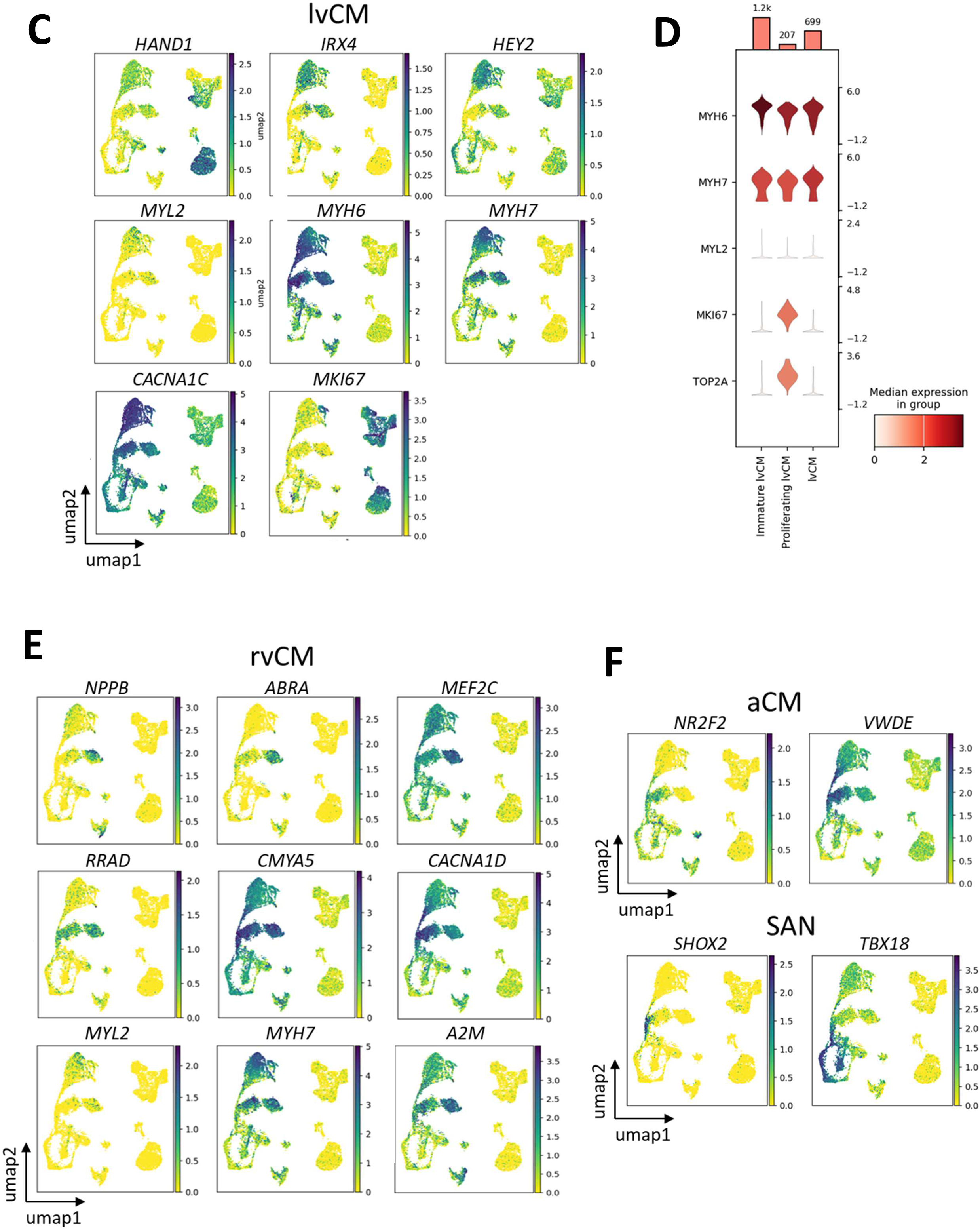
Single-cell transcriptomic analysis of day 3.75 progenitors and day 20 differentiated cells in the SHF-P2, SHF-P3 and GiWi protocols. (A) UMAP of a total of 12,910 cells from 6 samples of 1) SHF-P2, d3.75; 2) GiWi-12CHIR, d3.75; 3) GiWi-12CHIR, d20, 4) GiWi-8CHIR, d20, 5) SHF-P2, d20; 6) SHF-P3, d20. Left, UMAP with annotated cell type; Right, UMAP with sample view. Clustering resolution is 0.8. (B) Embedding plots of representative marker genes expression of first heart field (FHF), second heart field (SHF), primitive streak (PS) and definitive endoderm (DE) progenitors in log_2_FC. (C) Embedding plots of representative marker genes expression of hPSC-lvCM in log_2_FC. (D) Violin plot of the differential expression of *MYH6*, *MYH7*, *MYL2*, *MKI67* and *TOP2A* in immature lvCM, lvCM and proliferating lvCM differentiated from the GiWi protocol. Bars represent cell number for each cluster. Y-axis of violin plot represents log_2_(TPM+1) values for gene expression. (E) Embedding plots of representative marker genes expression of hPSC-rvCM in log_2_FC. (F) Embedding plots of representative marker genes expression of hPSC-aCM and hPSC-SAN in log_2_FC.

### Single-cell transcriptomics of the day 3.75 progenitors

We further analyzed the scRNA-seq data comparing the day 3.75 progenitors generated from SHF-P2 and GiWi protocols (**Figure 4A**). Comparison of the transcriptomes revealed the two major and closely related cell populations from the GiWi protocol (clusters 0, 2) being FHF progenitors defined by the DEGs including *BMPER*, *DCC*, *HS3ST3A1*, *LGR5*, *NRG*2, *PDGFRA*, *HAND1*, *MEIS1*, *LIX1*, *GATA6*, *GYPB* and *TBX20* in both clusters (**Figure 4A,B**). Further comparison of the DEGs in cluster 0 and 2 showed relatively higher expression of *MKI67*, *TOP2A*, *PODXL*, *TNC*, *VIM* and *PKP2* in cluster 0 than cluster 2, suggesting cluster 0 composing earlier and more proliferative FHF progenitors than those in cluster 2 (**Supplementary Figure S8**). In contrast, the SHF-P2 protocol-derived progenitors are more heterogenous with 5 cell populations identified as anterior SHF (aSHF), definitive endoderm (DE-2), primitive streak (PS), mesendoderm progenitors and mesenchymal cells (**Figure 4A**). A large cell population (cluster 1) from the SHF-P2 protocol-derived day 3.75 progenitors was identified as aSHF with upregulated expression of DEGs including *LHX1*, *APLNR*, *VAV3*, *LIPG*, *DLL3*, *DACT1*, *FOXC1*, *TBX6*, *CYP26A1* and *CXCR4* (**Figure 4C**). Clusters 3 and 6 from GiWi and SHF-P2 protocols, respectively were identified as DE progenitors by expression of the marker genes *FOXA2* and *SOX17* (**Figure 4D**). The PS cells (cluster 4) highly expressed the DEGs *TBXT* and *POU5F1* (**Figure 4D**). Cluster 5 comprising mostly cells derived from SHF-P2 protocol and a small portion from GiWi protocol was identified as mesendoderm progenitors that express both the endoderm marker genes of *FOXA2* and *SOX17* and the DEGs expressed in both FHF and SHF mesoderm cells, such as *BMPER* and *PDGFRA* (FHF), *LHX1* and *APLNR* (SHF) (**Figure 4D,E**). A small cell population (cluster 7) expressed *PRRX1* as the most significant DEG in this cluster was identified as mesenchymal cell (**Figure 4D**). Furthermore, most of the progenitor populations highly expressed *MKI67* and *TOP2A* indicating the proliferative ability of the early progenitors (**Figure 4D,E)**. The DEGs that define each progenitor population are shown in **Figure 4E**.

**Figure 4.**
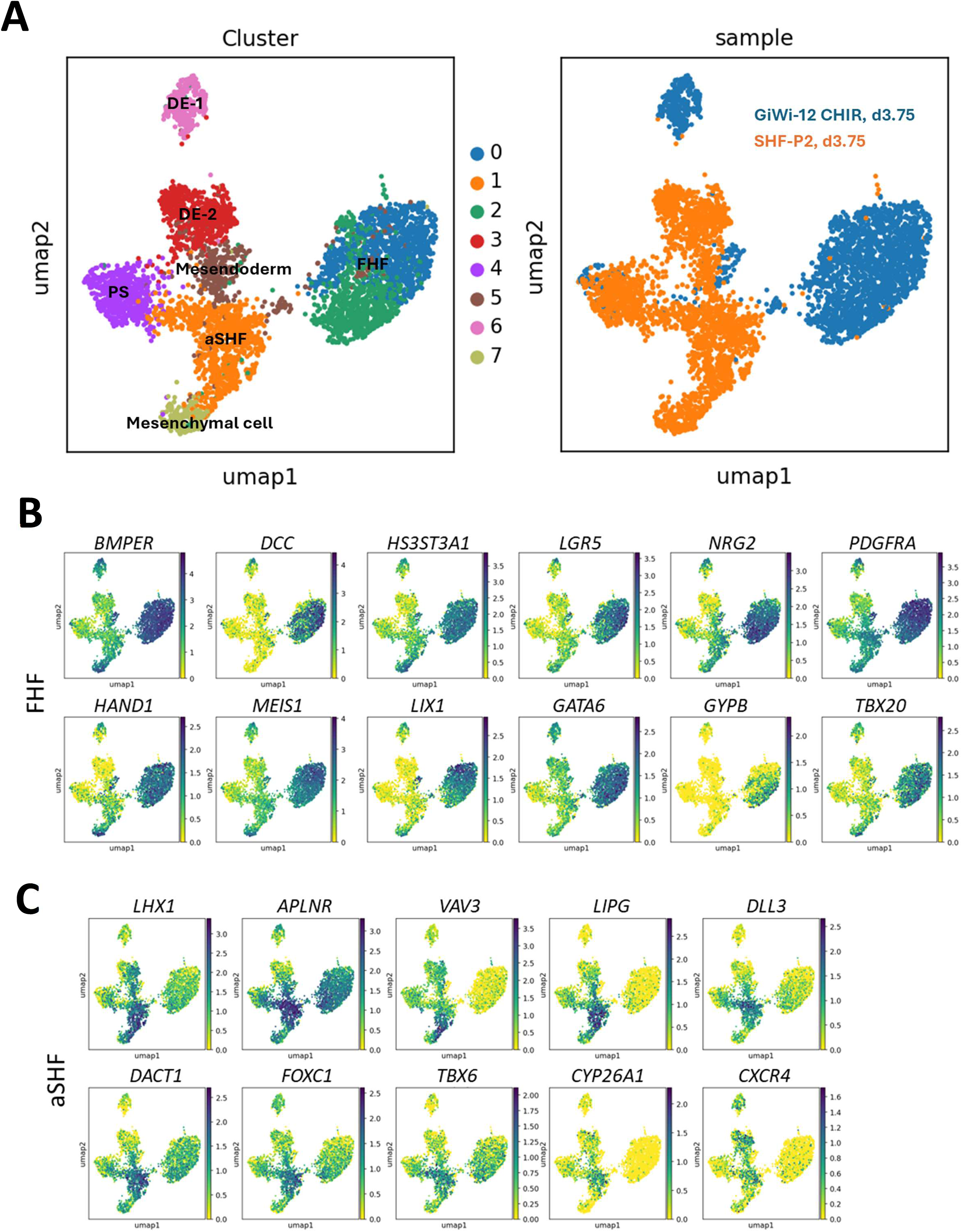
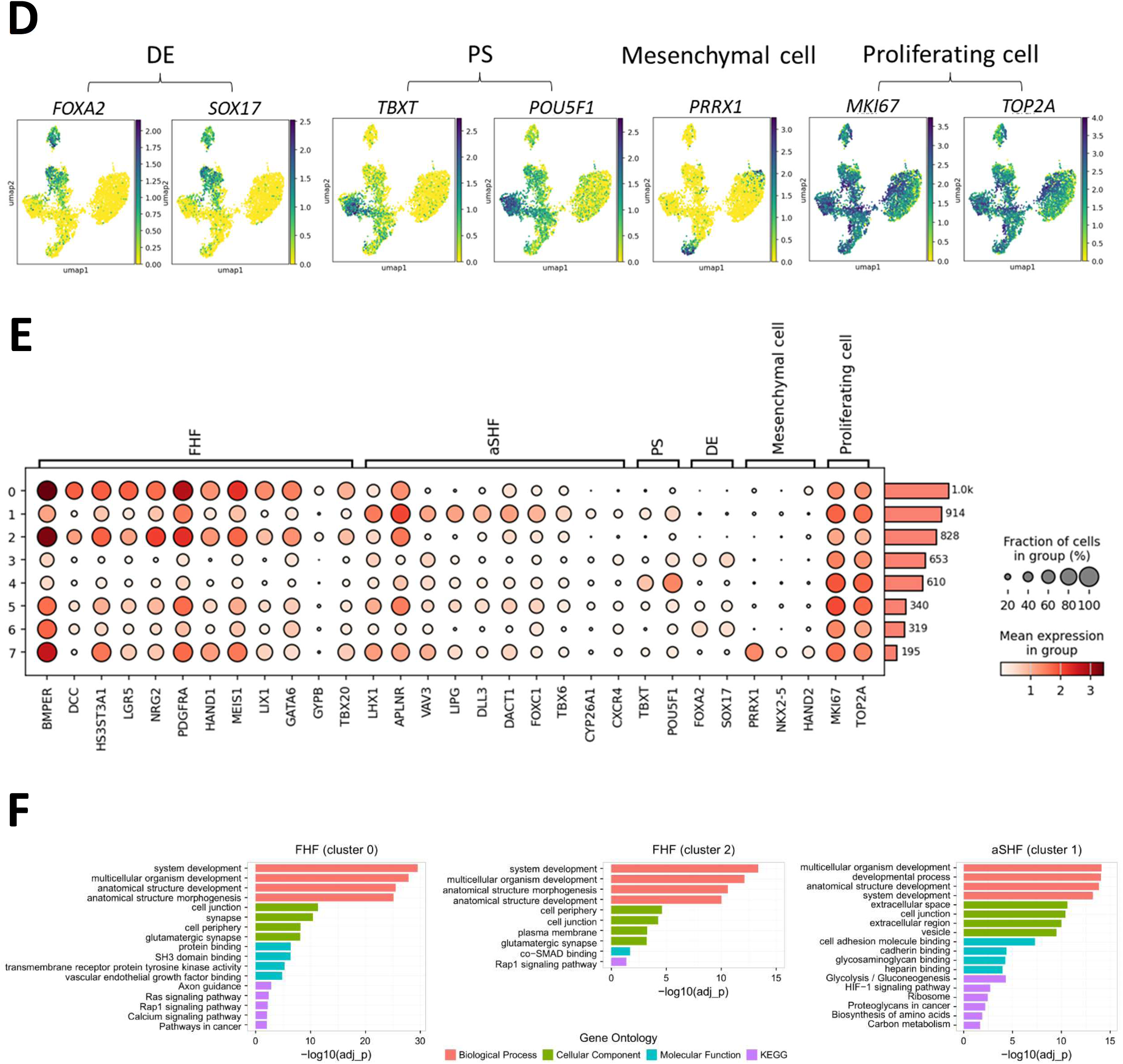
Single-cell transcriptomic analysis of day 3.75 progenitors generated from the SHF-P2, SHF-P3 and GiWi protocols. (A) UMAP of a total of 4888 cells of the day 3.75 progenitors differentiated from GiWi-12CHIR and SHF-P2 protocols. Left, UMAP of cluster view with annotated cell type; Right, UMAP of sample view. Clustering resolution is 0.5. (B) Embedding plots of representative significantly upregulated DEGs of FHF progenitors in log_2_FC. (C) Embedding plots of representative significantly upregulated DEGs of anterior SHF (aSHF) progenitors in log_2_FC. (D) Embedding plots of representative marker genes expression of definitive endoderm (DE), primitive streak (PS), mesenchymal cell and proliferating cell in log_2_FC. (E) Dot plot of the representative marker genes expression used to annotate the cell types. Bars represent cell number for each cluster. Cells with low gene expression of 0-0.5 were removed. (F) GO term analysis of the day 3.75 progenitors derived from GiWi and SHF-P2 protocols. The top 4 most significantly upregulated terms in Biological Process (BP), Cellular Component (CC), Molecular Function (MF) and KEGG Pathway are presented as -log10(adj_P_val) in bar graphs.

The GO term analysis shows the most significantly upregulated Biological Process (BP) including system development, multicellular organism development, anatomical structure development shared in both FHF and aSHF progenitors, while the most significantly upregulated Cellular Component (CC) in FHF is synapse and cell periphery, in contrast to aSHF is extracellular space and cell junction (**Figure 4F and Supplementary File 1**). The most significantly upregulated KEGG pathways in FHF progenitors are Axon guidance and Ras and Rap1 signaling pathway, in contrast to aSHF progenitors being Glycolysis / Gluconeogenesis, HIF-1 signaling and Ribosome pathway, suggesting fast growing and migratory SHF progenitors in contrast to the FHF progenitors at this timepoint (**Figure 4F and Supplementary File 1**).

Interestingly, HIF-1 signaling was identified specifically and significantly upregulated in the aSHF progenitors but not in FHF progenitors, consistent with FGF2 signaling interacting with hypoxia-induced signaling during the SHF development ^61,62^.

### Single-cell transcriptomics of the CM cell types

We next analyzed the scRNA-seq data of the day 20 differentiated cells from the SHF-P2, SHF-P3 and GiWi protocols. Since marker genes for the chamber-specific CMs are not well defined, we aimed to find the marker genes specifically expressed in lvCM, rvCM, aCM and SAN. A high-resolution transcriptomic analysis was performed and identified 18 clusters that represent the major CM cell types and small cell populations of non-CM cell types from each protocol based on marker gene expression, differential gene expression and GO term analysis (**Figure 5A, B**). The CMs differentiated from the GiWi protocol were identified as a major cell population of lvCM including immature lvCM (cluster 0 and 16), lvCM (cluster 7) and proliferating lvCM (cluster 8), consistent with the analysis of the full dataset which includes the day 3.75 progenitors and day 20 cells (**Figure 5A**). The analysis of day 20 samples also identified a small population of aCM derived from the GiWi protocol (cluster 6) by expression of *VWDE* (**Figure 5A,B**). We found *FGF12* was specifically and highly expressed in clusters 0, 7, 8 and 16 that were identified as lvCM, which is consistent with a previous single-cell transcriptome study of human fetal heart tissue showing *FGF12* is a specific marker gene for human LV ^63^ (**Figure 5B)**. The other DEGs of *MYL3*, *MYL4*, *LAMA2*, *LAMA4*, *TTN*, *CACNA1C* and *RYR2* highly expressed in all the lvCMs, while the mitochondrial genes of *MT-RNR1*, *MT-RNR2*, *MT-ND5* and *MT-ND4* are differentially expressed in cluster 7 (lvCM) and *MKI67* and *TOP2A* are differentially expressed in cluster 8 that distinguishes the lvCM, proliferating lvCM from the immature lvCM, respectively (**Supplementary Figure S9**). Only a few lnc RNAs including *ENSG00000289426*, *ENSG00000289578*, and *ENSG00000286339* and annotated genes of *CLEC2A*, *ANKRD4*5 and *SPTLC3* distinguish cluster 16 from 0 (**Supplementary Figure S10**). The CMs derived from the SHF-P2 protocol were identified as one major cell population of rvCM (cluster 2) **(Figure 5A,B)**. The *ABRA* gene was exclusively expressed in rvCM together with the DEGs of *A2M* and *MEF2C* could be specific marker genes to identify rvCM, given that *MEF2C* is a key transcription factor in aSHF development and formation of RV and OFT during mouse heart development ^59,64^ (**Figure 5B**). The CMs derived from the SHF-P3 protocol were identified as two major CM populations of aCM (cluster 1) and SAN (cluster 11) by DEGs of *VWDE* for aCM and *SHOX2* for SAN (**Figure 5A,B)**. Interestingly, we found the gene *VWDE* was specifically and greatly expressed in aCM and SAN, in contrast to *NR2F2* which was identified as a marker gene for aCM previously ^51^ but showed low expression in the aCM derived from both the SHF-P3 and GiWi protocols (**Figure 5B**). Moreover, *SHOX2* was exclusively expressed in SAN that distinguishes SAN from aCM (**Figure 5B**).

**Figure 5.**
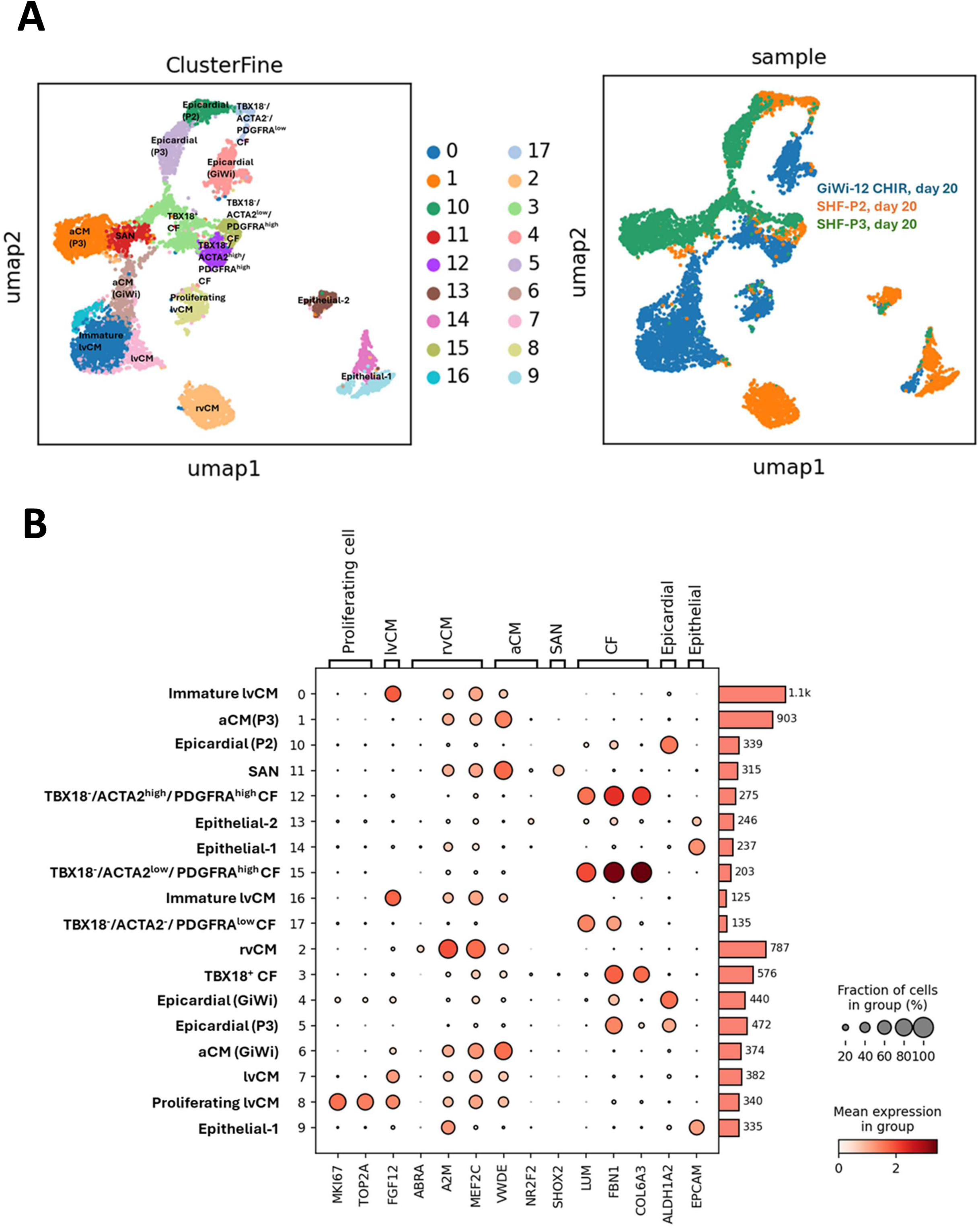
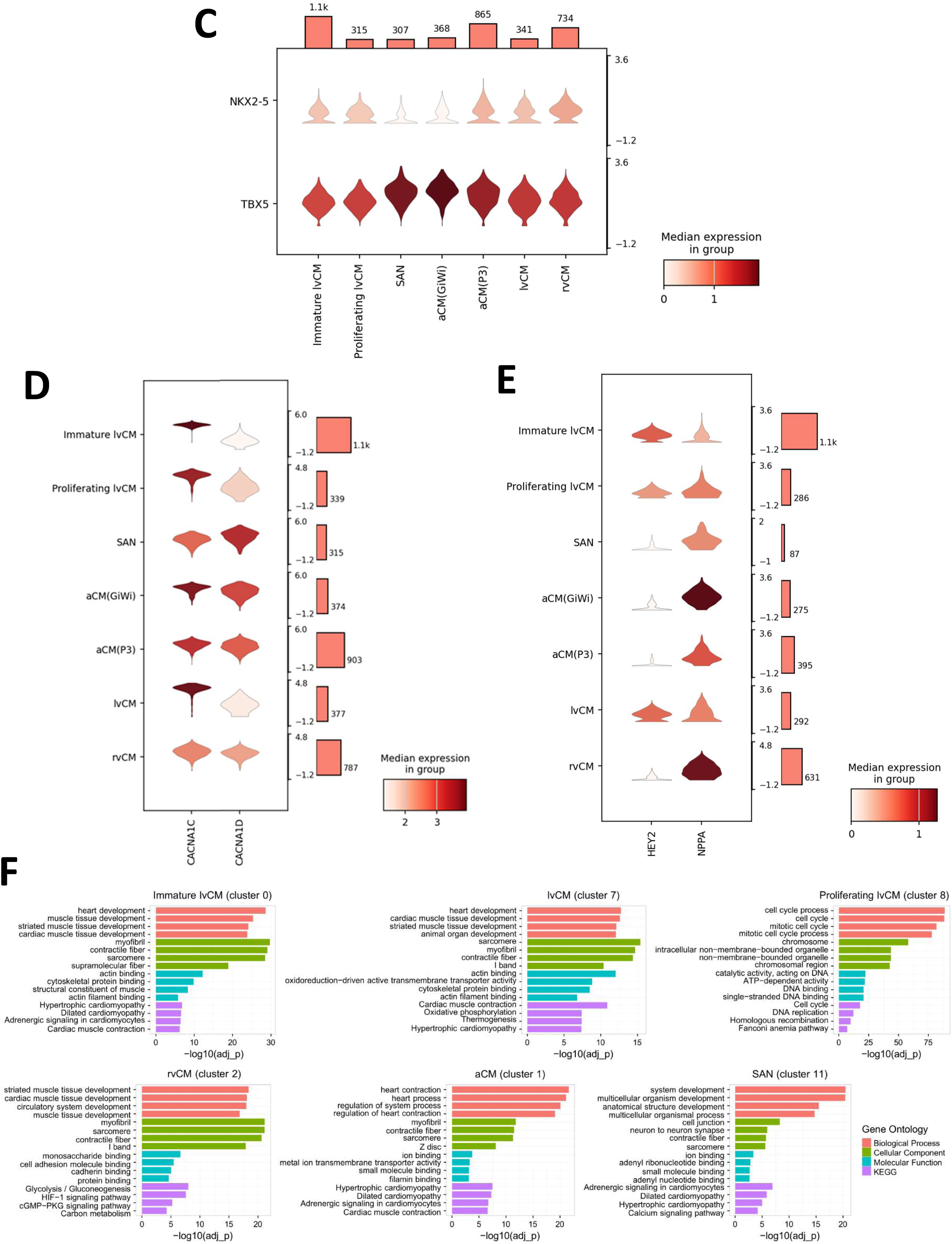
Single-cell transcriptomic analysis of day 20 differentiated cells from the SHF-P2, SHF-P3 and GiWi protocols. (A) UMAP of a total of 7597 cells of the day 20 differentiated cells from GiWi-12CHIR, SHF-P2 and SHF-P3 protocols. Left, UMAP of cluster view with annotated cell type; Right, UMAP of sample view. Clustering resolution is 0.9. (B) Dot plot of the representative marker genes expression used to annotate the cell types. Bars represent cell number for each cluster. Cells with low gene expression of 0-1 were removed. (C) Violin plot of the differential expression of *TBX5* and *NKX2-5* in the chamber-specific hPSC-CMs. Bars represent cell number for each cluster. Y-axis of violin plot represents log_2_(TPM+1) values for gene expression. (D) Violin plot of the differential expression of *CACNA1C* and *CACNA1D* in the chamber-specific hPSC-CMs. Bars represent cell number for each cluster. Y-axis of violin plot represents log_2_(TPM+1) values for gene expression. (E) Violin plot of the differential expression of *HEY2* and *NPPA* in the chamber-specific hPSC-CMs. Bars represent cell number for each cluster. Y-axis of violin plot represents log_2_(TPM+1) values for gene expression. (F) GO term analysis of the day 20 differentiated hPSC-CMs derived from GiWi, SHF-P2 and SHF-P3 protocols. The top 4 most significantly upregulated terms in Biological Process (BP), Cellular Component (CC), Molecular Function (MF) and KEGG Pathway are presented as - log10(adj_P_val) in bar graphs.

*NKX2-5* and *TBX5* are two critical transcription factors that are differentially expressed in lvCM, rvCM, aCM and SAN, so we also analyzed their expression in the day 20 CMs from the 3 protocols (**Figure 5C**). Overall, *TBX5* expressed higher relative to *NKX2-5* in all types of CMs. However, *NKX2-5* expression was relatively higher in rvCM than lvCM, proliferating lvCM and immature lvCM, consistent with the expression pattern of these two genes in FHF- and SHF-lineages (**Figure 5C**). *TBX5* expression was extremely high in SAN and aCM, and there was almost no expression of *NKX2-5* in SAN as shown in previous studies ^65^, and low expression in aCM (**Figure 5C**).

Two isoforms of the voltage-gated calcium channel alpha1 subunit genes, *CACNA1C* and *CACNA1D* were also differentially expressed in chamber-specific hPSC-CMs. The immature lvCM, proliferating lvCM and lvCM had extremely high expression of *CACNA1C* and very low expression of *CACNA1D*, in contrast to SAN which had significantly higher expression of *CACNA1D* than *CACNA1C*; rvCM and aCM had comparable expression of both isoforms (**Figure 5D**). This is consistent with the published chamber-specific expression of these channel isoforms ^66^.

*Hey2* and *Nppa* were identified as specific molecular markers to label compact and trabecular ventricular myocardium, respectively ^54,58,59^, so we analyzed their expression in our chamber-specific hPSC-CMs differentiated from the GiWi, SHF-P2 and SHF-P3 protocols (**Figure 5E and Supplementary Figure S11**). Interestingly, the day 20 differentiated rvCMs from the SHF-P2 protocol highly express *NPPA* but low expression of *HEY2*, in contrast to the day 20 differentiated GiWi-immature lvCMs that have relatively higher expression of *HEY2* than *NPPA* while comparable expression in GiWi-lvCM and GiWi-proliferating lvCM (**Figure 5E and Supplementary Figure S11**). The differential expression of these two ventricular myocardium genes may suggest a different subpopulation of compact and trabecular CMs present in the GiWi-immature lvCMs, lvCMs and proliferating lvCMs, and in the SHF-P2-rvCMs.

We also explored cell-surface markers for isolation of chamber-specific hPSC-CMs. SIRPA (CD172a) was previously identified as a surface marker that highly expressed in cardiac precursors and cardiomyocytes during hPSC-CM differentiation, as well as in human fetal LV and LA (left atrial) and human adult heart ^67^. However, we found that the SHF-P2-rvCMs and SHF-P3-aCMs had significantly lower expression of SIRPA than the GiWi-lvCMs and GiWi-aCMs as shown in our scRNA-seq data (**Supplementary Figure S12A**). In contrast, the surface marker CD109 expressed more in the SHF-P2-rvCMs, and there was almost no CD109 expression in the GiWi-lvCMs or GiWi-aCMs (**Supplementary Figure S12A**). We also confirmed the SIRPA and CD109 expression by flow cytometry in the Nutlin-3a purified ^68^ DF19-9-11T iPSC-CMs differentiated from the SHF-P2 and GiWi protocols (**Supplementary Figure S12B**). These results suggest that SIRPA and CD109 are differentially expressed in the FHF- and SHF-derived hPSC-CMs.

The GO term analysis was also performed on the CM populations derived from the GiWi, SHF-P2 and SHF-P3 protocols. All the CM populations share the most significantly upregulated BP terms including striated muscle tissue development and cardiac muscle tissue development, however, one distinct BP in rvCM (cluster 2) is circulatory system development that is not identified in lvCM (cluster 0 and 7), suggesting vasculature development alongside the CM differentiation from the SHF progenitors (**Figure 5E and Supplementary File 2**). The immature lvCM (cluster 0), lvCM (cluster 7) and rvCM (cluster 2) have similar most significantly upregulated CC including myofibril, contractile fiber and sarcomere, but some distinct MF in which the rvCM has monosaccharide binding, cell adhesion molecular binding and cadherin binding but lvCM has actin binding, cytoskeletal protein binding and structural constituent of muscle (**Figure 5E and Supplementary File 2**), suggesting the rvCM exhibit more mature cell-to-cell coupling than the lvCMs. The most significantly upregulated KEGG pathways that are shared in immature lvCM (cluster 0) and lvCM (cluster 7) include cardiac muscle contraction and hypertrophic cardiomyopathy (**Figure 5E and Supplementary File 2**). However, the most significantly upregulated KEGG pathways in rvCM are distinct from the lvCM and consistent with the aSHF progenitor-upregulated KEGG pathways including glycolysis / gluconeogenesis, HIF-1 signaling cGMP-PKG signaling and carbon metabolism (**Figure 5E and Supplementary File 2**). Furthermore, the most significantly upregulated BP, CC, MF and KEGG pathway in cluster 8 which is identified as proliferating lvCM by marker genes (**Figure 5A**) are related to cell cycle, DNA binding and DNA replication, distinct from other CM populations (**Figure 5E and Supplementary File 2**). The most significantly upregulated BP in aCM (cluster 1) include heart contraction, heart process and regulation of system process, while the enriched BP in SAN (cluster 11) include system development, multicellular organism development and anatomical structure development (**Figure 5E and Supplementary File 2**); And the aCM (cluster 1) and SAN (cluster 11) share similar most significantly upregulated CC, MF and KEGG pathway, although the aCM shows more specialized for heart contraction in BP and KEGG while SAN shows more neuron to neuron synapse in CC, adenyl ribonucleotide binding in MF and calcium signaling in KEGG pathway (**Figure 5E and Supplementary File 2**). In summary, GO term analysis identified distinguishing features between the CM clusters, for example, with the prominence of glycolysis / gluconeogenesis and HIF-1 signaling in the rvCM cluster relative to the other clusters.

### Single-cell transcriptomics of non-CM cell types

Transcriptomic analysis of the day 20 differentiated cells also revealed some non-CM cell types including epicardial, epithelial cell and cardiac fibroblast (CF) (**Figure 5A,B**). Clusters from GiWi (cluster 4), SHF-P3 (cluster 5) and SHF-P2 (cluster 10) protocols were identified as epicardial cells which specifically expressed the marker gene *ALDH1A2* (**Figure 5B**). In addition, *TBX18* and *WT1*, two typical marker genes for epicardial cells were also highly expressed in the three populations (**Supplementary Figure S13**). Interestingly, *UPK3B* has been identified as another marker gene for epicardial cells in human fetal heart ^63,69^, but was not expressed abundantly in the hPSC-derived epicardial cells (**Supplementary Figure S13**).

Multiple CF populations were identified by the DEGs including *LUM*, *FBN1*, *COL6A3*, *COL3A1*, *COL1A1*, *TBX18*, *TCF21*, *PDGFRA* and *POSTN* (**Supplementary Figure S14**). We found *LUM*, *FBN1* and *COL6A3* are the most specifically expressed genes in the CFs and can distinguish the CF populations from epicardial cells (**Figure 5B**). Two small cell populations (Epithelial −1; Epithelial-2) mainly derived from SHF-P2 protocol were identified as epithelial cells by the specific marker gene *EPCAM* (**Figure 5B**).

### Comparison of gene expression profiles of the chamber-specific hPSC-CMs differentiated from our protocols with other protocols as well as human fetal heart tissue

To explore how the gene expression profiles of the hPSC-lvCM, rvCM and aCM generated from GiWi, SHF-P2 and SHF-P3 protocols compare to the counterparts generated from other differentiation protocols ^43^, as well as human fetal heart of 5-25 weeks of gestation ^63^, we performed the transcriptomic analysis across all three scRNA-seq datasets.

The comparison of the DEGs in the day 20 differentiated hPSC-CMs from our dataset and Yang’s dataset ^43^ was performed by comparing the specific clusters for the chamber-specific CMs from each dataset. The comparison of the immature lvCM, lvCM and proliferating lvCM differentiated from our GiWi protocol (**Figure 5A**) with the LVLCM and Prol_LVLCM differentiated from FHF Day20 cells in Yang’s dataset ^43^ revealed 672, 491 and 458 shared DEGs, respectively (**Figure 6A and Supplementary Figure S15, File 3**). We further analyzed the gene expression of the shared DEGs as shown in each Venn Diagram and show the top 50 most significantly upregulated DEGs selected by our dataset (**Figure 6B**) or by Yang’s dataset (**Supplementary Figure S16**). Further comparison of the top 50 significantly upregulated DEGs selected by either GiWi-immature lvCM or Yang’s LVLCM revealed that *NAV1* and *FBN2* are among the top 50 most significantly upregulated DEGs in both datasets (**Supplementary Table S2 and File 3**); The comparison of the GiWi-lvCM with Yang’s LVLCM showed that the mitochondria genes of *MT-CYB* and *MT-ATP6*, and *NAV1* are among the top 50 most significantly upregulated DEGs in both datasets (**Supplementary Table S2 and File 3**); The comparison of the GiWi-proliferating lvCM with Yang’s Prol_LVLCM identified that *MKI67*, *TOP2A* and *CENPF* which are the marker genes for cell proliferation, DNA replication and mitosis are among the top 50 most significantly upregulated DEGs in both datasets (**Supplementary Table S2 and File 3**).

**Figure 6.**
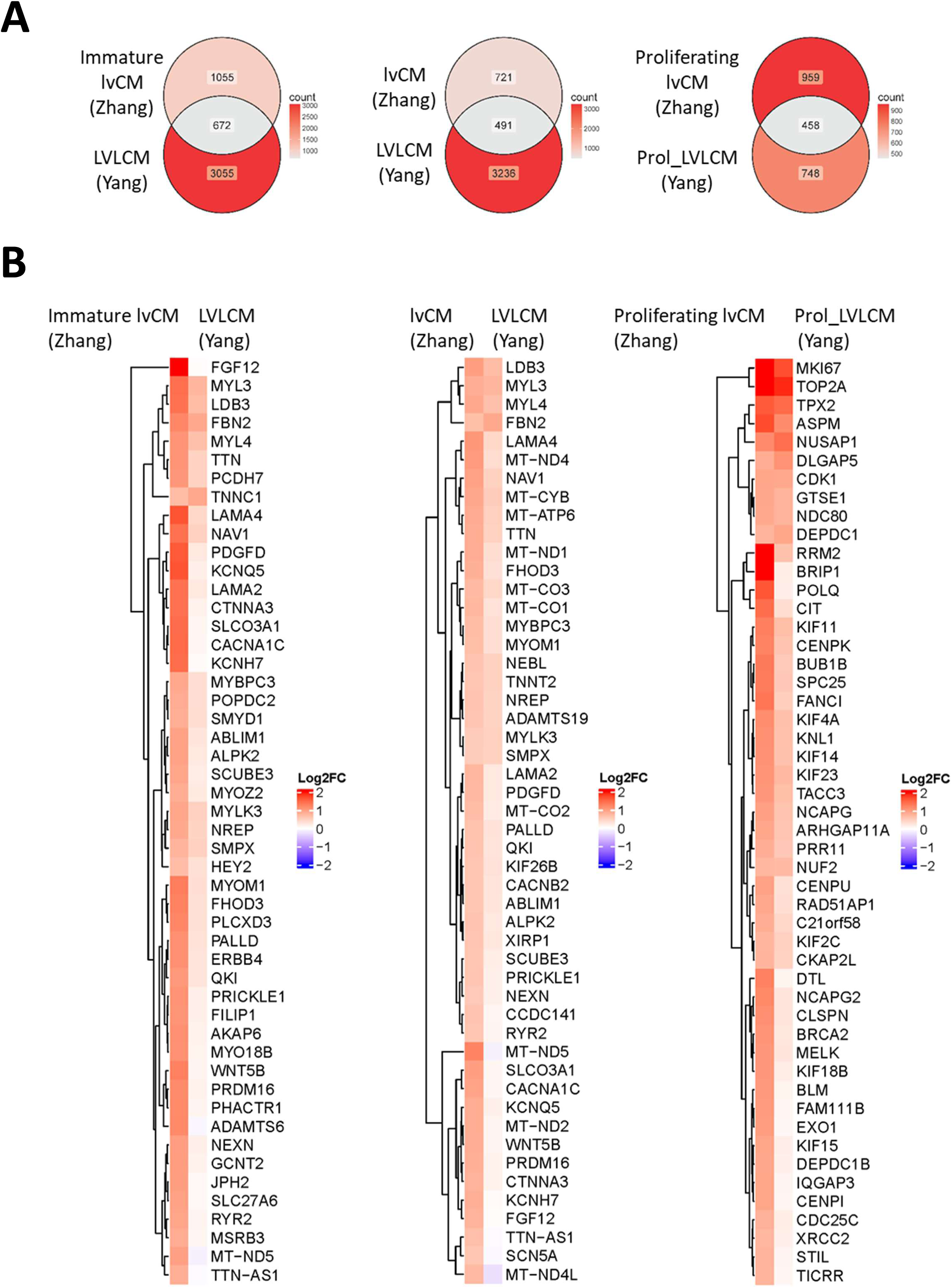
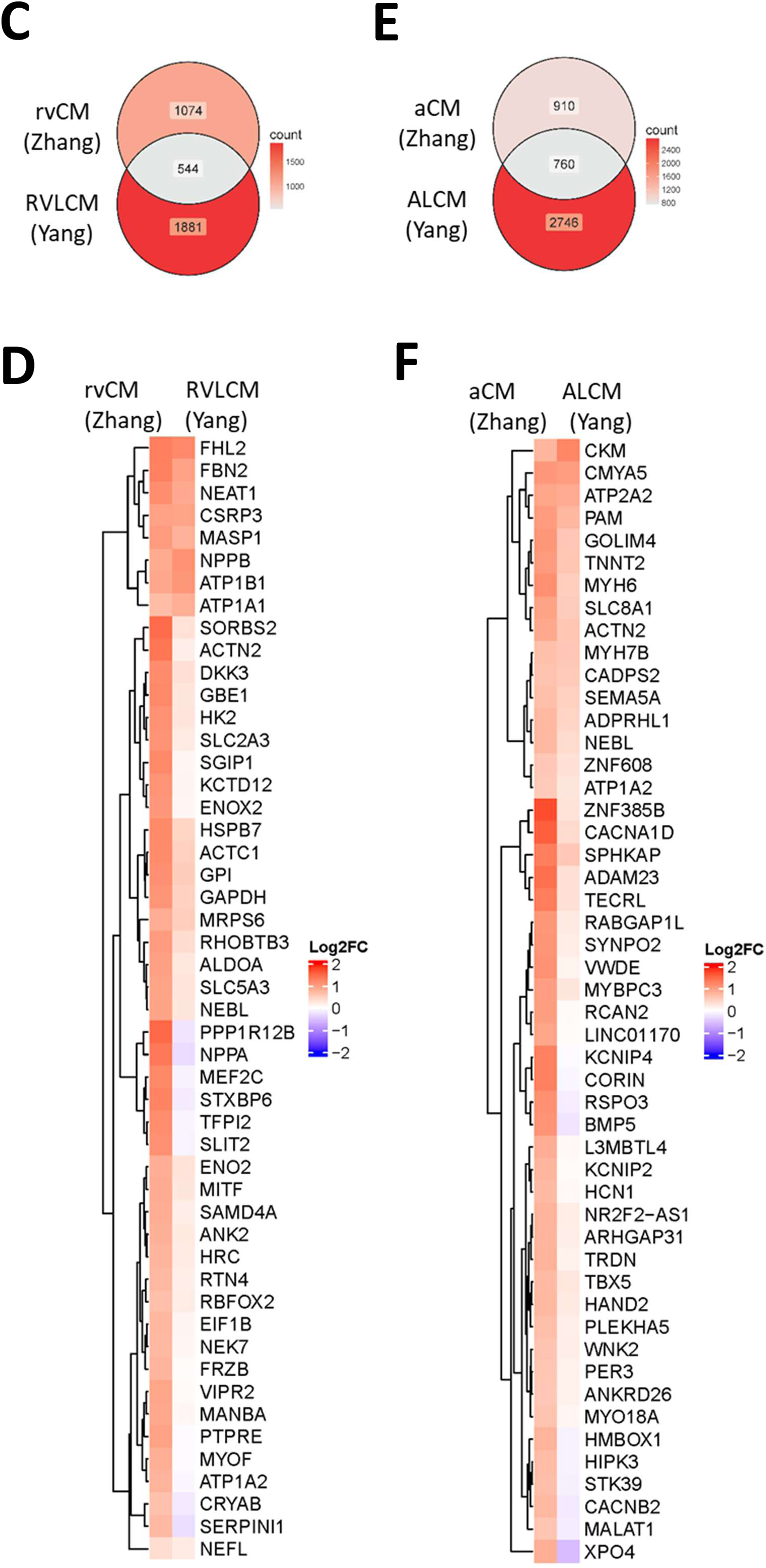
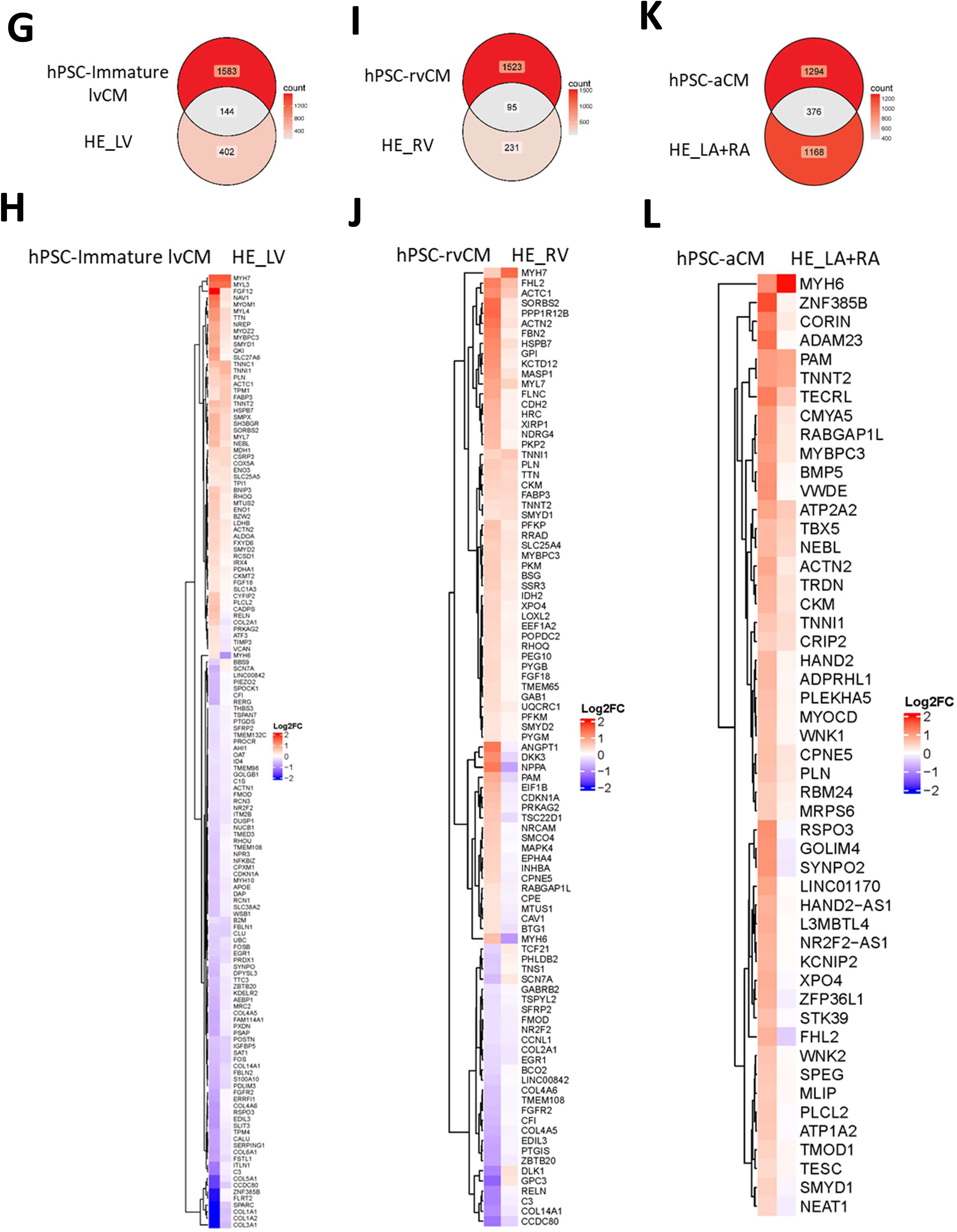
Comparison of gene expression profiles of the chamber-specific hPSC-CMs generated from GiWi, SHF-P2 and SHF-P3 protocols with other protocol generated hPSC-CMs and human fetal heart tissue. (A) Venn diagrams of the shared DEGs comparing the immature-lvCM, lvCM and proliferating lvCM from our dataset with the LVLCM and Prol_LVLCM from Yang’s dataset ^43^. (B) Heatmap with unsupervised hierarchical clustering of gene expression of the top 50 significantly upregulated shared DEGs ordered by the significance from our dataset, when comparing the immature-lvCM, lvCM and proliferating lvCM from our dataset with the LVLCM and Prol_LVLCM from Yang’s dataset ^43^. (C) Venn diagram of the shared DEGs comparing the rvCM from our dataset with the RVLCM from Yang’s dataset ^43^. (D) Heatmap with unsupervised hierarchical clustering of gene expression of the top 50 significantly upregulated shared DEGs ordered by the significance from our dataset, when comparing the rvCM from our dataset with the RVLCM from Yang’s dataset ^43^. (E) Venn diagram of the shared DEGs comparing the aCM from our dataset with the ALCM from Yang’s dataset ^43^. (F) Heatmap with unsupervised hierarchical clustering of gene expression of the top 50 significantly upregulated shared DEGs ordered by the significance from our dataset, when comparing the aCM from our dataset with the ALCM from Yang’s dataset ^43^. (G) Venn diagrams of the shared DEGs comparing the immature-lvCM from our dataset with the human fetal HE_LV from Cui’s dataset ^63^. (H) Heatmap with unsupervised hierarchical clustering of gene expression of the shared DEGs, when comparing the immature-lvCM from our dataset with the HE_LV from Cui’s dataset ^63^. (I) Venn diagrams of the shared DEGs comparing the rvCM from our dataset with the human fetal HE_RV from Cui’s dataset ^63^. (J) Heatmap with unsupervised hierarchical clustering of gene expression of the shared DEGs, when comparing the rvCM from our dataset with the HE_RV from Cui’s dataset ^63^. (K) Venn diagrams of the shared DEGs comparing the aCM from our dataset with the human fetal HE_LA+RA (left and right atria) from Cui’s dataset ^63^. (L) Heatmap with unsupervised hierarchical clustering of gene expression of the shared DEGs, when comparing the aCM from our dataset with the HE_LA+RA from Cui’s dataset ^63^.

The comparison of the DEGs in the day 20 differentiated rvCM from our SHF-P2 protocol with the RVLCM differentiated from the aSHF Day 20 cells in Yang’s dataset ^43^ revealed 544 shared DEGs (**Figure 6C and Supplementary Figure S15, File 3**). We further analyzed the gene expression of the shared DEGs as shown in the Venn Diagram (**Figure 6C**) and show the top 50 most significantly upregulated DEGs selected by our dataset (**Figure 6D**) or by Yang’s dataset (**Supplementary Figure S17**). Comparison of the top 50 significantly upregulated DEGs in either SHF-P2-rvCM or Yang’s RVLCM showed that *FBN2, MASP1* and *FHL2* are among the top 50 most significantly upregulated DEGs in both datasets (**Supplementary Table S2 and File 3**).

The comparison of the DEGs in the day 20 differentiated aCM from our SHF-P3 protocol with the ALCM differentiated from the pSHF Day 20 cells in Yang’s dataset ^43^ revealed 760 shared DEGs (**Figure 6E and Supplementary Figure S15, File 3,**). We further analyzed the gene expression of the shared DEGs as shown in the Venn Diagram (**Figure 6E**) and show the top 50 most significantly upregulated DEGs selected by our dataset (**Figure 6F**) or by Yang’s dataset (**Supplementary Figure S18**). The comparison of the top 50 significantly upregulated DEGs in either SHF-P3-aCM or Yang’s ALCM showed that *CMYA5*, *ATP2A2, SPHKAP*, *SLC8A1* and *PAM* are among the top 50 most significantly upregulated DEGs in both datasets (**Supplementary Table S2 and File 3**).

We also analyzed the shared DEGs from our and Yang’s datasets to identify the different gene expression pattern, i.e. the shared DEGs that are upregulated in our dataset but downregulated in Yang’s dataset. The comparison of the immature lvCM and lvCM from our GiWi protocol with the LVLCM from Yang et al. ^43^ showed the mitochondrial genes including *MT-ATP8* was upregulated in the GiWi-lvCM, along with *MT-ND3*, *MT-ND4L*, *MT-ND5* and *MT-ND6* were upregulated in both the GiWi-lvCM and immature lvCM, but were all downregulated in the LVLCM from the Yang’s dataset (**Supplementary File 4**). Given that these mitochondrial genes encode the core subunits of the electron transport chain (ETC) that are essential for mitochondrial machinery and shift the cell’s reliance from glycolysis to oxidative phosphorylation, it may indicate the GiWi-lvCM is a relatively more mature hPSC-CM population than the LVLCM from Yang et al. ^43^. The different gene expression patterns, when comparing our SHF-P2-rvCM with the RVLCM from Yang et al. ^43^, showed the gene *PPP1R12B* that modulates muscle contraction, *NPPA* which is a specific molecular marker for ventricular trabecular CMs during heart development, *ANGPT1* for angiogenesis, vascular maturation and endothelial cell stability, *ADAMTS19* and *TFPI2* involved in extracellular matrix remodeling, *SLIT2* as a tumor suppressor gene, as well as the SHF marker genes of *MEF2C* and *HAND2*, are significantly upregulated in our SHF-P2-rvCM but downregulated in the RVLCM in Yang’s dataset (**Figure 6D and Supplementary File 4**). These different gene expression patterns suggest a more specific SHF-derived, trabecular rvCMs differentiated from our SHF-P2 protocol than the RVLCMs derived from Yang et al^43^. The most different gene expression when comparing our SHF-P3-aCM with the ALCM from Yang et al. ^43^, include the gene *CORIN*, which encodes a membrane-bound serine protease that converts pro-atrial natriuretic peptide (pro-ANP) into active ANP, and *KCNIP4*, *BMP5* and *RSPO3*, all of the genes are upregulated DEGs in our SHF-P3-aCM but downregulated in Yang’s ALCM (**Figure 6F and Supplementary File 4**).

In summary, the comparison of the heart field- and chamber-specific hPSC-CMs derived from our GiWi and SHF protocols with Yang et al. ^43^ demonstrated the most similar CM population being the proliferating lvCM that have *MKI67* and *TOP2A* as the top significantly upregulated DEGs in both clusters. The GiWi-lvCM is relatively more mature than the GiWi-immature lvCM and LVLCM from Yang’s dataset as shown in the upregulation of core mitochondrial genes (**Supplementary Table S2**). The marker genes identified in our dataset for the chamber-specific hPSC-CMs, including *FGF12* for lvCM, *ABRA*, *A2M* and *MEF2C* for rvCM, and *VWDE* for aCM (**Figure 5B**) were not found on the top of most significantly upregulated DEGs in Yang et al ^43^ (**Supplementary Table S2**). Furthermore, *NPPA* that highly express in trabecular CM was significantly upregulated in the rvCM derived from our SHF-P2 protocol but downregulated in RVLCM from Yang et al^43^.

We next performed similar analysis to compare our dataset with Cui et al. dataset of the human fetal heart tissue ^63^. The comparison of the DEGs of the day 20 differentiated GiWi-immature lvCM, lvCM and proliferating lvCM with the LV cells (HE5W_LV - HE25W_LV) from Cui’s dataset ^63^ revealed 144, 111 and 87 shared DEGs, respectively (**Figure 6G and Supplementary Figure S19, File 5**). We further analyzed the gene expression of all the shared DEGs and show the heatmap for the expression pattern comparing the two datasets (**Figure 6H and Supplementary Figure S19**). We found the GiWi-immature lvCM and HE_LV have quite similar expression pattern and share the important marker genes for LV, including *FGF12*, *MYL3*, *NAV1* and *MYOM1* (**Figure 6H** and **Supplementary Table S2, File 5**). *FGF12* was identified as human specific LV marker gene in Cui et al^63^ and is also significantly upregulated in the GiWi-immature lvCM. *MYL3* encodes essential myosin light chain protein that regulates muscle contraction, functioning primarily in ventricles and slow skeletal muscle, in contrast to *MYL4* acting predominantly in atria and embryonic tissue. *NAV1* is among the top 50 most significantly upregulated DEGs in all 3 datasets including Yang’s dataset (**Supplementary Table S2**). *MYOM1* encodes myomesin-1, a crucial structural protein in cross-striated cardiac and skeletal muscles. The most differently expressed genes when comparing our data with Cui’s data include *MYH6*, *VCAN* and *TIMP3* which are significantly upregulated in the GiWi-immature lvCM but downregulated in the HE_LV ^60^, suggesting a less mature phenotype in the GiWi-immature lvCMs (**Figure 6H**). The GiWi-lvCM and proliferating lvCM have similar gene expression profile as the GiWi-immature lvCM when compared to the HE_LV (**Supplementary Figure S19 and File 5**).

We next compared the gene expression profile of the rvCM differentiated from our SHF-P2 protocol with the HE_RV from Cui’s dataset ^63^. The comparison of the DEGs in each dataset revealed 95 shared DEGs (**Figure 6I and Supplementary File 5**). Further analysis of the shared DEGs showed the SHF-P2-rvCM has similar expression pattern as the HE_RV, including *PPP1R12B*, *SORBS2*, *HSPB7*, *GPI*, *MASP1*, *FHL2* and *ACTC1* are among the top 50 most significantly upregulated DEGs in both datasets (**Figure 6J and Supplementary Table S2, File 5**). Interestingly, *ANGPT1*, *DKK3*, *NPPA*, *PAM* and *MYH6* which are significantly upregulated DEGs in the SHF-P2-rvCM but downregulated in the HE_RV, with functions of these genes for vessel maturation and stabilization (*ANGPT1)*, tumor suppressor (*DKK3*), trabecular myocardium expression (*NPPA*) and cardiac muscle contraction (*MYH6*). *TCF21*, *PHLDB2*, *TNS1*, *SCN7A*, *DLK* and *GPC3* are significantly downregulated DEGs in the SHF-P2-rvCM but upregulated in HE_RV (**Figure 6J**), likely due to the mixture of non-CMs in the HE_RV tissue samples.

The comparison of the DEGs of the day 20 differentiated aCM from our SHF-P3 protocol (**Figure 5A**) with the HE_LA+RA from Cui’s dataset ^63^ revealed 376 shared DEGs (**Figure 6K**). Analysis of the gene expression of the shared DEGs showed similar expression pattern between the SHF-P3-aCM and the HE_LA+RA (**Supplementary Table S2, File 5, 6**). Further analysis of the gene expression of the shared DEGs show the top 50 most significantly upregulated DEGs selected by our dataset (**Figure 6L**) or by Cui’s dataset (**Supplementary Figure S20**).

*ZNF385B*, *TECRL*, *VWDE*, *CORIN*, *CMYA5*, *ATP2A2*, *TRDN*, *PAM*, *MYH6* and *TBX5* are among the top 50 most significantly upregulated DEGs in both datasets (**Supplementary Table S2, File 5**). *VWDE* (von Willebrand factor D and EGF domains) was identified as an aCM marker gene in our scRNA-seq data (**Figure 5B**) was also a significantly upregulated DEG in HE_LA+RA (**Supplementary Table S2, File 5**). In addition, *ATP2A2*, *CMYA5* and *PAM* were also among the top 50 most significantly upregulated DEGs in Yang’s dataset (**Supplementary Table S2**). The differently expressed genes, however, include *MYH7*, *FHL2*, *SYNPO2* and *GOLIM4* that are significantly upregulated in SHF-P3-aCM but downregulated in HE_LA+RA (**Supplementary File 5,6**). The epithelial marker genes of *KRT19*, *KRT18*, *KRT8* and *ITLN1*, along with *ANKRD1* and *CALD1* were upregulated in HE_LA+RA but downregulated in SHF-P3-aCM (**Supplementary File 5, 6**), again likely due to the non-cardiomyocytes in the HE_LA+RA tissue samples.

In summary, the comparison of the chamber-specific hPSC-CMs derived from our GiWi, SHF-P2 and P3 protocols with the human fetal heart tissue (human embryos of 5-25 weeks of gestation)^63^ revealed overlapping expression of the marker genes and DEGs for chamber-specificity. *FGF12* which was found as a human-specific LV-CM marker gene in human fetal heart tissue by Cui et al. ^63^ is also identified as the marker gene of GiWi-immature lvCM, lvCM and proliferating lvCM in our dataset (**Figure 5B**). Furthermore, the DEGs of *MYL3*, *NAV1* and *MYOM1* are well aligned in both datasets for lvCM. The significantly upregulated DEGs in our SHF-P2-rvCM including *PPP1R12B*, *SORBS2*, *HSPB7*, *GPI*, *MASP1*, *FHL2* and *ACTC1* are well aligned with the top DEGs in HE_RV. The most obvious difference, however, when comparing the GiWi-immature lvCM, lvCM with HE_LV, and SHF-P2-rvCM with HE_RV is *MYH6* expression, which is significantly upregulated in the ventricular hPSC-CMs but downregulated in the human fetal LV and RV (**Figure 6H, J and Supplementary Figure S19**). The isoforms of *MYH6* and *MYH7* expression are in opposite way in human vs. rodents, in which the switch is from fetal *Myh7* to adult *Myh6* vs. *MYH7* is the predominant isoform in adult human heart. The switch from *MYH6* to *MYH7* is already established by 5 weeks of gestation as shown in Cui’s dataset ^63^. When *MYH6* to *MYH7* switch occurs in human cardiogenesis remains unknown, but newly differentiated hPSC-CMs predominantly express *MYH6* ^70^. In addition, *NPPA* is a significantly upregulated DEG in SHF-P2-rvCM, GiWi-lvCM and proliferating lvCM (not in GiWi-immature lvCM) but downregulated in HE_LV and HE_RV (**Figure 6H, J and Supplementary Figure S19**). These different expression patterns may suggest that GiWi-immature lvCM comprise more compact ventricular CMs similarly as the fetal human LV, while GiWi-lvCM and proliferating lvCM may have comparable or slightly more trabecular than compact ventricular CMs (**Figure 5E and Supplementary Figure S11**), and the SHF-P2-rvCM comprise significant more trabecular than compact ventricular CMs compared to the fetal human RV. *VWDE* which is identified as the marker gene for aCM in our dataset (**Figure 5B**) is also among the top significantly upregulated DEGs in the HE_LA+RA (**Supplementary Table S2, Figure S20**). Furthermore, the significantly upregulated DEGs including *ZNF385B*, *TECRL*, *CORIN*, *CMYA5*, *ATP2A2*, *TRDN*, *PAM*, *MYH6* and *TBX5* are well aligned in both SHF-P3-rvCM and HE_LA+RA (**Supplementary Table S2, Figure S20**).

We conclude that the transcriptomes of the chamber-specific hPSC-CMs derived from our protocols including the GiWi, SHF-P2 and SHF-P3 protocols are aligned with the counterparts derived from Yang et al. ^43^ with the overlapping, significantly upregulated DEGs, particularly the proliferating lvCM in both datasets. However, the comparative studies show our GiWi-immature lvCM and lvCM, SHF-P2-rvCM and SHF-P3-aCM are better aligned with the chamber-specific CMs in human fetal heart^63^ with great overlapping expression of marker genes and DEGs in chamber specificity than the hPSC-CMs derived from Yang et al^43^ (**Supplementary Table S2**).

### Electrophysiology and calcium cycling comparing the chamber-specific hPSC-CM populations

To determine how the functional phenotypes reflect the chamber-specificity established by the scRNA-seq, DF19-9-11T hiPSC-CMs differentiated for 20 days by the GiWi, SHF-P2 and SHF-P3 protocols were dissociated to single cells, plated at low density onto glass coverslips, and recorded 3–7 days later using the perforated-patch current-clamp configuration at 37 °C. Averaged 1 Hz–paced action potential (AP) waveforms showed broadly similar cardiac AP morphologies across the hiPSC-CMs from the three protocols (**Figure 7A**). After correction for the liquid junction potential, maximum diastolic potential was more negative in SHF-P2–rvCMs (−76.0 ± 5.5 mV, n = 32) than in GiWi-lvCM (−72.9 ± 5.1 mV, n = 42; p = 0.045) and SHF-P3–aCM (−73.6 ± 5.9 mV, n = 29) (**Figure 7B**). Action potential duration at 80% repolarization did not differ among the three preparations (**Figure 7C**). Maximum upstroke velocity (dV/dt_max_) was comparable in GiWi-lvCM (199.5 ± 57.8 V/s) and SHF-P2–rvCM (217.2 ± 46.7 V/s; p = 0.45), but dV/dt_max_ was lower in SHF-P3–aCM (162.1 ± 82.0 V/s) than in either ventricular preparation (p = 0.040 vs. lvCM; p = 0.0025 vs. rvCM) (**Figure 7D**). Cell capacitance, an established indicator of cardiomyocyte size,^71–73^ was significantly larger in SHF-P2–rvCM (120.4 ± 53.9 pF) than in GiWi-lvCM (78.4 ± 20.9 pF, p = 0.0002) and SHF-P3–aCM (92.4 ± 43.1 pF, p = 0.038), indicating that the SHF-P2 protocol generates substantially larger cardiomyocytes than the GiWi protocol (**Figure 7E**).

**Figure 7.**
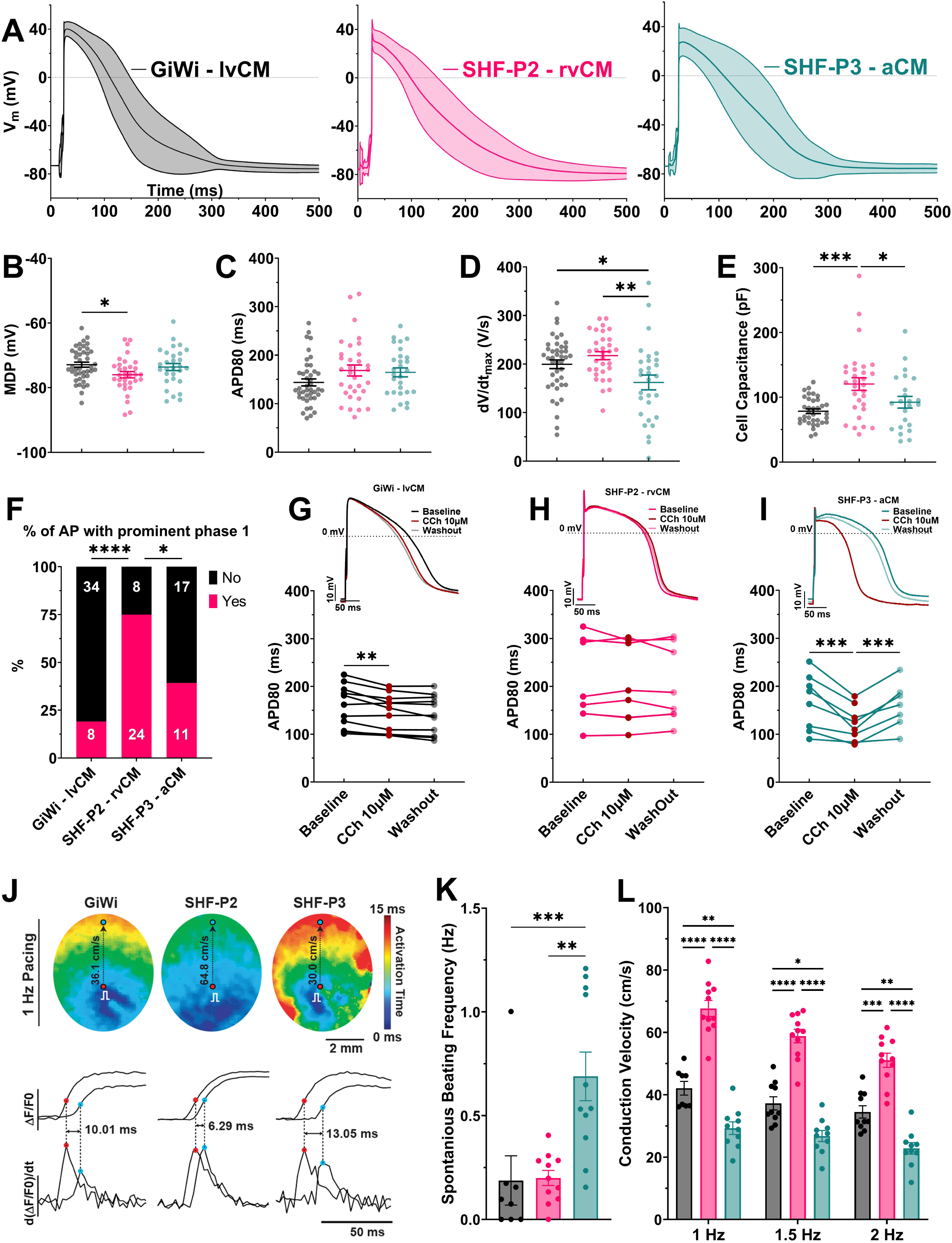
Electrophysiology analysis of the hPSC-CMs differentiated from GiWi, SHF-P2 and SHF-P3 protocols. DF19-9-11T hiPSC-CMs were differentiated for 20 days using the GiWi (lvCM, black/grey), SHF-P2 (rvCM, magenta) or SHF-P3 (aCM, teal) protocols. For single-cell recordings (A–I), cells were dissociated, plated at low density on glass coverslips and recorded 3–7 days later in the perforated-patch current-clamp configuration at 37 °C. (A) Average action potential waveforms recorded during 1 Hz pacing for GiWi-lvCM (left), SHF-P2–rvCM (center) and SHF-P3–aCM (right). Solid line, mean; shaded area, ± SD. (B–E) Action potential and passive membrane parameters measured at 1 Hz: (B) maximum diastolic potential (MDP), corrected for the liquid junction potential; (C) action potential duration at 80 % repolarization (APD80); (D) maximum upstroke velocity (dV/dt_max); and (E) whole-cell capacitance. Each symbol represents one cell; horizontal bars show mean ± SEM. n = 42 (GiWi-lvCM), 32 (SHF-P2–rvCM) and 29 (SHF-P3–aCM) cells. Groups were compared by ordinary one-way ANOVA with Tukey’s multiple comparisons test. *p<0.05, **p<0.01, ***, p<0.005 (F) Proportion of cells displaying prominent phase 1 repolarization, defined as a repolarization rate of < −2.5 V/s following the upstroke (see Supplemental Figure S20). Numbers within bars give the count of cells in each category. Groups were compared by Pearson’s chi-square test of independence (χ² = 23.45, df = 2, p = 8.1 × 10⁻⁶), followed by pairwise Fisher’s exact tests with Holm–Bonferroni correction for three comparisons. *p<0.05, ****p<0.001 (G–I) Effect of the muscarinic agonist 10 µM carbachol (CCh) on APD80 in (G) GiWi-lvCM, (H) SHF-P2–rvCM and (I) SHF-P3–aCM. Upper insets, representative action potentials at baseline (group color), during CCh (dark red) and after washout (grey). Lower panels, paired APD80 measurements for individual cells at baseline, during CCh and after washout; each line represents one cell. n = 11 (GiWi-lvCM), 8 (SHF-P2–rvCM) and 8 (SHF-P3–aCM) cells. G and H were analyzed by repeated-measures one-way ANOVA; I by a mixed-effects model (REML) owing to one missing value; both with Šídák’s multiple comparisons test. **p<0.01, ***p<0.001 (J–L) Optical mapping of purified hiPSC-CM monolayers plated on soft micropatterned PDMS and loaded with the intracellular Ca²⁺ indicator Calbryte™. (J) Upper, representative isochronal activation maps during 1 Hz pacing; the pacing site is indicated; color bar indicates activation time (0–15 ms); scale bar, 2 mm. Lower, representative Ca²⁺ transients (ΔF/F₀) and first derivative d(ΔF/F₀)/dt recorded from two sites within each monolayer, illustrating the inter-site activation delay used to calculate conduction velocity; scale bar, 50 ms. (K) Spontaneous beating frequency of unpaced monolayers; n = 8 (GiWi-lvCM), 11 (SHF-P2–rvCM) and 11 (SHF-P3–aCM) monolayers; ordinary one-way ANOVA with Tukey’s multiple comparisons test. (L) Conduction velocity during pacing at 1, 1.5 and 2 Hz; two-way ANOVA with Tukey’s multiple comparisons test comparing groups within each pacing rate. In L, each symbol represents one monolayer with n= 11 (GiWi-lvCM), 11 (SHF-P2–rvCM) and 10 (SHF-P3–aCM) and bars show mean ± SEM. Statistical significance is indicated as *p < 0.05, **p < 0.01, ***p < 0.001, ****p < 0.0001; comparisons without a bracket were not statistically significant.

We noted that some action potentials in all groups exhibited a rapid repolarization after the upstroke (phase 1), which in native cardiomyocytes is generated principally by the transient outward K⁺ current, I_to_^10,74,75^. Using a dV/dt cutoff of < −2.5 V/s to identify cells with prominent phase 1 repolarization (**Supplementary Figure S21**), the proportion of cells displaying phase 1 differed across the three hiPSC-CM preparations. Phase 1 was present in 75.0 % of SHF-P2–rvCM (24/32; 95 % CI 57.9–86.7), 39.3 % of SHF-P3–aCM (11/28; 95 % CI 23.6–57.6) and 19.0 % of GiWi-lvCM (8/42; 95 % CI 10.0–33.3). Pairwise comparison showed that SHF-P2–rvCM differed from both GiWi-lvCM (p = 4.9 × 10⁻⁶) and SHF-P3–aCM (p = 0.017), whereas GiWi-lvCM and SHF-P3–aCM did not differ from one another (p = 0.099) (**Figure 7F**). Overall, characterization of APs from the three hiPSC-CM preparations showed resting potentials within 3 mV of one another and comparable APD80, with the populations separated by a slower upstroke in SHF-P3– aCM and by larger cell size and a markedly higher prevalence of phase 1 repolarization in SHF-P2–rvCM.

To test for chamber-specific pharmacology of the GiWi-lvCM, SHF-P2-rvCM and SHF-P3-aCM preparations, the day 20 differentiated CM populations were challenged with the muscarinic agonist carbachol (CCh, 10 µM), which in native cardiac tissue activates the atrial-selective acetylcholine-gated K⁺ current (I_K,ACh_) via M2 receptor–Gᵢ–GIRK1/4 signaling.^76–79^ In GiWi-lvCM, during CCh superfusion there was a small decrease in APD80 from 157.7 to 145.0 ms (p = 0.0019) that did not recover on washout (140.9 ms; p = 0.41), consistent with a gradual drift in APD in this preparation rather than a receptor-mediated response (**Figure 7G**). SHF-P2–rvCM showed no clear effect of CCh or its washout (average APD80 of 204.7, 202.4 and 200.5 ms, respectively) (**Figure 7H**). In contrast, SHF-P3–aCM exhibited a marked shortening of APD80 from 167.1 to 122.2 ms (p = 0.0003) with superfusion of CCh that reversed fully on washout to 166.2 ms (p = 0.0006) (**Figure 7I**). A reversible, receptor-mediated repolarization response to muscarinic stimulation provides strong electrophysiological evidence for a distinct atrial-like phenotype in SHF-P3–aCM relative to the ventricular GiWi-lvCM and SHF-P2–rvCM.

To further characterize the functional properties of the distinct cell preparations, we performed optical mapping using the intracellular calcium-sensitive fluorescent dye Calbryte™ on DF19-9-11T hiPSC-CMs differentiated from the GiWi, SHF-P2 and SHF-P3 protocols. The hiPSC-CMs were purified and plated as a monolayer on a soft micropatterned polydimethylsiloxane (PDMS) platform that facilitates a biomimetic environment to promote maturity.^80^ All cell preparations formed confluent monolayers that were amenable to optical mapping and captured at 1 Hz pacing, with activation spreading uniformly from the pacing site across the field of view (**Figure 7J, upper**). Local activation was resolved from the maximum of the first derivative of the Ca²⁺ transient, d(ΔF/F₀)/dt (**Figure 7J, lower**). Conduction velocity (CV) differed significantly among the three preparations at each pacing rate tested, with SHF-P2–rvCM the fastest, GiWi-lvCM intermediate and SHF-P3–aCM the slowest (1 Hz: 67.7 ± 2.6, 36.5 ± 0.2 and 29.3 ± 2.0 cm/s; mean ± SEM; n = 11, 8 and 10 monolayers) (**Figure 7J,L**). Conduction velocities in all three preparations were close to the 30–100 cm/s range reported for adult human ventricular myocardium,^81^ and CV decreased progressively with increasing pacing frequency from 1 Hz to 2 Hz in every group, consistent with the rate dependence of conduction in native myocardium (**Figure 7L**). Spontaneous frequency was higher in SHF-P3–aCM (0.69 ± 0.12 Hz) than in either GiWi-lvCM (0.19 ± 0.12 Hz) or SHF-P2–rvCM (0.20 ± 0.04 Hz) (**Figure 7K**), comparable with the higher rate observed in iPSC-derived atrial CMs in previous studies.^82,83^ We also characterized the properties of the calcium transients (CaTs). CaT amplitude measured via fluorescence intensity was significantly smaller in SHF-P3–aCM than in GiWi-lvCM, while SHF-P2–rvCM were intermediate (Supplementary Figure S21). CaT duration at 30, 50 and 80 % decay was significantly shorter in SHF-P3–aCM than in the two ventricular preparations (**Supplementary Figure S22**). These results align with previously reported differences in calcium transient amplitude between primary atrial and ventricular myocytes.^84^ Furthermore, the initial decay phase of the CaT is faster in SHF-P2-rvCMs as shown by the shorter CaTD30 at all three pacing rates in SHF-P2-rvCMs compared to GiWi-lvCMs (**Supplementary Figure S22D**). Overall, optical mapping of intracellular calcium demonstrated clear cell-to-cell conduction with rate-dependent slowing of CV approaching the behavior of native myocardium, and SHF-P2–rvCM showed the fastest conduction, while SHF-P3–aCM was distinguished by faster spontaneous activity, slower conduction and smaller, faster-decaying calcium transients consistent with atrial physiology relative to ventricular.

## DISCUSSION

Many inherited and acquired heart diseases exhibit chamber-specific pathology; therefore, chamber-specific hPSC-CM models are needed to investigate mechanisms of disease and develop therapies. Here, we developed novel protocols to generate SHF progenitors by activating FGF signaling following mesoderm formation and subsequently differentiation to rvCM by inhibition of Wnt and with additional retinoic acid treatment to aCMs. The generation of SHF progenitors was demonstrated by the large population of CXCR4^+^ cells, a marker for SHF, as well as the gene expression profile. The differentiated cells from the SHF progenitors were defined by scRNA-seq transcriptomes, and SHF-derived rvCMs and aCMs were characterized by electrophysiology, the gene expression profile. The differentiated cells from the SHF progenitors were defined by scRNA-seq transcriptomes, and the SHF-derived rvCMs and aCMs were characterized by electrophysiology, Ca^2+^cycling and pharmacology ^78,79^.

The study identified several novel marker genes for the chamber-specific hPSC-CMs. *FGF12* is specific and highly expressed in the GiWi-immature lvCMs, consistent with its expression in human LV during development ^63^. *ABRA* is exclusively expressed in the SHF-P2-rvCMs, along with *A2M* and *MEF2C* which are significantly upregulated DEGs in the SHF-P2-rvCMs and can be used as marker genes for rvCMs. Interestingly, NR2F2 was identified as an atrial marker gene but expressed at a low level in the SHF-P3-aCMs, in contrast, VWDE is specifically and highly expressed in SHF-P3-aCMs and SANs.

The day 20 differentiated rvCMs from our SHF protocol show more mature phenotype with higher expression of ventricular hPSC-CM maturation makers including *MYLl2*, *TNNI3* and *MYH7* relative to the day 20 GiWi-lvCM populations. Furthermore, the gene expression by qRT-PCR and our scRNA-seq data show *NPPA* is highly expressed in the SHF-P2-rvCMs while *HEY2* is expressed higher in GiWi-immature lvCMs, suggests predominantly trabecular CMs compromise the rvCMs compared to more compact CMs in the GiWi-lvCMs.

The chamber-specific hPSC-CMs generated from GiWi, SHF-P2, SHF-P3 protocols show more similar gene expression profiles to the human fetal heart tissue ranges from 5-25 weeks of gestation (HE_5w-HE_25w) than the counterparts of the hPSC-CMs generated from Yang et al.^43^. The transcriptomes of GiWi-immature lvCM and lvCM are well aligned with the human fetal LV tissue. The most similar gene expression profiles when comparing the GiWi-lvCM populations with Yang et al. ^43^ are the proliferating lvCMs. Our SHF-P2-rvCM show more upregulated genes for the SHF including *MEF2C* and *HAND2*, as well as the genes that promote angiogenesis and vasculature maturation than the RVLCM from Yang et al. ^43^. The striking difference when comparing the GiWi-lvCM vs. HE_LV and SHF-P2-rvCM vs HE_RV is *MYH6* expression, which is upregulated in both hPSC-lvCMs and hPSC-rvCMs but downregulated in human fetal LV and RV. Another major difference is *NPPA* expression, which is upregulated in the SHF-P2-rvCM but downregulated in both the RVLCM from Yang et al. ^43^ and the fetal RV tissue ^63^, may suggest more trabecular than compact CMs in the SHF-P2-rvCM than the counterparts in the literature. The *RSPO3* gene (R-spondin 3) that encodes a secreted protein and is an activator of the Wnt/β-catenin signaling is upregulated in the SHF-P3-aCM but downregulated in both the ALCM from Yang et al and the fetal LA+RA tissue.

The day 3.75 progenitors and the day 20 differentiated cells from our protocols used in the scRNA-seq were from the same batch of differentiation and samples were collected at different timepoints in each protocol in parallel. Yang et al. ^43^ used a different strategy in which they sorted the FHF, aSHF and pSHF progenitors and reaggregated the progenitors to further differentiate to the chamber-specific CMs. However, sorting progenitors may interrupt the differentiation trajectory and it is not clear how the manipulation of isolation, sorting and reaggregation impacts the series of developmental events that are precisely temporally regulated, such as progenitor proliferation, the switch from proliferation to commitment of CM fate, gradients of morphogens, signaling pathway network activity, etc., during *in vitro* differentiation_43._

Maturity and heterogeneity of hPSC-CMs are important metrices for establishing robust hPSC-CM models and for therapeutic applications. Our scRNA-seq data show the GiWi protocol differentiated hPSC-CMs comprise three CM populations, immature lvCM, lvCM and proliferating lvCM, and a small population of aCM, with the immature lvCM being the largest population. In contrast, the SHF-P2 protocol generated only one rvCM population with evidence of a mature CM phenotype compared to the GiWi-lvCM populations such as maturation markers expression, larger cell size measured both by imaging and whole-cell capacitance, more negative MDP, and rapid CV in monolayers ^71,82,85^. The underlying cause of the differences in maturity of the preparations is likely related to the distinct protocols employed which include not only different signaling molecules but also different basal media formulations with different vitamins (e.g. ascorbic acid, retinoic acid, etc.), fatty acids and ionic compositions including calcium levels. Similarly, the SHF-P3 protocol also generated one aCM population. These results suggest the SHF protocols of the same duration may generate more mature CMs.

Comparison of the electrophysiology and CaTs of the different hiPSC-CMs shows some anticipated differences but also remarkable similarities. Most distinct were the SHF-P3-aCMs which exhibited slower dV/dt_max_, smaller amplitude CaT with a faster decay phase relative to the ventricular CMs. Furthermore, the SHF-P3-aCMs showed a clear shortening of APD in response to muscarinic stimulation that was absent in the ventricular CMs as anticipated for the atrial-specific I_K,Ach_. A rapid phase 1 repolarization and notch in the action potential was most observed in the SHF-P2-rvCMs consistent with a more prominent I_to_ in RV compared to LV. However, the duration of the average AP was similar between the three preparations despite the anticipation of a shorter APD in aCMs and rvCMs. Comparable APDs may reflect the early developmental stage of the CMs or lack of the complex microenvironment present in the native myocardium that, for example, leads to the well-known transmural gradient in APD.

The relative purity of CMs from the different protocols is another key metric. Optimization of the CHIR concentrations at day 0-1 in GiWi protocol to generate relative pure CMs are oftentimes needed across hPSC lines. However, we didn’t observe much variation of differentiation of rvCM and aCM from line to line with 8 µM CHIR in the SHF-P2 and P3 protocol, so it’s not necessary to optimize the CHIR concentrations for the SHF protocols. Although with the optimized CHIR concentration the GiWi protocol produced on average higher purity of hPSC-CMs (∼70% cTnT^+^ cells) than the SHF protocols (∼50% cTnT^+^ cells) at 20 days differentiation. Also the relatively higher purity of the CMs differentiated from GiWi protocol may be partially due to the proliferating CMs in the GiWi differentiation that continue to increase the CM populations but on the other hand, it may increase the heterogenous CM populations. For some applications, pure CMs are desirable and a variety of purification methods have been described. Surprisingly, we found that the lactate metabolic selection and the magnetic antibody-based cell sorting (MACS) ^86,87^ used commonly with GiWi-lvCMs was ineffective for the SHF-P2-rvCMs. However, we observed that purification using the small molecule inhibitor of MDM2, nutlin-3a, can generate relatively pure SHF-P2-rvCMs (∼90% cTnT^+^ cells) ^68,88^, although the yield was low. These different responses to purification strategies support the distinct phenotypes of the GiWi-lvCMs and SHF-P2-rvCMs.

Overall, we have developed novel monolayer protocols to generate SHF progenitors and differentiate them to rvCMs and aCMs which are benchmarked against lvCMs and native heart. The results show distinct transcriptional patterns and functional properties in the different CM populations. We anticipate the rvCMs and aCMs will prove useful in future disease modeling studies and potentially therapies.

## ACKNOWLEDGMENTS

This research project was supported by WARF/UW-Madison Stem Cell and Regenerative Medicine Pilot Award (JZ), NIH R01HL184472 (TJK, JZ, LLE), U01HL148690 (TJK), R01HL178095 (SP, TJK, JZ) and NSF EEC-1648035 (SP, TJK, JZ) and R01HL170521 (LLE, TJK, JZ). We sincerely thank Dr. Xinmin Zhang at BioInfoRx, Madison, WI to help with the scRNA-seq data analysis, and the University of Wisconsin Biotechnology Center Gene Expression Core and Next Gen Seq Core, and WiCell Research Institute and the Flow Cytometry core labs at UW-Madison for the support of the research.

## AUTHOR CONTRIBUTIONS

J.Z., conceptualization; protocol development and hPSC differentiation; methodology; investigation; validation; acquisition and interpretation of data including flow cytometry, qRT-PCR, immunolabelling, and scRNA-seq process including sample preparation, data analysis, curation and validation, and data sharing and submission to GEO; funding acquisition and writing – original draft and revisions. G.C.K., hPSC differentiation; protocol development and writing; collection of flow cytometry data. V.L. and H.J.J., patch-clamp and optical mapping; methodology; investigation; acquisition and interpretation of electrophysiology data; writing – electrophysiology and methods. J.H., hPSC differentiation, flow cytometry data collection and optical mapping. E.Z., X.F. H.F. and A.V.A., hPSC differentiation; flow cytometry data collection. A.V.G., optical mapping data review and analysis. L.L.E., funding acquisition and writing – revision. S.P.P., methodology; investigation; funding acquisition and writing – draft and revision. T.J.K., conceptualization; methodology; investigation; validation; funding acquisition and writing – draft and revisions.

---

**Figure S1.**
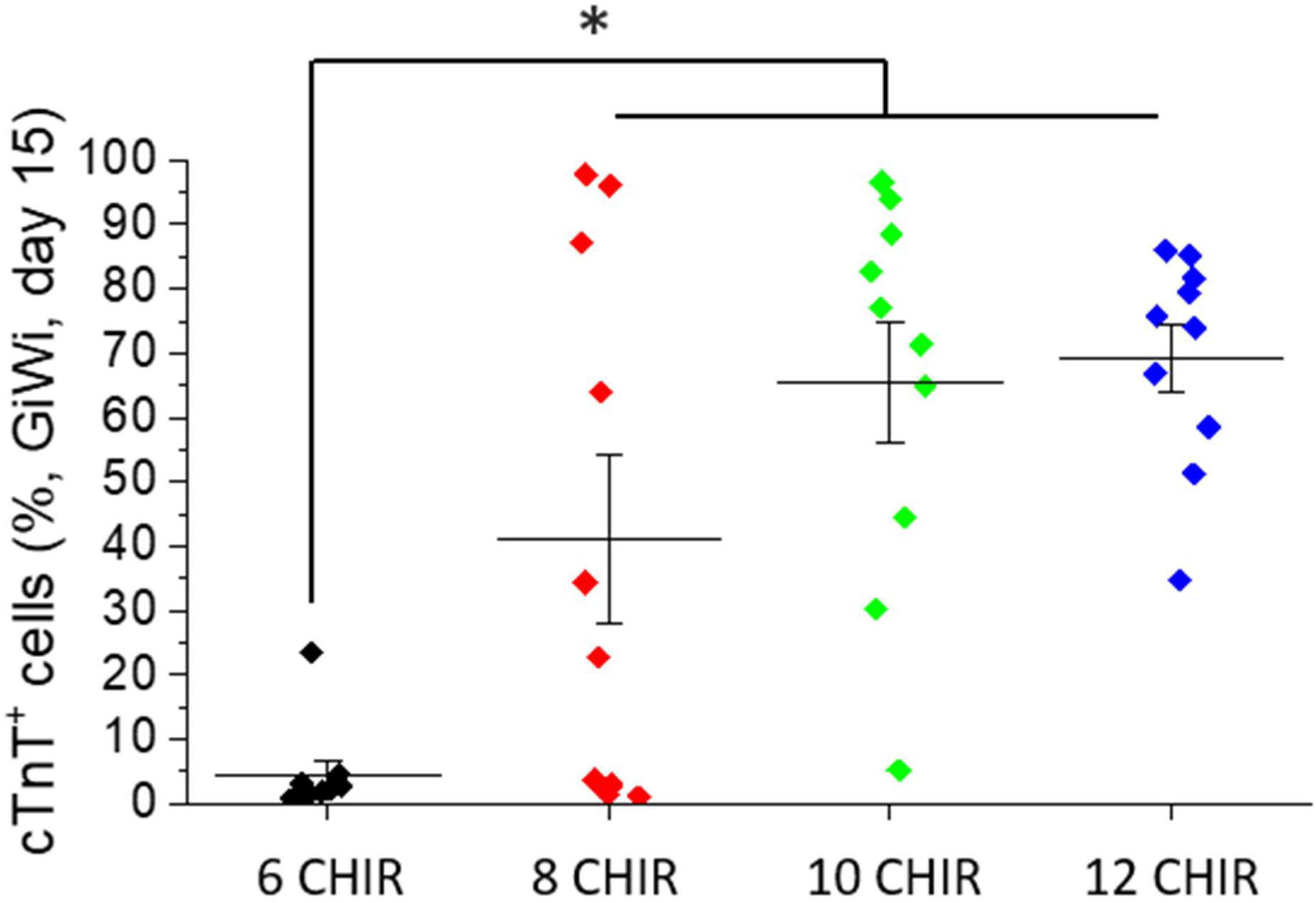
Average of cTnT^+^ cells generated on day 15 of the GiWi protocol with varied CHIR ncentrations of 6-12 µM, measured by flow cytometry. N = 10 batches of differentiation. \**P*<0.05, e-way ANOVA with Tukey post-hoc test.

**Figure S2.**
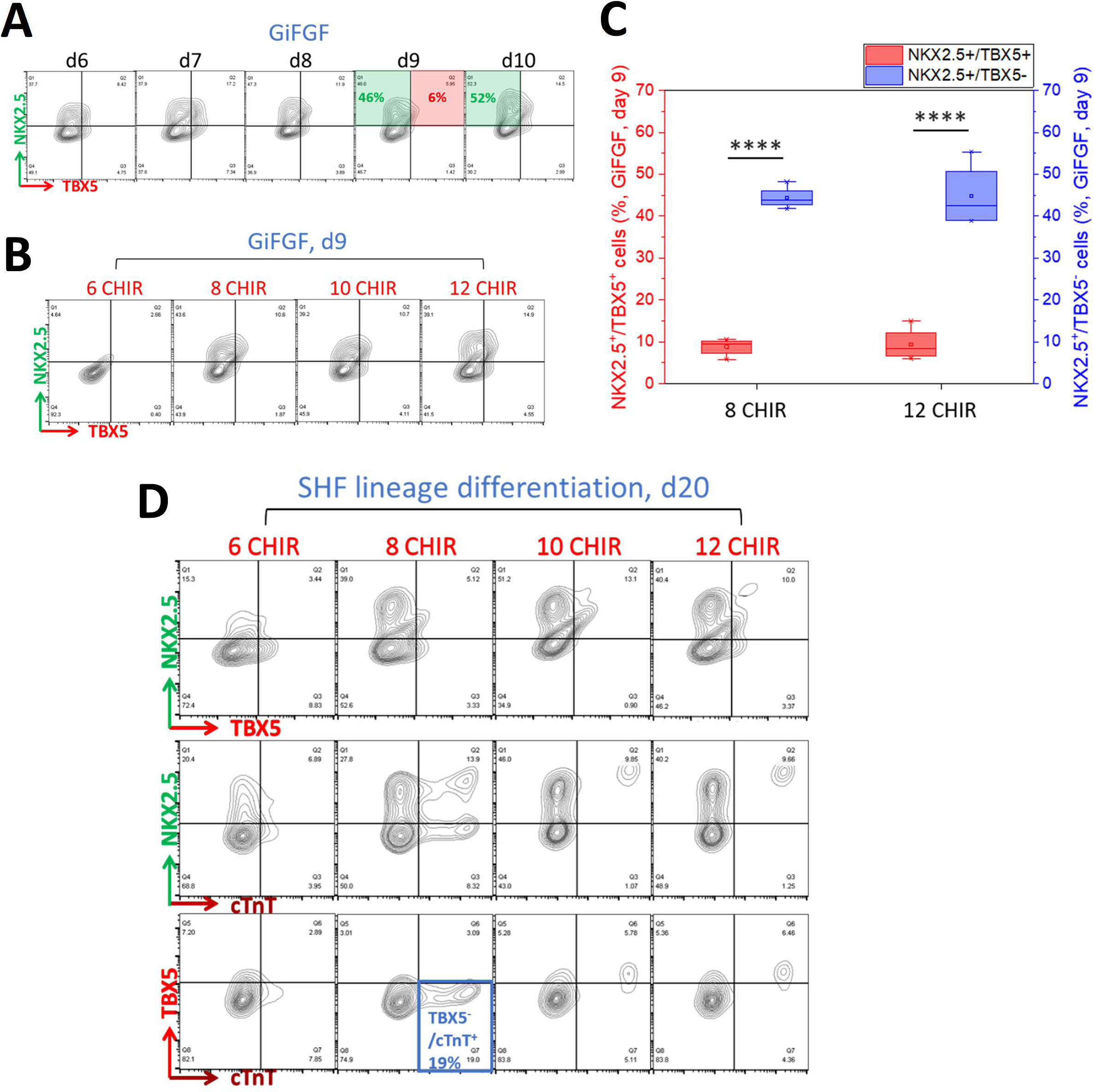
SHF lineage differentiation using the hES3-TBX5^TdTomato/W^/NKX2-5^eGFP/W^ reporter line. Generation of NKX2.5^+^/TBX5^-^SHF progenitors at day 6-10 in the GiFGF protocol measured by flow tometry. **B.** Generation of NKX2.5^+^/TBX5^-^SHF progenitors with varying CHIR concentrations of 6-12 M in the GiFGF protocol measured by flow cytometry. **C.** Average of the NKX2.5^+^/TBX5^+^ and KX2.5^+^/TBX5^-^as the FHF and SHF progenitors (%) at day 9 in the GiFGF protocol, respectively, easured by flow cytometry. **D.** Differentiation of SHF progenitors generated in **B** to TBX5^-^/cTnT^+^ CMs ith treatment of BMP4 (10 ng/ml) and Wnt 5a (100 ng/ml) at day 9 for 4 days and measured by flow tometry at day 20 with varying CHIR concentrations of 6-12 µM at day 0-1. N = 10 batches of fferentiation. **** *P*<0.0001, student t-test.

**Figure S3.**
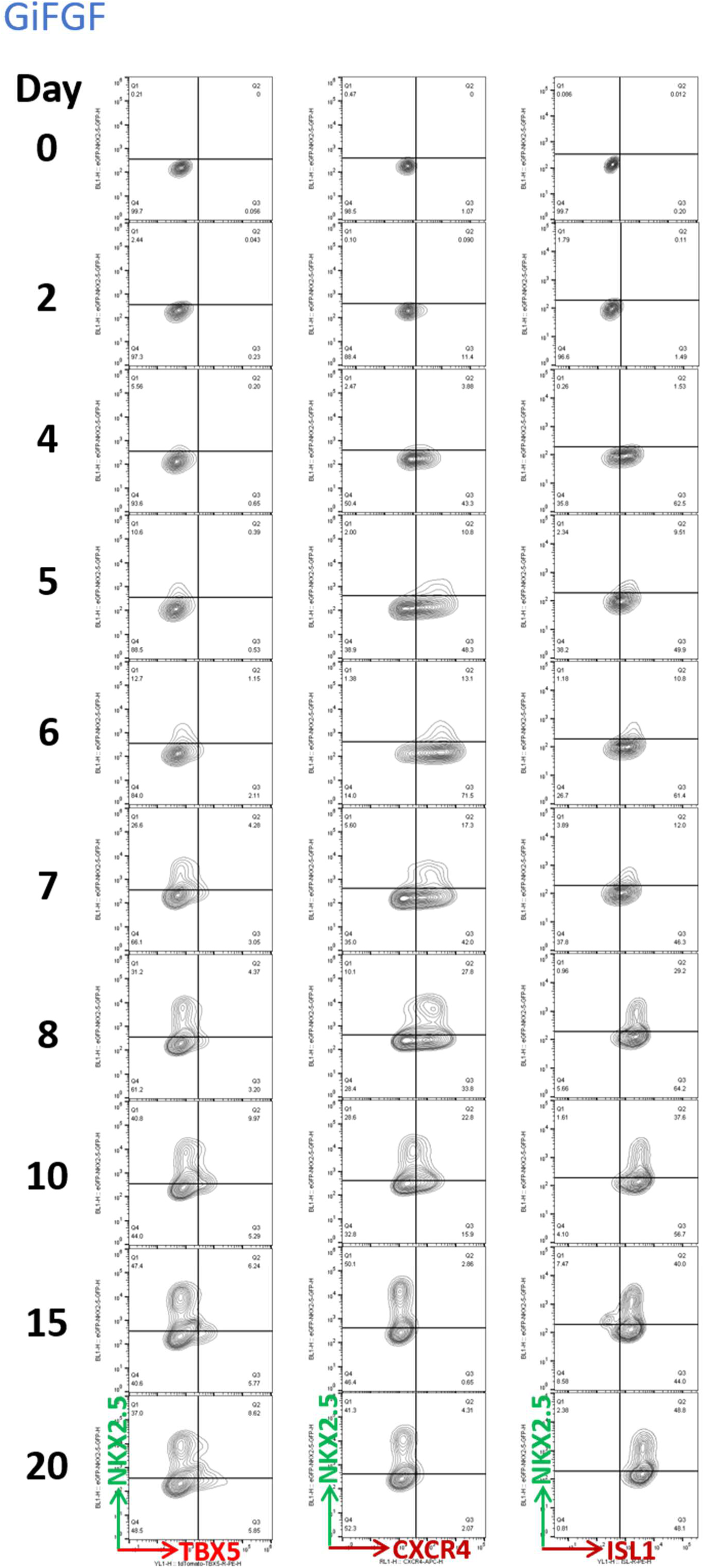
Flow cytometry analysis of CXCR4^+^, Isl1^+^ and NKX2-5^+^ and TBX5^+^ cells generated in the time course of day 0-20 of the GiFGF protocol using the hES3-TBX5^TdTomato/W^/NKX2-5^eGFP/W^ reporter line.

**Figure S4.**
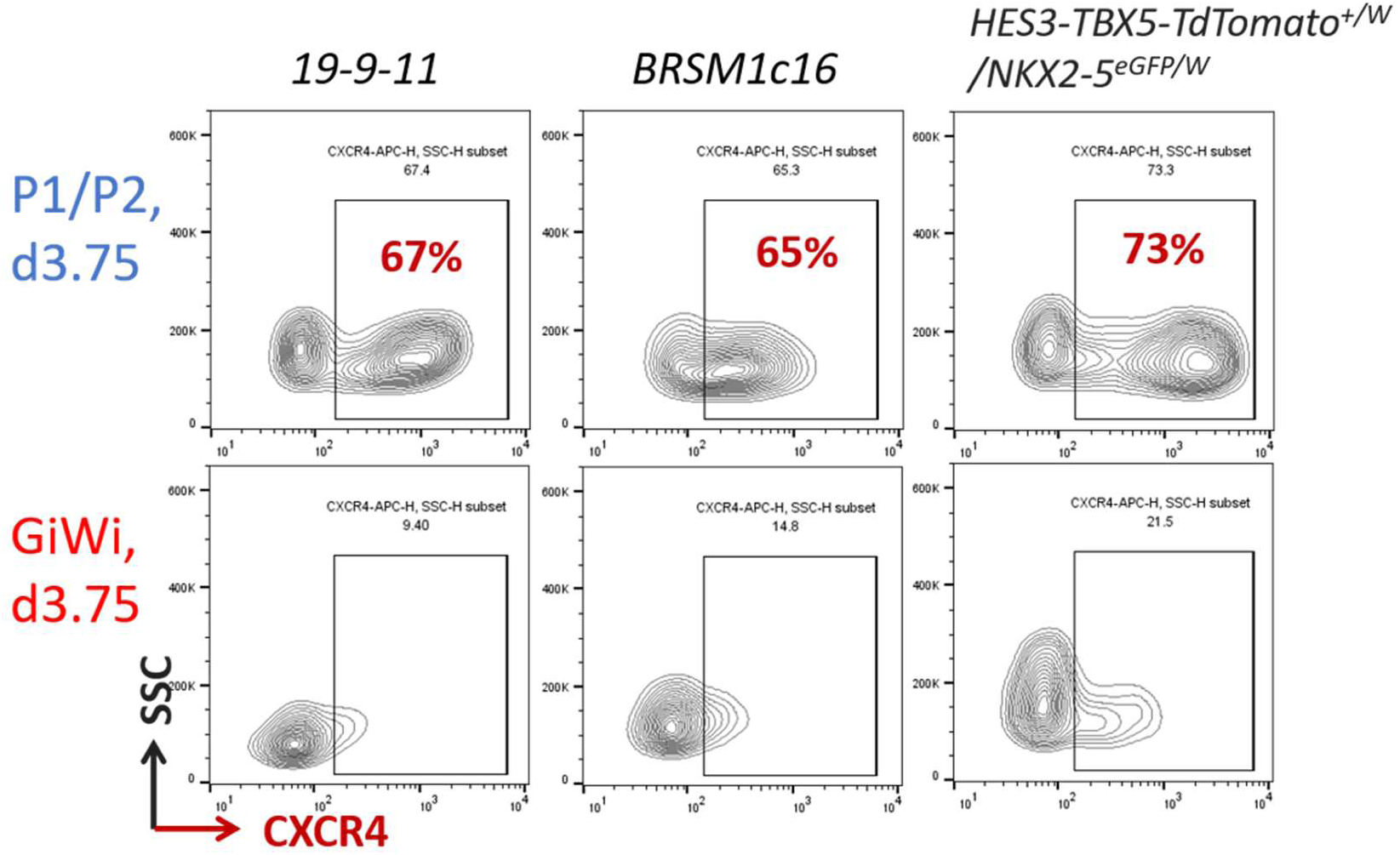
Flow cytometry nalysis of CXCR4^+^ cells enerated at day 3.75 of the HF-P1/P2 and GiWi rotocol in DF19-9-11T and RSM1c16 iPSC lines and ES3-TBX5^TdTomato/W^/NKX2-^eGFP/W^ reporter line.

**Figure S5.**
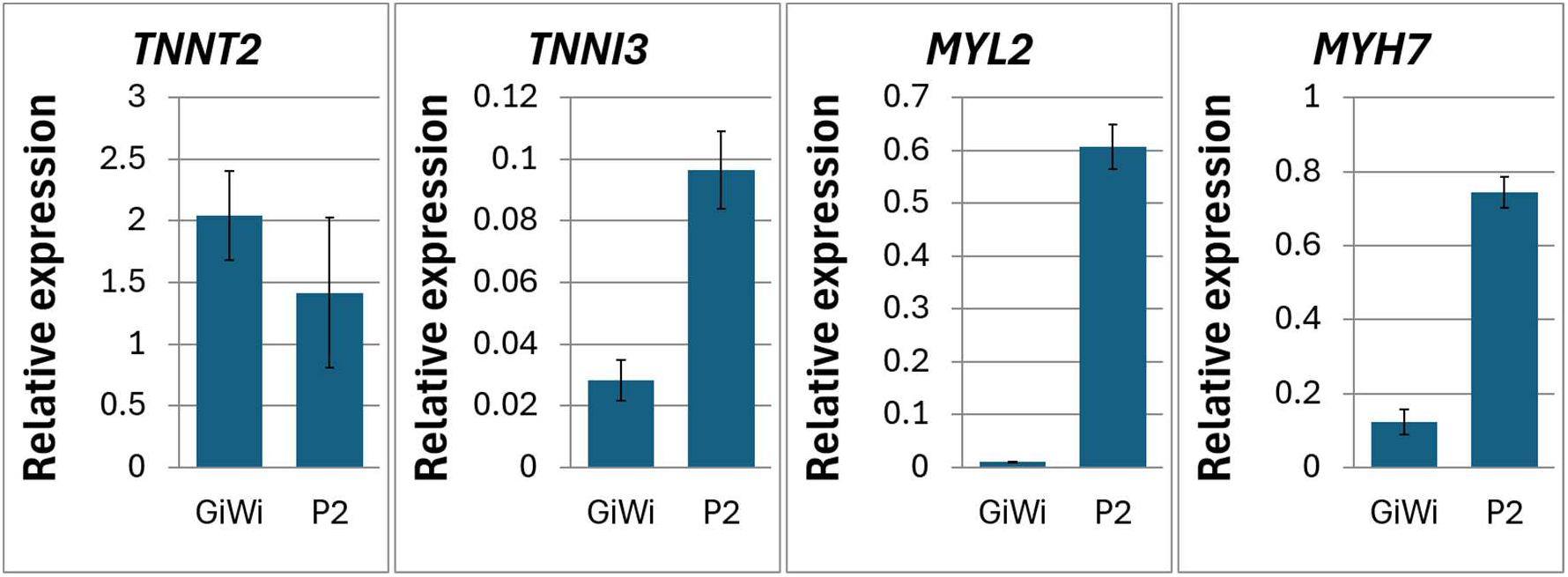
Expression of maturation marker genes for hPSC-CMs measured by qRT-PCR on the day differentiated cells using GiWi, SHF-P2 and P3 protocols.

**Figure S6.**
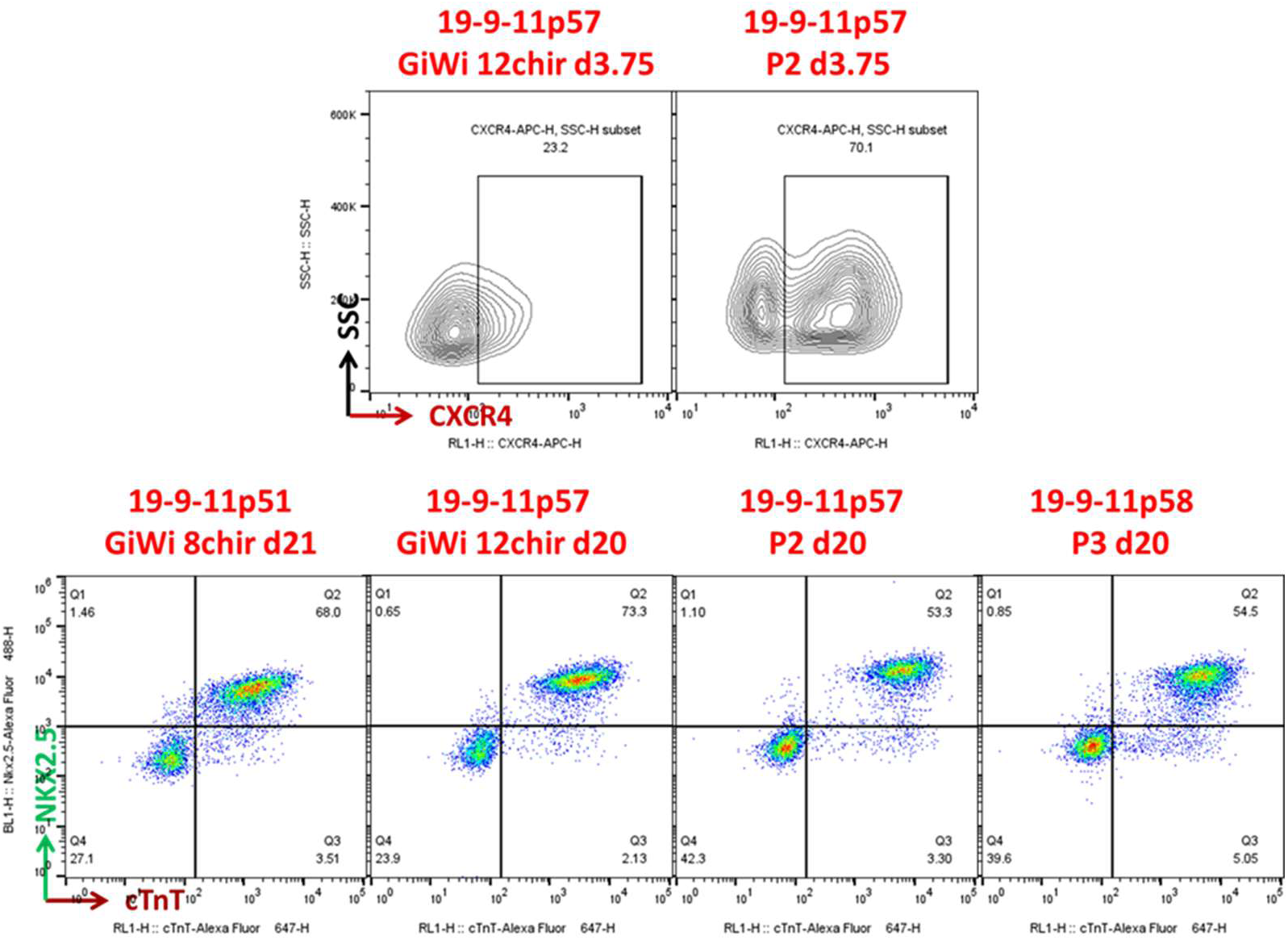
The samples for scRNA-seq including the day 3.75 progenitors from GiWi and SHF-P2 protocols and the day 20 differentiated cells from GiWi-8CHIR, GiWi-12CHIR, SHF-P2 and SHF-P3 protocols underwent flow cytometry for CXCR4^+^ cells generated at day 3.75 and cTnT^+^ cells generated at day 20 in the protocols from DF19-9-11T line.

**Figure S7.**
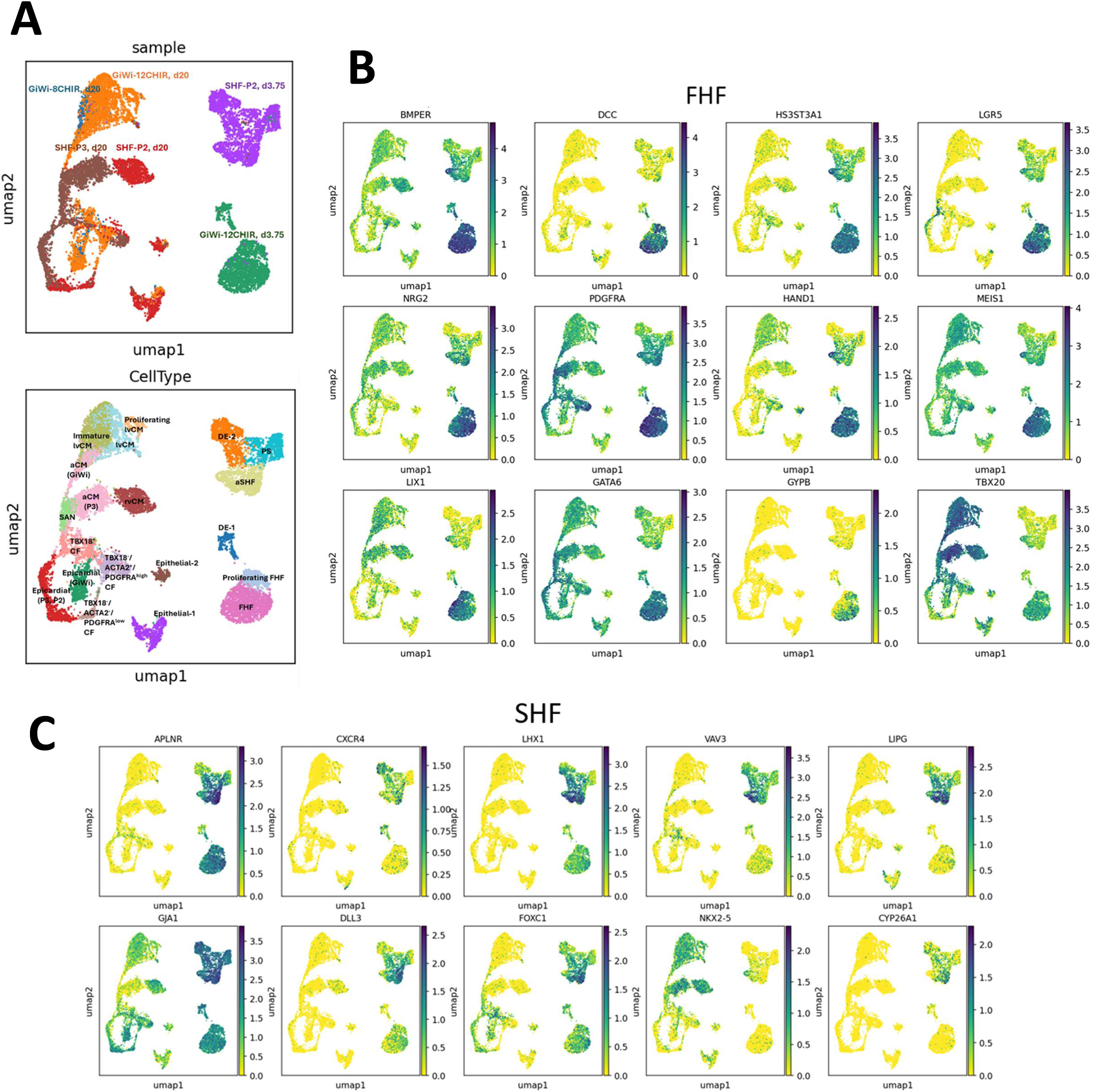
Single cell transcriptomics for heart field-specific gene expression in the day 3.75 ogenitors and day 20 cells differentiated from GiWi, SHF-P2 and P3 protocols. **A.** UMAP in cell type sample view of the day 3.75 and 20 differentiated cells from the protocols. **B.** Embedded plot for ression of marker genes including DCC, GYPB and HAND1 and the DEGs including BMPER, 3ST3A1, LGR5, NRG2, PDGFRA, MEIS1, LIX1, GATA6 and TBX20 that are highly expressed in the 3.75 progenitors derived from the GiWi-12CHIR protocol. **C.** Embedded plot for expression of marker es including CXCR4, LHX1 and FOX1 and the DEGs including APLNR, VAV3, LIPG, GJA1, DLL3, X2-5 and CYPA6A1 that are highly expressed in the day 3.75 progenitors derived from the SHF-P2 protocol.

**Figure S8.**
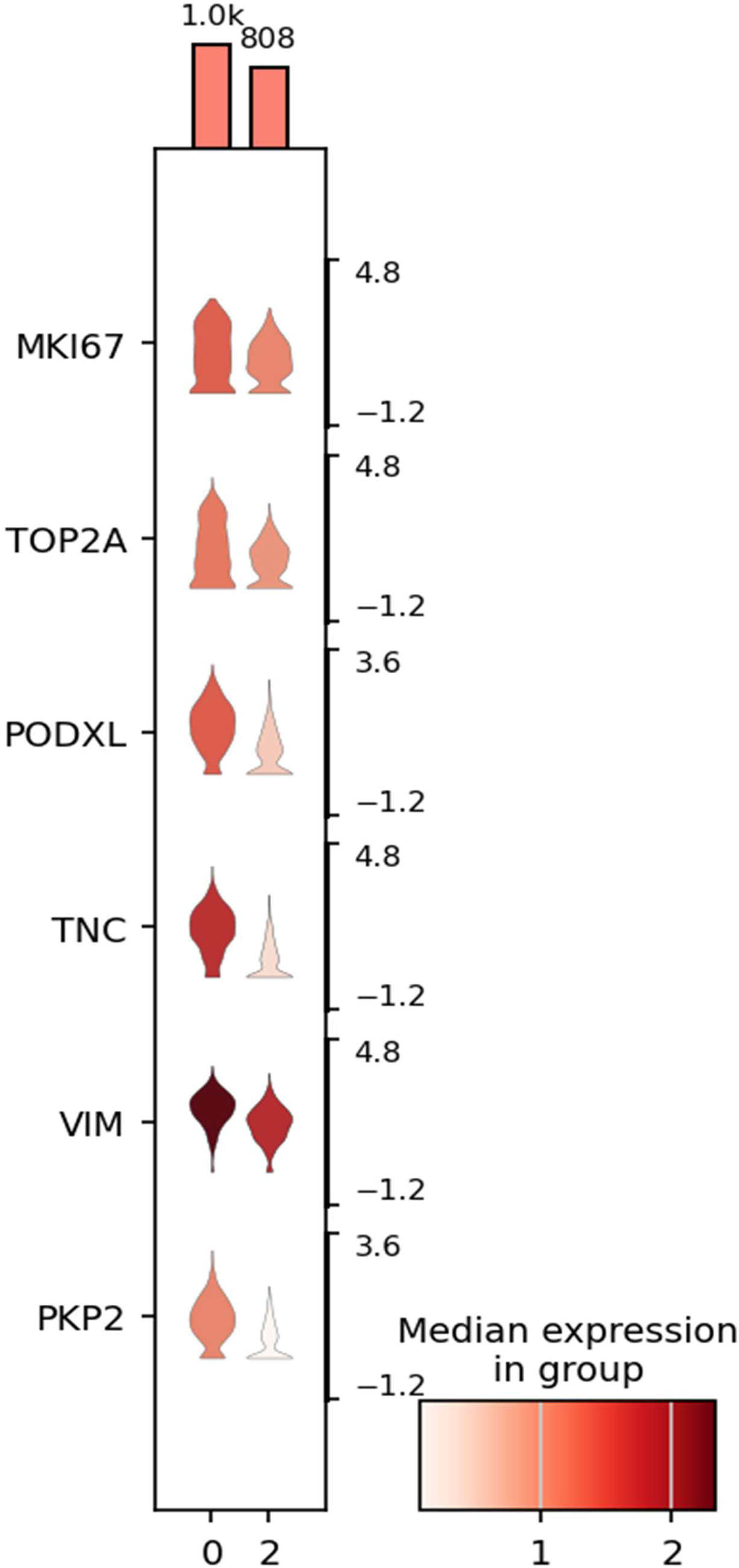
Violin plot of the DEGs in comparison of cluster 0 and 2 of the day 3.75 progenitors nerated from the GiWi-12CHIR protocol. Bars represent cell number for each cluster. Y-axis of violin t represents log_2_(TPM+1) values for gene expression.

**Figure S9.**
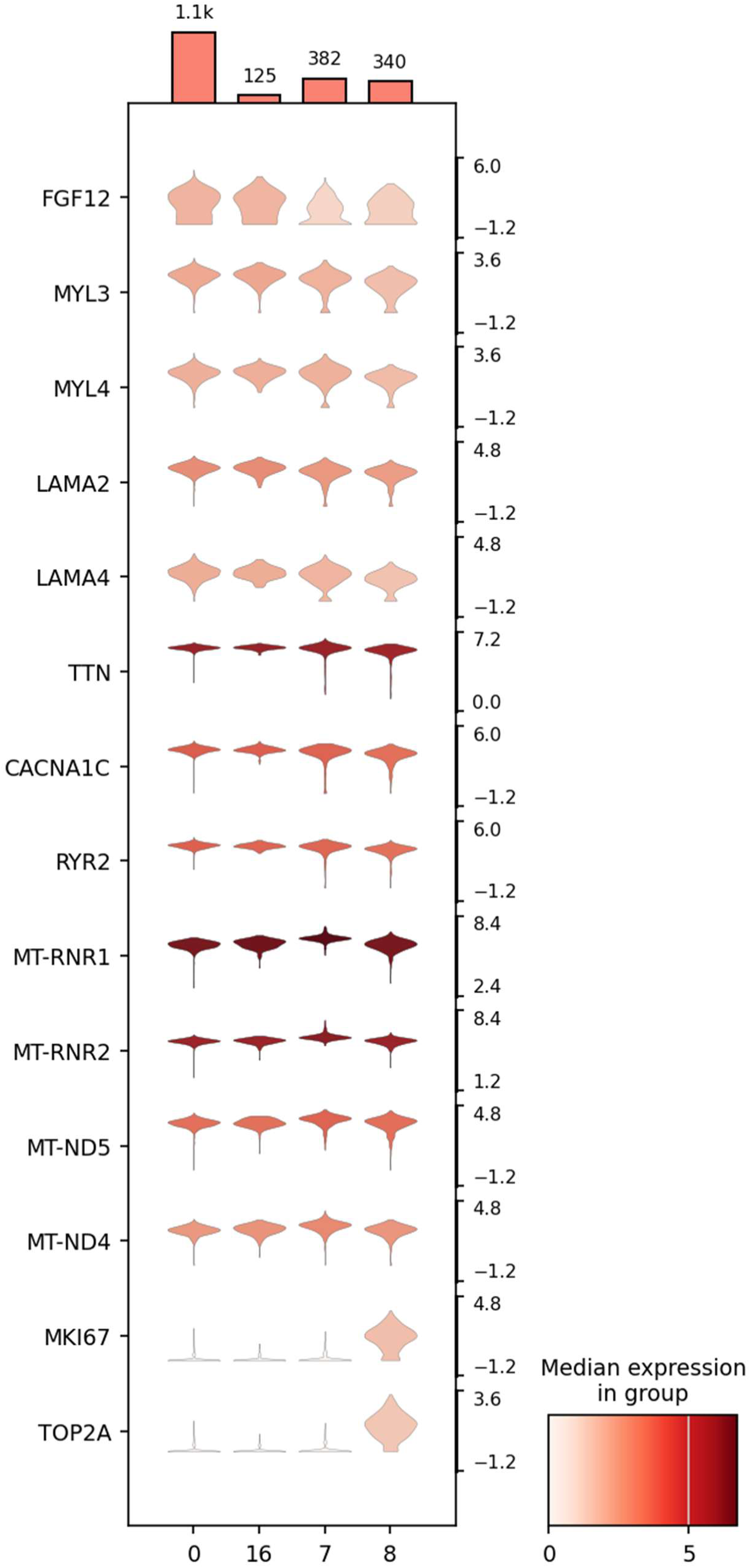
Violin plot of the top DEGs in the immature M (cluster 0 and 16), lvCM (cluster 7) and proliferating lvCM differentiated from the GiWi-12CHIR protocol. Bars represent cell number for each cluster. Y-axis of violin plot represents log2(TPM+1).

**Figure S10.**
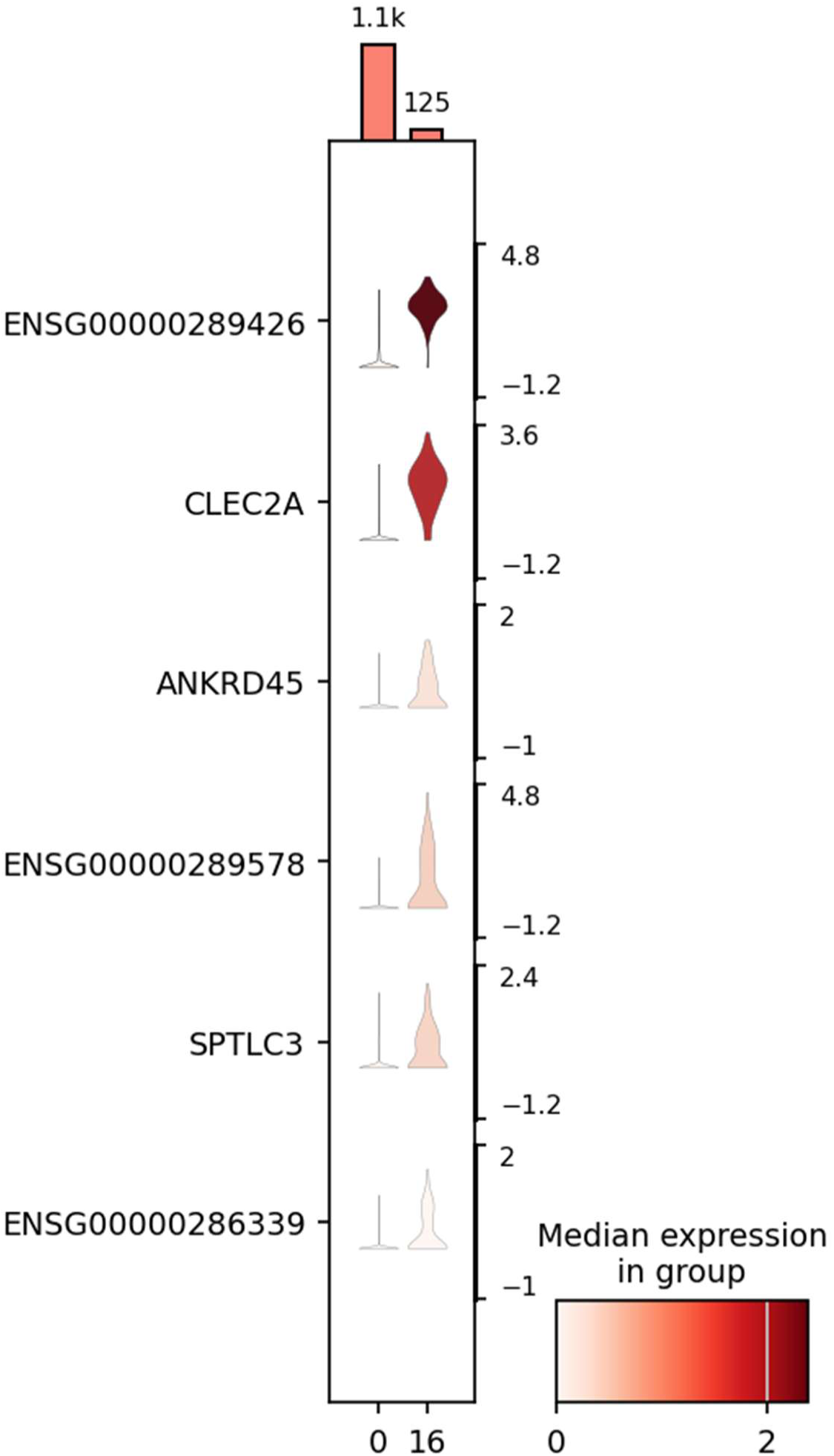
Violin plot of the DEGs that distinguishes cluster 16 from cluster 0 annotated as immature lvCM. Bars represent cell number for each cluster. Y-axis of violin plot represents log_2_(TPM+1) values for gene.

**Figure S11.**
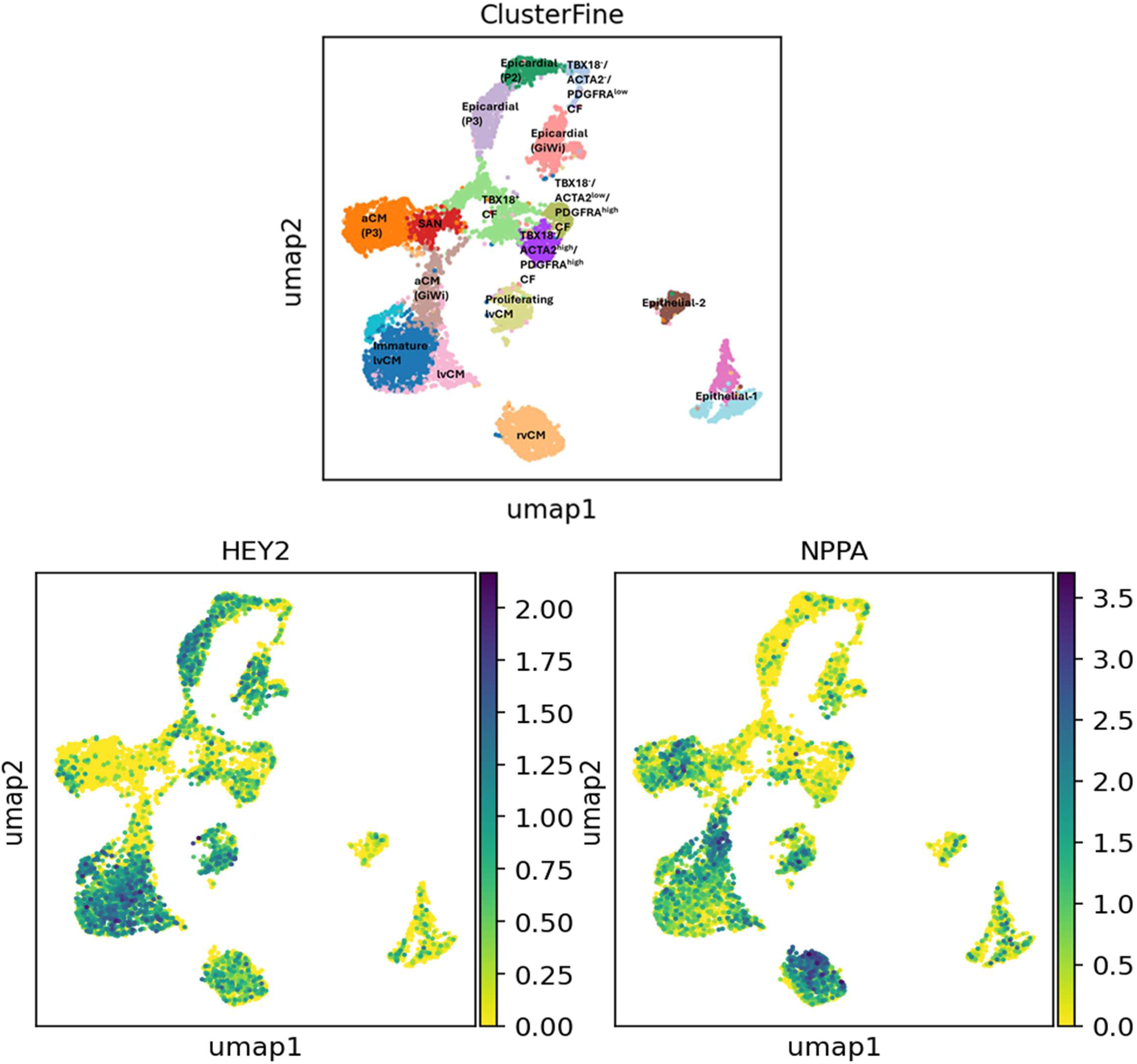
Embedding plot of the scRNA-seq data shows expression of *HEY2* and *NPPA* in all the day differentiated cells from GiWi, SHF-P2 and SHF-P3 protocols.

**Figure S12.**
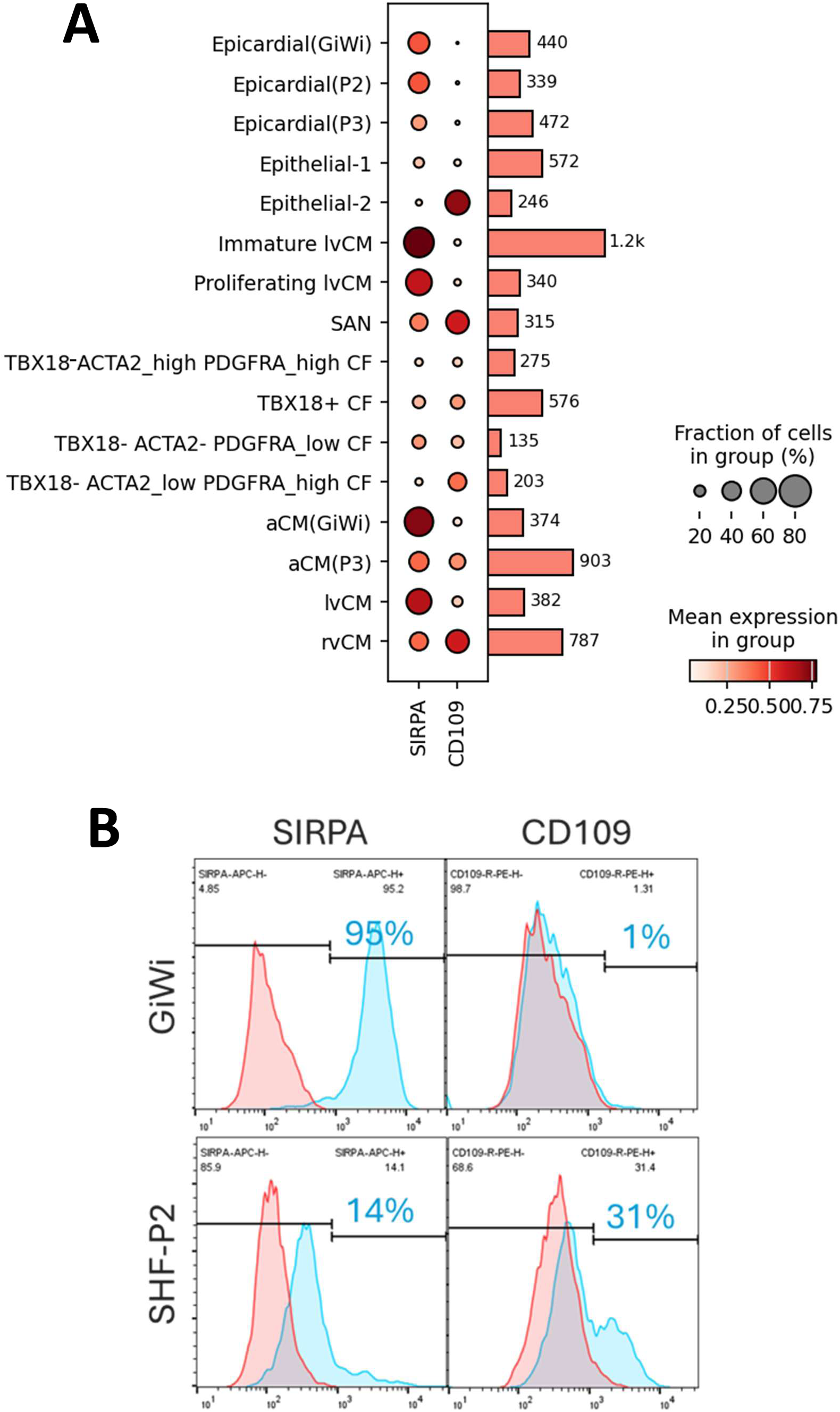
Expression of surface marker SIRPA and CD109 in the GiWi and SHF-P2 differentiated SC-CMs. **A.** scRNA-seq data shows SIRPA is highly expressed in the GiWi-derived lvCMs and aCMs low expression in the SHF-P2-derived rvCMs and SHF-P3-derived aCMs. **B.** Flow cytometry shows the ified hPSC-CMs differentiated from the GiWi protocol highly express SIRPA, but almost no CD109^+^ ls. In contrast, only 14% of the purified rvCMs differentiated from the SHF-P2 protocol are SIRPA^+^ cells, 31% are CD109^+^ cells.

**Figure S13.**
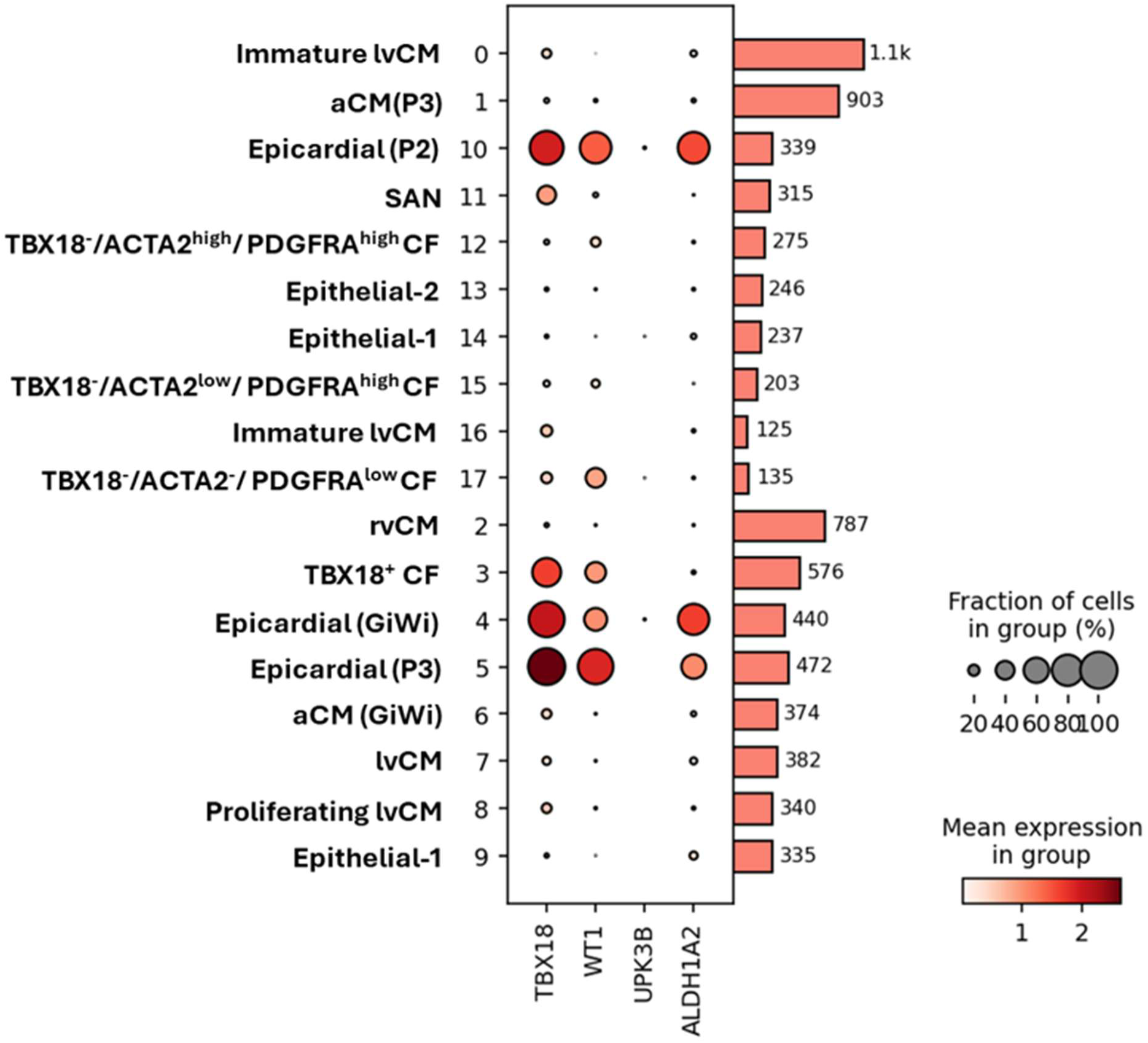
Dot plot for epicardial marker genes expression including ALDH1A2, TBX18 and WT1 in day 20 differentiated cell populations from GiWi, SHF-P2 and P3 protocols. Bars represent cell number for each cluster. Cells with low gene expression of 0-1 were removed.

**Figure S14.**
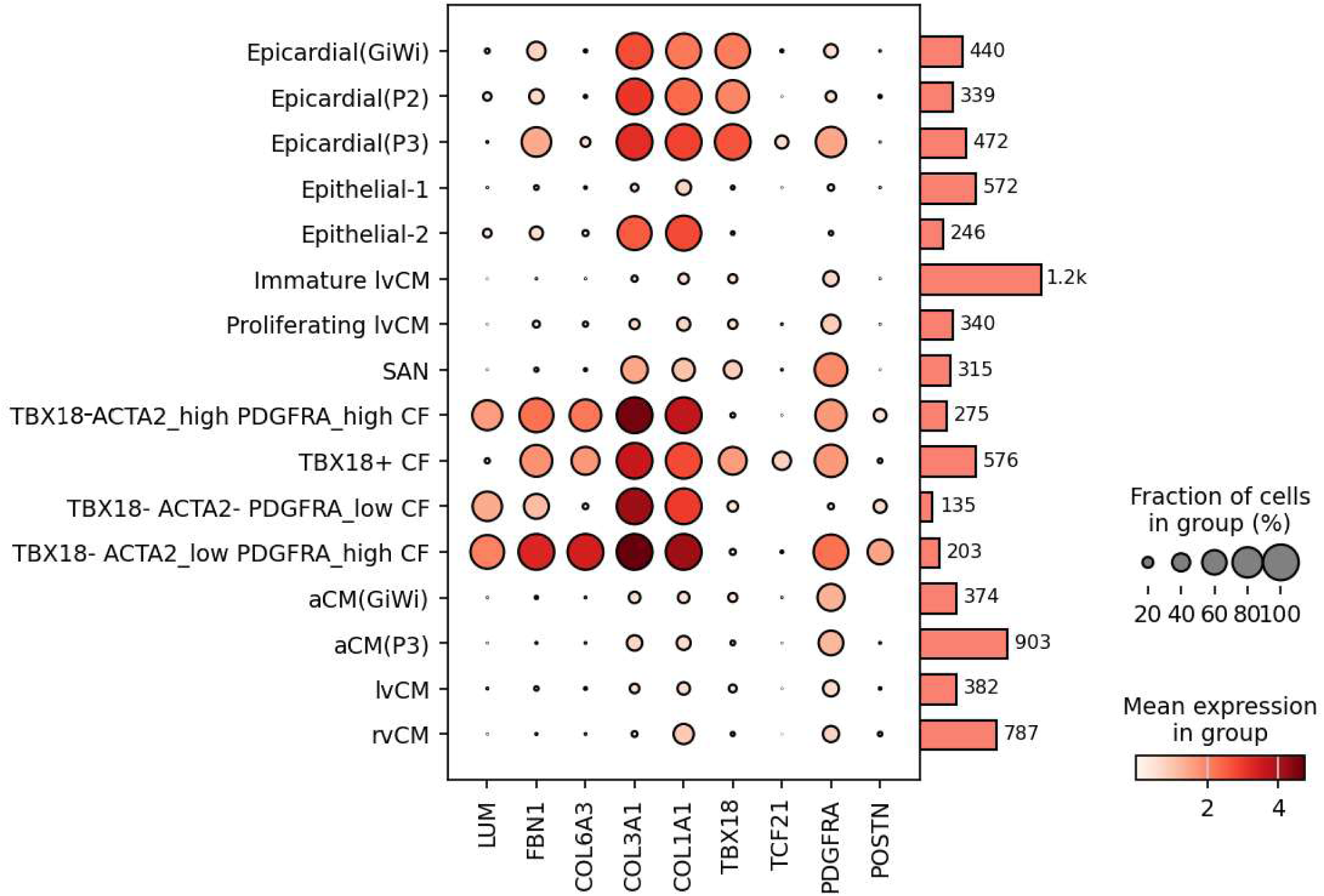
Dot plot for cardiac fibroblast (CF) marker genes expression including LUM, FBN1 and L6A3 that distinguish CF populations from epicardial cells in the day 20 differentiated cells from Wi, SHF-P2 and P3 protocols. Bars represent cell number for each cluster. Cells with low gene ression of 0-1 were removed.

**Figure S15.**
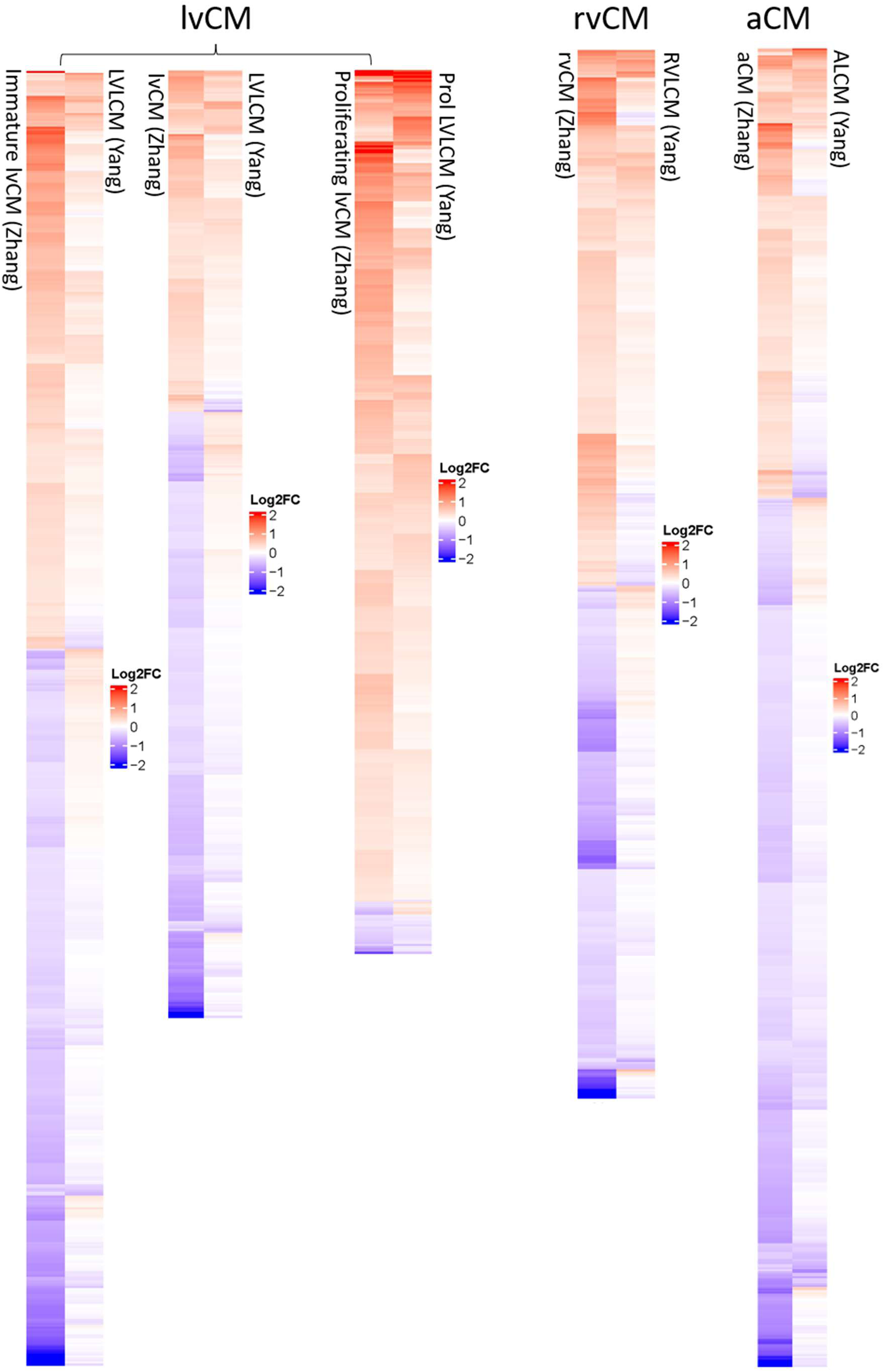
Heatmap of all the shared DEGs to compare the gene expression pattern of the chamber-cific hPSC-CMs derived from our dataset (Zhang) and Yang’s dataset.

**Figure S16.**
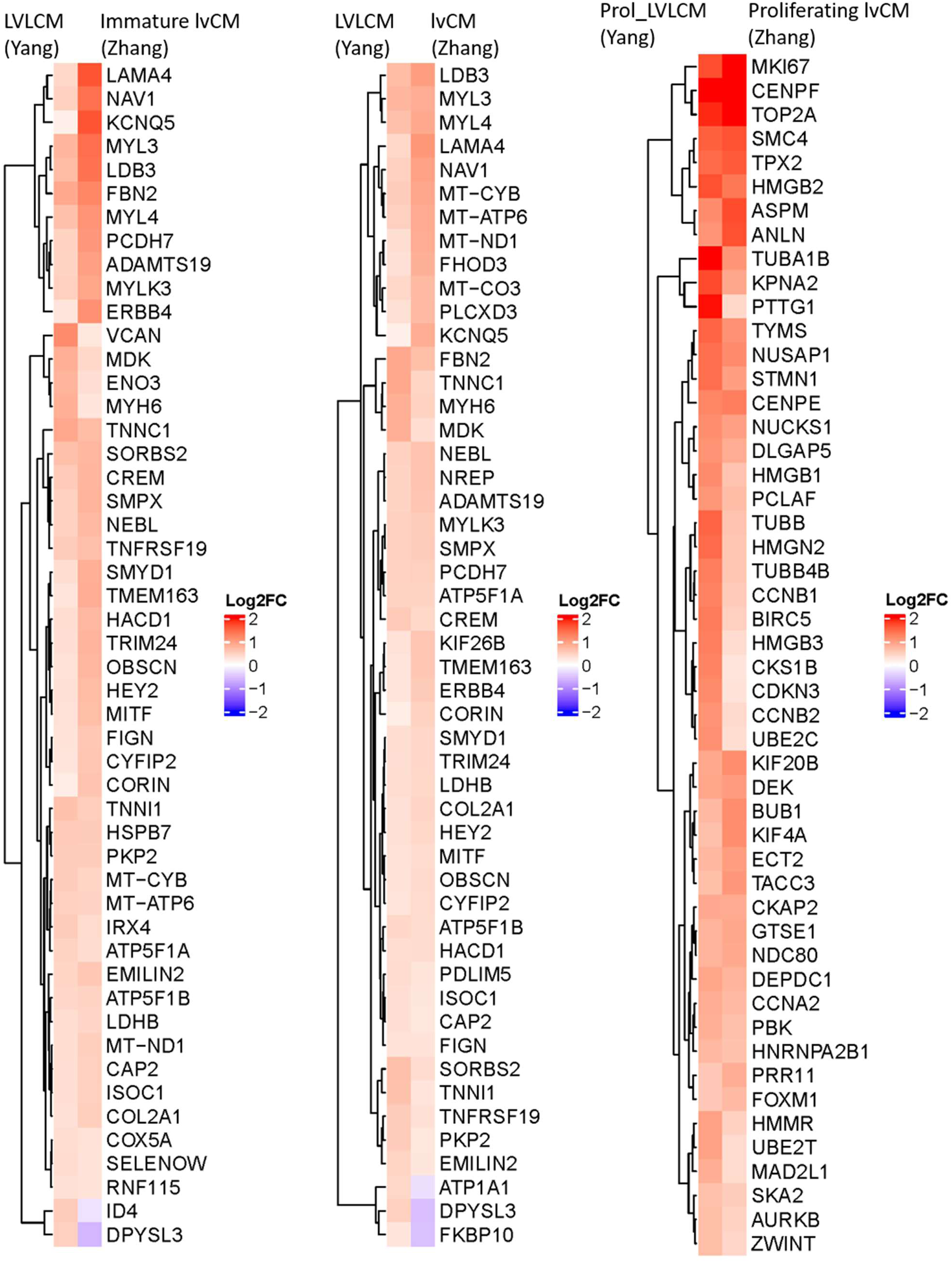
Heatmap of the top 50 most significant upregulated genes ordered by Yang’s dataset to mpare the shared DEGs of LVLCM (Yang) vs. immature lvCM, LVLCM (Yang) vs. lvCM (Zhang), d Prol_LVLCM (Yang) vs. Proliferating lvCM (Zhang).

**Figure S17.**
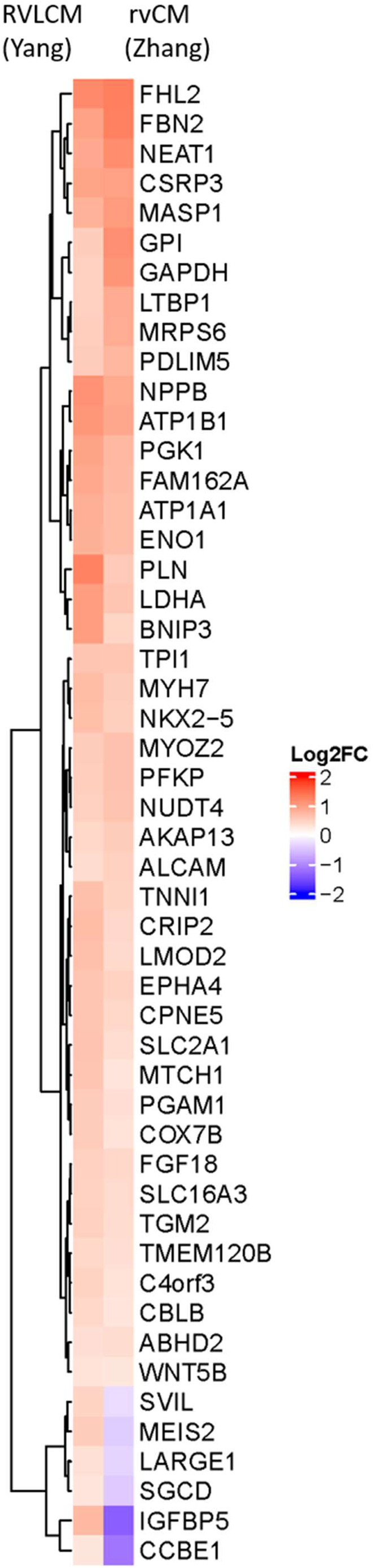
Heatmap of the top 50 most Figure S18. Heatmap of the top 50 most significant upregulated genes ordered by Yang’s dataset to compare the shared DEGs between RVLCM (Yang) and rvCM (Zhang).

**Figure S18.**
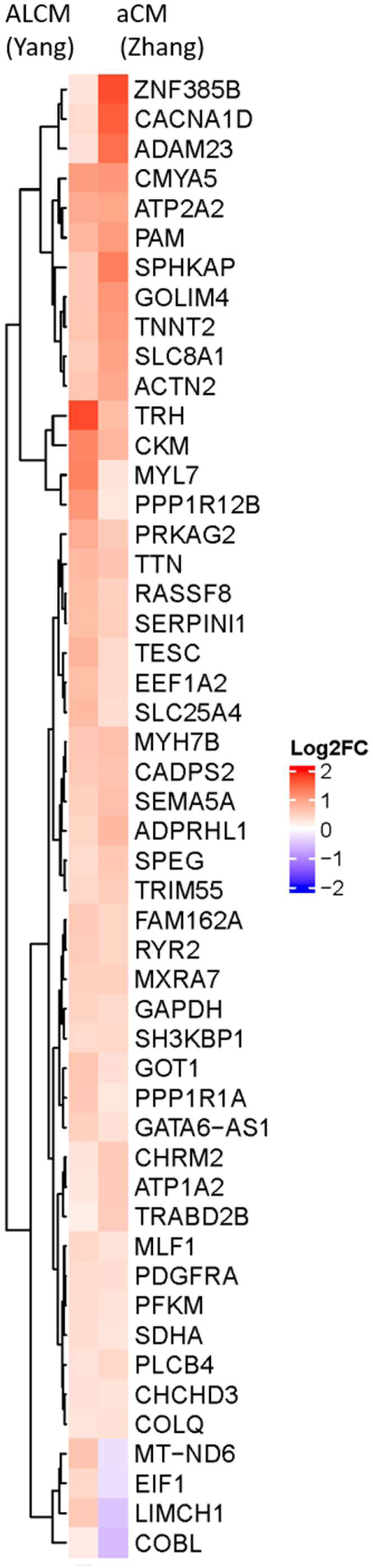
Heatmap of the top 50 most significant upregulated genes ordered by Yang’s dataset to compare the shared DEGs between ALCM (Yang) and aCM (Zhang).

**Figure S19.**
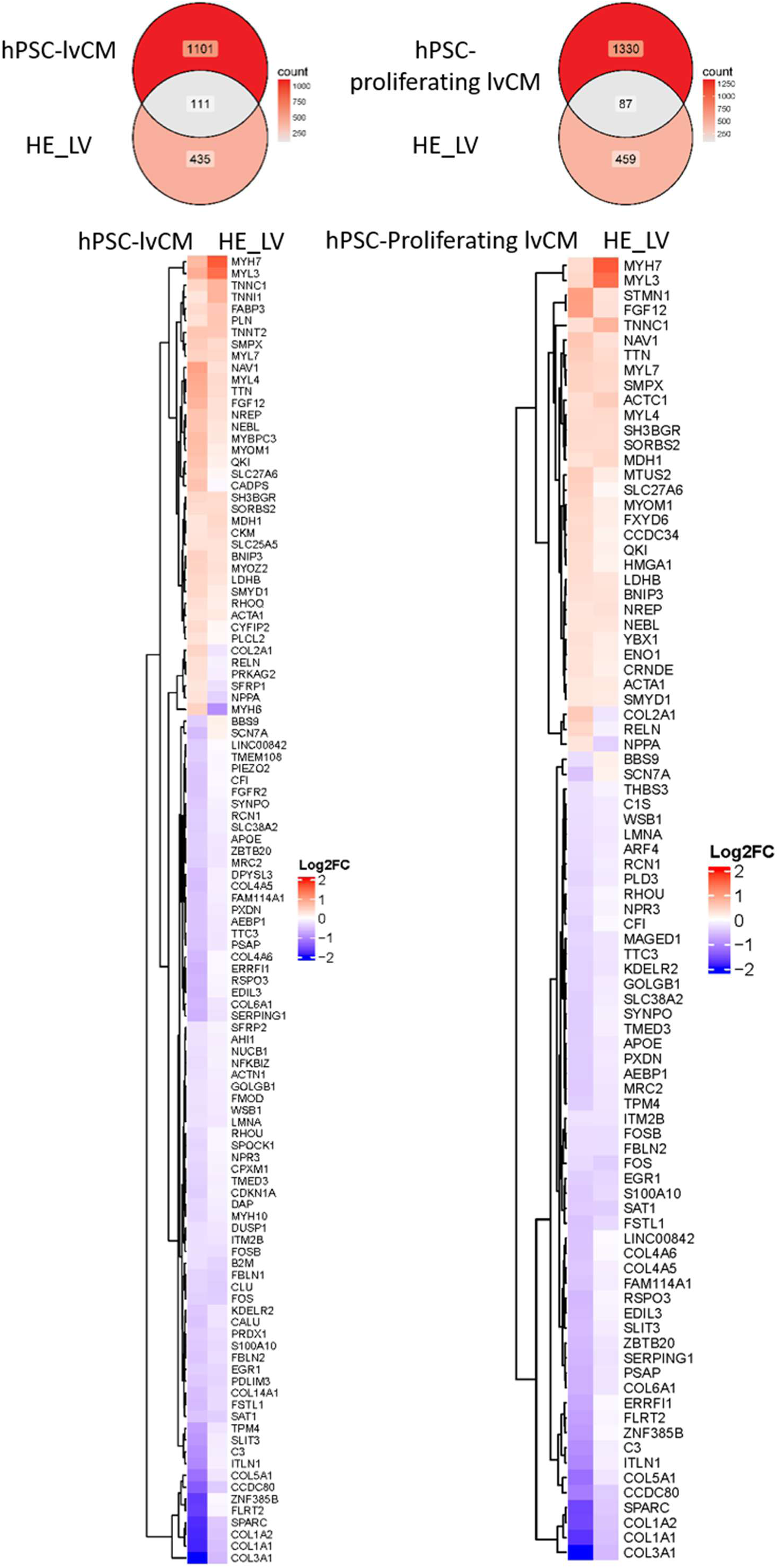
Venn diagram and Heatmap of the shared DEGs to compare the hPSC-lvCM and hPSC-proliferating lvCM from our dataset with the HE_LV from Cui’s dataset.

**Figure S20.**
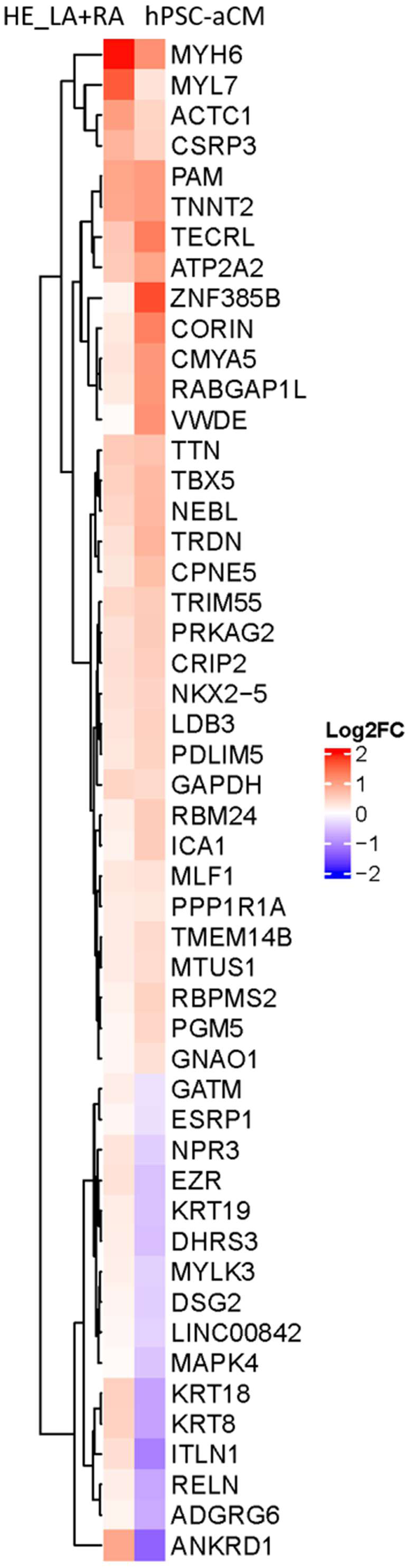
Heatmap of the top 50 most significant upregulated genes ordered by Cui’s dataset to compare the shared DEGs between HE_LA+RA (Cui) and hPSC-aCM (Zhang).

**Figure S21.**
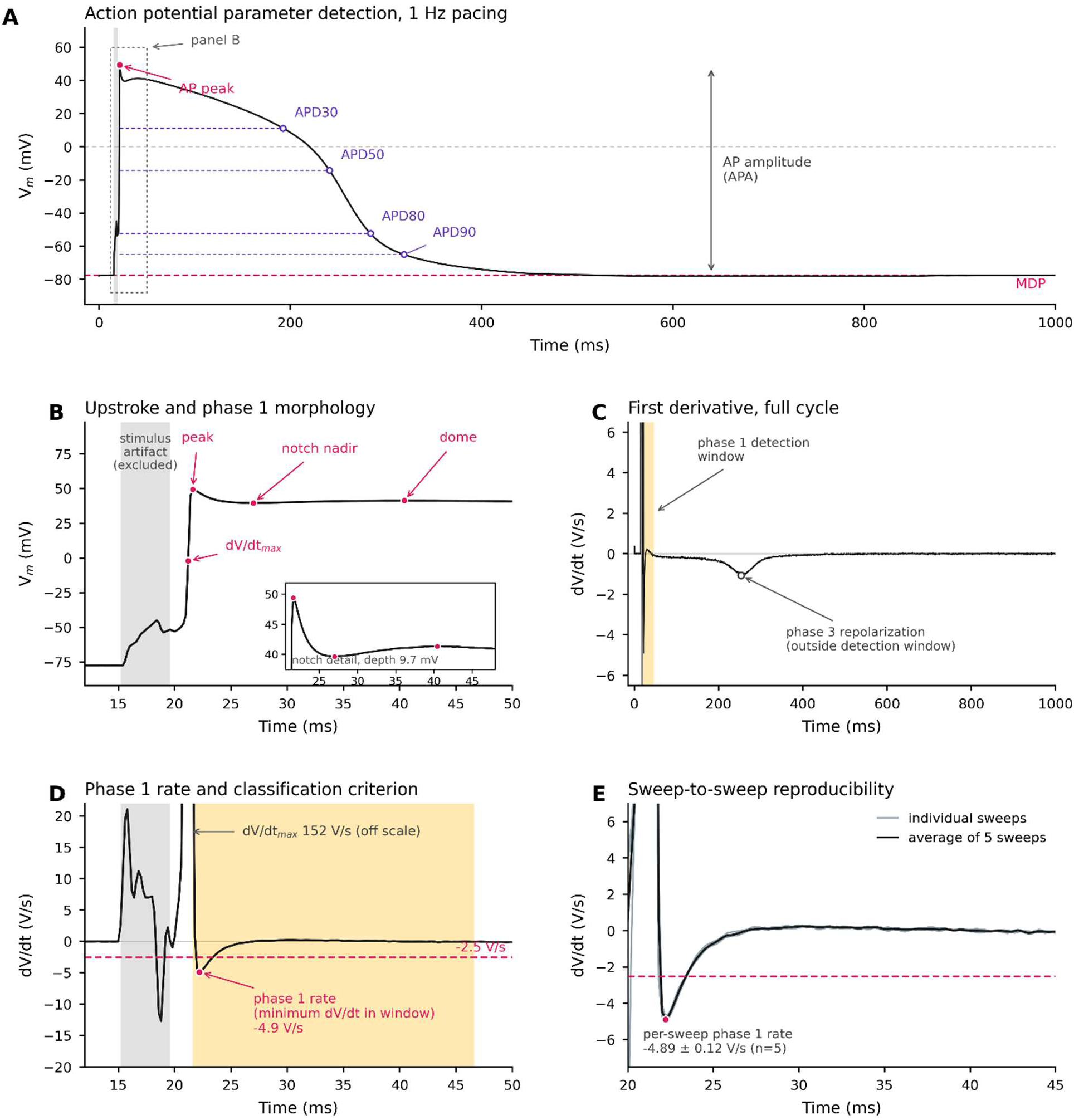
Automated detection of action potential parameters and classification of phase 1 repolarization. **A.** presentative averaged action potential (AP). The maximum diastolic potential (MDP, dashed magenta line) is the median mbrane potential during the final 200 ms of the cycle, and AP amplitude (APA) as the difference between the AP peak and P. AP duration at 30, 50, 80 and 90 % repolarization (APD30, APD50, APD80, APD90) is measured from the AP peak to the t crossing of the corresponding repolarization level (purple dashed lines and open circles). Grey shading marks the stimulus fact, which is excluded from all analyses. The dashed box indicates the region enlarged in B. For this cell: MDP −88.9 mV or correction (−77.7 mV uncorrected), APA 127.1 mV, APD30 171 ms, APD50 219 ms, APD80 262 ms, APD90 297 ms. **B.** argement of the upstroke and early repolarization. Maximum upstroke velocity (dV/dt_max) is identified as the largest positive/dt occurring after the stimulus artifact; the AP peak, the notch nadir (voltage minimum within the phase 1 detection window) the dome (voltage maximum following the nadir) are detected within the same window. Inset, notch region at expanded scale; ch depth, measured from AP peak to nadir, is 9.7 mV in this cell. **C.** First derivative of the averaged action potential over the cycle. Yellow shading indicates the phase 1 detection window, extending 25 ms from the AP peak; grey shading marks the luded stimulus artifact. The open circle marks the negative dV/dt deflection produced by phase 3 repolarization, which falls side the detection window and is therefore not considered in the phase 1 classification. **D.** Enlargement of the derivative around upstroke. The phase 1 repolarization rate is defined as the most negative dV/dt within the detection window (magenta circle, 9 V/s in this cell). The dashed magenta line marks the −2.5 V/s classification threshold; cells with a phase 1 rate more negative n this threshold are scored as displaying prominent phase 1 repolarization. dV/dt_max (152 V/s) lies above the plotted range. The five individual sweeps (grey) and their aligned average (black) over the phase 1 window, showing sweep-to-sweep roducibility of the detected phase 1 rate (−4.89 ± 0.12 V/s, mean ± SD, n = 5 sweeps).

**Figure S22.**
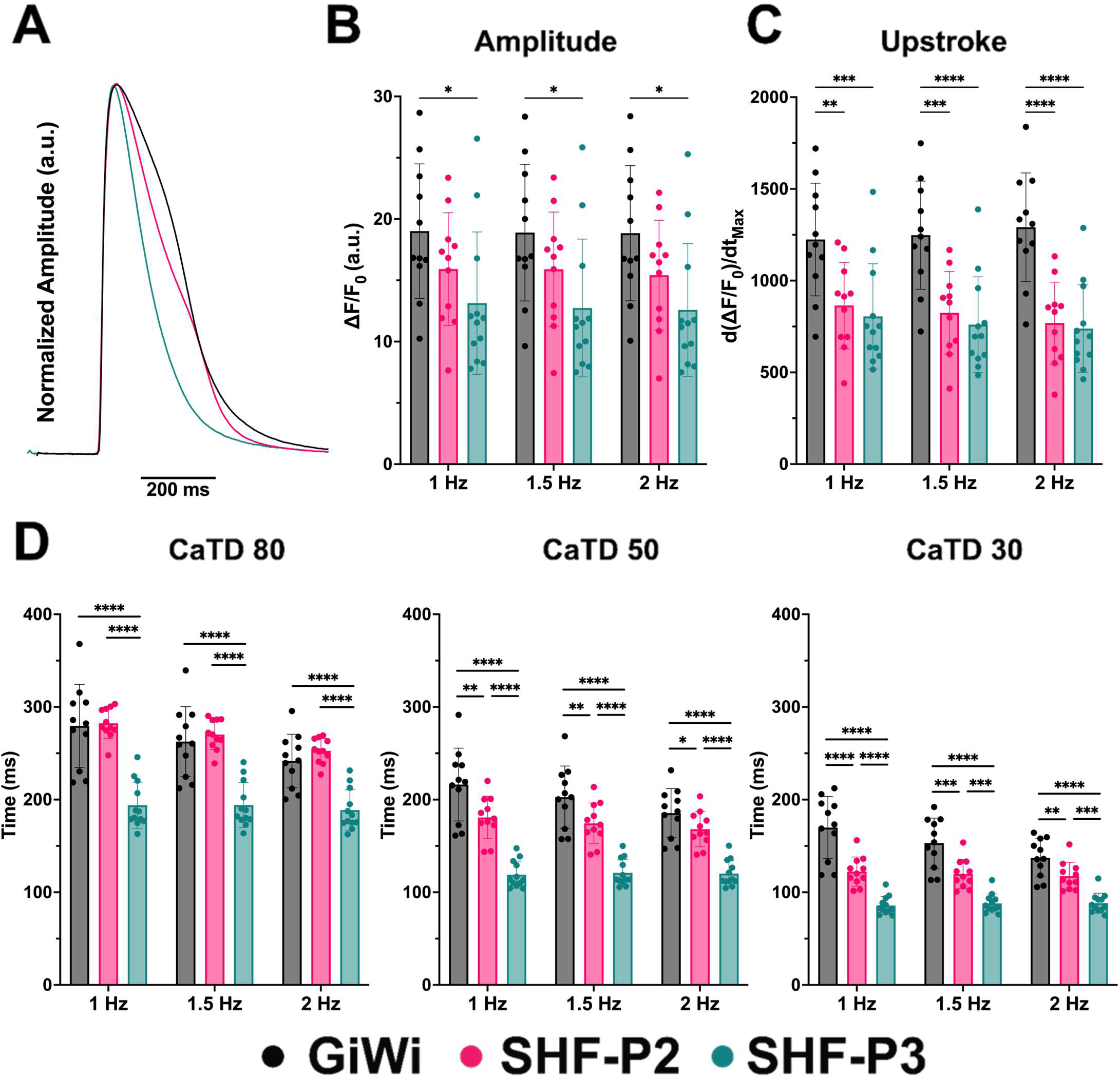
Calcium transient characterization of GiWi-, SHF-P2-, and SHF-P3-derived diomyocytes using optical calcium imaging. **A.** Representative calcium transient traces. **B.** Calcium nsient amplitude. **C.** Peak upstroke velocity. **D.** Calcium transient duration at 80%, 50%, and 30% olarization (CaTD80, CaTD50, and CaTD30) measured at pacing frequencies of 1, 1.5, and 2 Hz. Data presented as mean ± SEM. *n* = 11 for GiWi, *n* = 11 for SHF-P2, and *n* = 12 for SHF-P3. *P* values were ermined by one-way ANOVA with Tukey’s multiple comparisons test. \**P* < 0.05; \*\**P* < 0.01; \*\*\**P* < 01; \*\*\*\**P* < 0.0001.

## Methods

### Human PSC culture

Human healthy iPSC lines DF19-9-11T and WC032i-6007-1 (WiCell Research Institute), Brugada Syndrome disease iPSC line BRSM1c16 (SCN5A D356N), and a hESC reporter line hES3-TBX5^TdTomato/W^/NKX2-5^eGFP/W1^ were used in this study. hPSCs were cultured on Matrigel (GFR, Corning) coated 6-well plates in StemFlex medium (Thermo Fisher Scientific). Cells were passaged using Versene solution (Thermo Fisher Scientific) every 4-5 days. Cells were washed with DPBS without Ca^++^/Mg^++^ (Gibco) and incubated with 1 ml/well Versene solution at 37°C for 3 minutes. Aspirating Versene solution and adding 1ml/well mTeSR1 medium to gently pipette up and down and scrape to cell clusters. Cell clusters were pellet down by centrifugation at 1000 rpm for 3 minutes, resuspended in StemFlex medium, and plated in 6-well plate at the split ratio of 1:6 to 1:18. Mycoplasma tests were routinely performed (monthly) for all the cell culture in this study.

### GiWi protocol

Human PSCs were dissociated with 1 ml/well Versene solution (Thermo Fishser Scientific) at 37°C for 5 minutes and seeded on Matrigel coated 6-well plates at the density of 200,000 cells/cm^2^ in mTeSR1 medium supplemented with 10 μM ROCK inhibitor (Y-27632) (Biogems). Cells were cultured for 5-6 days in mTeSR1 medium (WiCell Research Institute) until reaching over 100% confluence when differentiation started (day 0). At day 0, the medium was changed to 2.5 ml RPMI+B27 without insulin (Gibco) and supplemented with 12 µM CHIR99021 (Biogems), and cells were treated with CHIR99021 in the medium for 24 hours (day 1). At day 1, the medium was changed to 3 ml RPMI+B27 without insulin and cells were cultured in this medium for 48 hours (day 3). At day 3, a combined media was prepared by collecting 1.5 ml of the spent media and mixing with same volume of fresh RPMI+B27 without insulin and supplemented with 5 µM IWP2 (Biogems), and cells were treated with IWP2 in the media for 48 hours (day 5). At day 5, media was changed to 2.5 ml RPMI+B27 without insulin. At day 7 the media was changed to 2.5 ml RPMI+B27 with insulin (Gibco), and cells were fed every other day with the same medium and differentiated until day 20.

### SHF-P1 and P2 protocols

Human PSCs were dissociated with 1 ml/well Versene solution (Thermo Fisher Scientific) at 37°C for 5 minutes and seeded on Matrigel coated 6-well plates at the density of 200,000 cells/cm^2^ in mTeSR1 medium supplemented with 10 μM ROCK inhibitor (Y-27632) (Biogems). Cells were cultured for 5-6 days in mTeSR1 medium (WiCell Research Institute) until reaching over 100% confluence when differentiation started (day 0). At day 0, the medium was changed to 2.5 ml RPMI+B27 without insulin (Gibco) and supplemented with 12 µM CHIR99021 (Biogems) (SHF-P1 protocol) or supplemented with 1ug/ml insulin (Sigma) (SHF-P2 protocol). and cells were treated in this medium for 24 hours (day 1). At day 1, medium was changed to 2.5ml RPMI+B27 without insulin and cells were cultured in the medium for 30 hours (day 2.75). At day 2.75, the medium was changed to 2.5ml CFBM-medium (**Supplementary Table S1**) supplemented with 75 ng/ml bFGF (Waisman Biomanufacturing), and the cells were treated with bFGF in the medium for 24 hours. At day 3.75, the medium was changed to 2.5ml of the CFBM-medium supplemented with 5 µM IWP2 (Biogems), and cells were treated with IWP2 in the medium for 48 hours (day 5.75). At day 5.75, cells were fed with 2.5ml CFBM+ medium (**Supplementary Table S1**), and cells were fed every other day with the same medium and differentiated until day 20.

### SHF-P3 protocol

Human PSCs were dissociated with 1 ml/well Versene solution (ThermoFishser) at 37°C for 5 minutes and seeded on Matrigel coated 6-well plates at the density of 200,000 cells/cm^2^ in mTeSR1 medium supplemented with 10 μM ROCK inhibitor (Y-27632) (Biogems). Cells were cultured for 5-6 days in mTeSR1 medium (WiCell Research Institute) until reaching over 100% confluence when differentiation started (day 0). At day 0, the medium was changed to 2.5 ml RPMI+B27 without insulin (Gibco) and supplemented with 12 µM CHIR99021 (Biogems), and cells were treated in this medium for 24 hours (day 1). At day 1, medium was changed to 2.5ml RPMI+B27 without insulin (Gibco) and cells were cultured in the medium for 30 hours (day 2.75). At day 2.75, the medium was changed to 2.5ml CFBM-medium (**Supplementary Table S1**) supplemented with 75 ng/ml bFGF (Waisman Biomanufacturing) and 500 nM Retinoic Acid (ATRA, Sigma), and the cells were treated with bFGF and ATRA in the medium for 24 hours.

At day 3.75, the medium was changed to 2.5ml of the CFBM-medium supplemented with 5 µM IWP2 (Biogems) and 500 nM ATRA, and cells were treated with IWP2 and ATRA in the medium for 48 hours (day 5.75). At day 5.75, cells were fed with 2.5ml CFBM+ medium (**Supplementary Table S1**), and cells were fed every other day with the same medium and differentiated until day 20.

### Flow cytometry

Singel cells were isolated at day 3.75 of the GiWi and SHF-P1, P2 and P3 protocols for analysis by flow cytometry. The day 3.75 differentiated cells were dissociated by TrypLE 10X (Gibco) treatment for 7-10 minutes at 37°C, and pipet up and down with 5 ml glass pipets for ∼40 times to isolate single cells. Isolated single cells were labeled with CXCR4 antibody for flow cytometry. The day 20 differentiated cells of the GiWi, SHF-P1, P2 and P3 protocols were dissociated by incubation with TrypLE 10X (Thermo Fisher) for 7 minutes and then 0.25% trypsin-EDTA (Gibco) plus 2% chick serum (Sigma) for another 7 minutes at 37°C, and single cells were isolated by pipetting up and down with 5 ml glass pipets for ∼40 times. Single cells were harvested to 15 ml conical tube, equal volume of EB20 medium^2^ was added to resuspend the cells. Cells were counted by hemocytometer, 0.5-1 million of cells/sample were used for antibody labeling. For labeling surface markers, cells were live labeled and fixed in 1% paraformaldehyde at 37°C water bath for 10 minutes in the dark. For labeling intracellular markers, cells were fixed in 1% paraformaldehyde and permeabilized in ice-cold 90% methanol for 30 minutes on ice. Cells were washed once in FACS buffer (DPBS without Ca^++^/Mg^++^, 0.5% BSA, 0.1% NaN3), centrifuged and resuspended in about 50 μl FACS buffer with (for intracellular markers) or without 0.1% Triton. For labeling surface markers, appropriate amount of the conjugated antibodies according to the manufacturer’s instruction were added to the cells in FACS buffer to make the final volume of 100 μl with FACS buffer, and incubated at room temperature in dark for 30 minutes. Cells were washed once with 3 ml FACS buffer and resuspended in 300 – 500 μl FACS buffer for analysis. For labeling intracellular markers, the primary antibody was diluted in 50 μl /sample FACS buffer plus 0.1% Triton and aliquoted to each sample for a total sample volume of 100 μl. Samples were incubated with the primary antibodies overnight at 4°C. Please refer to **Supplementary Table S3** for the primary antibodies used in this study. For the secondary antibody labeling, cells were washed once with 3 ml FACS buffer plus 0.1% Triton after the primary antibody labeling, centrifuged, and supernatant was aspirated but leaving ∼ 50 μl. Secondary antibody specific to the primary IgG isotype was diluted in FACS buffer plus Triton in a final sample volume of 100 μl at 1:1000 dilution. Samples were incubated at room temperature for 30 minutes in dark, washed in FACS buffer plus Triton and resuspended in 300 – 500 μl FACS buffer plus Triton for analysis. Data were collected on a Attune Nxt (Thermo Fisher Scientific) flow cytometers and analyzed using FlowJo.

### Immunocytochemistry

For imaging with EVOS microscope (Life Technologies), cells were cultured in either 6-well or 24-well plates. Cells were fixed with 4% paraformaldehyde for 15 minutes at room temperature, and permeabilized in 0.2% Triton X-100 (Sigma) for 0.5-1hr at room temperature. After fixing and permeabilization, cells were blocked with 5% non-fat dry milk (Bio-Rad) in 0.2% Triton X-100 solution and incubated for 1.5-2 hrs at room temperature on a rotator followed by two washes with DPBS. Primary antibodies (**Supplementary Table 3**) were added in 1% BSA in DPBS solution with 0.1% Triton X-100, and incubated overnight at 4°C. Cells were washed with 0.2% Tween 20 in DPBS twice and DPBS twice. Secondary antibody specific to the primary IgG isotype were diluted (1:1000) in the same solution as the primary antibodies and incubated at room temperature for 1hr in dark on a rotator. Cells were washed with 0.2% Tween 20 in DPBS twice and DPBS twice. Nuclei were labeled with Hoechst or DAPI.

### Quantitative RT-PCR

Cell samples were collected using 0.25% trypsin-EDTA (Invitrogen) to remove the cells from cell culture plates. Total RNA was purified using RNeasy Mini Kit (QIAGEN 74104). Any genomic DNA contamination was removed by RNase-Free DNase Set (QIAGEN 79254) with the RNeasy columns or by DNase I (Invitrogen) treatment for 15 minutes at room temperature. 500 ng of total RNA was used for Oligo (dT)_20_ – primed reverse transcription using SuperScript™ IV VILO™ Master Mix (Thermo Fisher 11756050). Quantitative RT-PCR was performed using 2x TaqMan™ Fast Advanced Master Mix (Thermo Fisher 4444557) and TaqMan Gene Expression Assays (Thermo Fisher, **Supplementary Table S4**) in triplicate for each sample and each gene. 0.5 μl of cDNA from RT reaction was added as template for each qRT-PCR reaction. The expression of genes of interest was normalized to *GAPDH*.

### Single-cell RNA sequencing, transcriptomic and GO term analyses

#### Sample preparation

DF19-9-11T iPSCs were differentiated using GiWi-12CHIR, GiWi-8CHIR, SHF-P2 and SHF-P3 protocols. Two progenitor samples that were differentiated at day 3.75 in GiWi-12CHIR and SHF-P2 protocols, and four samples of day 20 differentiated cells from GiWi-12CHIR, GiWi-8CHIR, SHF-P2 and SHF-P3 protocols were harvested and fixed at the specific timepoint during differentiation. The day 3.75 and day 20 differentiated cells were harvested from the same batch of differentiation, and the differentiation were performed in parallel for all the protocols. Differentiated cells at specific timepoints were harvested as the protocol below.

1. Wash with 2ml PBS (without calcium & magnesium, Corning, 21-040-CV)) for each well (6-well plate).
2. Add 1ml/well TrypLE 10x (Gibco), incubate for 5-7 min in incubator before cells detaching.
3. Aspirate the TrypLE 10x without disturbing the cells, add 1ml Trypsin-EDTA (0.25%) plus 2% chick serum, incubate for another 5-7 min in incubator.
4. Add RNase inhibitor (Millipore Sigma 3335402001, 40 U/ul) 1 ul/million cells and DNase I (RNase-Free, Qiagen 79254 (2.7 Kunitx Unit/ul), 10-50 U/ml to each well before harvest.
5. Harvest the cells using 5 ml glass pipette and transfer to 15ml conical tube.
6. Using 5 ml glass pipette, pipet up and down 35 times to dissociate the cells.
7. Add 2-3 ml cold EB20 media ^2^, pipet up and down to mix 8-10 times.
8. Take 20 ul aliquot, add 20 ul Trypan blue to count cells.

#### Sample fixing

Parse Evercode™ WT Mini v2 kit was used to fix the cells following the Parse User Manual (**Supplementary File 7**). Fixed cells were stored at −80°C until all samples were collected.

#### Single-cell RNA-seq library preparation and sequencing

Fixed cells were submitted to the University of Wisconsin Biotechnology Center Gene Expression Core (RRID: SCR_017757) for single-cell RNA-seq library preparation using the Parse Evercode WT Kit and subsequent sequencing by the DNA Sequencing Facility (RRID: SCR_017759).

Single-cell transcriptomic libraries were generated using the Evercode WT Mini V2 Kit (Parse Biosciences) according to the manufacturer’s protocol (v2.0.1), targeting approximately 3,000 cells per sample (approximately 20,000 cells total). Fixed cells were thawed, counted using a Luna FX7 automated cell counter with propidium iodide staining, and loaded into 96-well plates for combinatorial barcoding. Cells underwent three rounds of in situ split-pool barcoding, including reverse transcription and successive ligation reactions to introduce well-specific barcodes. Barcoded cells were subsequently pooled, lysed, and divided into two sublibraries prior to cDNA capture using streptavidin beads.

Following template switching and amplification, cDNA libraries were prepared through fragmentation, adapter ligation, and a fourth round of indexing PCR using unique dual i7/i5 indices. Libraries were purified by SPRI bead-based size selection, quantified with a Qubit 4 Fluorometer using the dsDNA High Sensitivity Assay, and assessed for fragment size distribution on an Agilent 4200 TapeStation using D1000 ScreenTape. Each sublibrary was normalized to 2 nM and pooled at equimolar concentrations.

Libraries were sequenced on an Illumina NovaSeq X Plus using paired-end chemistry with the following read configuration: 66 bp Read 1, 8 bp i7 index, 8 bp i5 index, and 86 bp Read 2. Sequencing was performed to a target depth of approximately 60,000 reads per cell.

#### Single-cell RNA-seq transcriptomic analysis

The Parse matrix data was analyzed using the Seurat package (https://satijalab.org/seurat/authors#citation), Seurat V3 ^3^. After visual QC of the single-cell data, the following parameters were used to select high quality cells: number of detected genes per cell between 500 and 8000, total number of RNA molecules (UMIs) less than 60000, percentage of mitochondrial gene expression <40%. The filtered cells were processed using standard Seurat workflow, including log-normalization of UMI counts, SCTransform, principal component analysis, UMAP on first 10 principal components, computation of shared nearest neighbor (SNN) graph, and Louvain graph based clustering at different resolutions (0.25, 0.5, 0.8). The results were visualized using cellxgene VIP (https://www.biorxiv.org/content/10.1101/2020.08.28.270652v2) and cell types were manually assigned to clusters at 0.5 resolution using known marker genes. The Wilcoxon Rank Sum test is used for differential gene expression analysis.

#### GO term analysis

Was performed using the open source g:Profiler (https://biit.cs.ut.ee/gprofiler/gost). The input to g:Profiler is the DEGs for each cluster. The output includes enriched terms in MF (Molecular Function), BP (Biological Process), CC (Cellular Component), KEGG, REAC and WP pathways.

### Single cell patch-clamp electrophysiology

#### Single cell perforated patch clamp recordings

Single hPSC-CMs were patch-clamped 3–7 days after sparse replating onto glass coverslips. Recordings were performed at 37 °C in the perforated-patch current-clamp configuration, which was chosen in preference to ruptured whole-cell to preserve intracellular Ca²⁺ handling, native CaMKII signaling and second-messenger pathways that are dialysed away under conventional EGTA- or BAPTA-buffered conditions and that are known to influence action potential morphology in cardiomyocytes. The extracellular (bath) solution contained (in mM): 154 NaCl, 4 KCl, 1 MgCl₂, 2 CaCl₂, 5 HEPES and 5.5 D-glucose (pH 7.4, adjusted with NaOH). The intracellular (pipette) solution contained (in mM): 125 K-aspartate, 20 KCl, 10 NaCl, 5 Na₂-ATP and 10 HEPES (pH 7.2, adjusted with KOH), supplemented with 0.22 mM amphotericin B freshly diluted from a DMSO stock immediately before each experiment. Patch pipettes were pulled from borosilicate glass capillaries to a tip resistance of 3–5 MΩ. Action potentials were evoked by 2-ms depolarizing current injections (0.5–1.5 nA) at 1 Hz and recorded with an Axopatch 200B amplifier (Molecular Devices) coupled to a Digidata 1440A digitizer (Molecular Devices); signals were sampled at 10 kHz and low-pass filtered at 5 kHz (4-pole Bessel). All reported membrane potentials were corrected offline for the measured liquid junction potential (−11.2 ± 0.9 mV).

#### Assessment of phase 1 repolarization

The transient outward potassium current, I_to_, was not recorded directly in this study. Instead, the prevalence of I_to_-dependent early repolarization was assessed at the level of action potential morphology, using the rate of repolarization immediately following the upstroke as a surrogate. Classification was performed once per cell. For each cell, five consecutive action potentials recorded during steady-state 1 Hz pacing were analyzed. Cells with a phase 1 repolarization rate more negative than −2.5 V/s were classified as displaying prominent phase 1 repolarization; all others were classified as lacking it. This threshold was selected empirically as the value separating the two modes of the phase 1 rate distribution across the pooled dataset (Supplemental Figure 20). Because this measure reflects action potential morphology rather than current amplitude, it indexes the prevalence of prominent phase 1 repolarization across a population and does not provide a quantitative estimate of I_to_ density in individual cells.

#### Carbachol challenge

To functionally distinguish atrial from ventricular cardiomyocyte identity, GiWi-lvCM, SHF-P2–rvCM and SHF-P3–aCM were challenged with 10 µM carbachol (CCh; Sigma-Aldrich, C4382). Action potentials were recorded continuously during a 1-min baseline period at 1.5 Hz pacing, followed by 3 min of carbachol superfusion and a 3-min washout at the same pacing frequency. A pacing rate of 1.5 Hz was used for this protocol, rather than the 1 Hz used for baseline AP characterization, to ensure reliable capture in SHF-P3–aCM, which display the highest spontaneous beating rate of the three preparations and would otherwise be prone to escape beats during the extended baseline–drug–washout recording. Only cells that maintained stable access resistance, maximum diastolic potential and AP morphology throughout the entire baseline–drug–washout protocol were retained for analysis. Action potential parameters (maximum diastolic potential, APD50, APD80, dV/dt_max) were extracted offline from the averaged AP of the last 30 s of each condition using BAPTA (Beat Analysis and Patch-clamp Trace Assessment; Leonov et al., bioRxiv 2023).

#### Statistical analysis of patch-clamp data

Group data are reported as mean ± SEM unless stated otherwise. Comparisons of unpaired single-cell parameters across the three CM populations (maximum diastolic potential, APD80, dV/dt_max and cell capacitance; Figure 7B–E) were performed by ordinary one-way ANOVA followed by Tukey’s multiple-comparisons test. The incidence of prominent phase 1 repolarization (Figure 7F) was compared across the three populations by Pearson’s chi-square test of independence on a 3 × 2 contingency table of cell counts, followed by pairwise Fisher’s exact tests with Holm–Bonferroni correction for three comparisons; effect size is reported as Cramér’s V, and proportions are given with Wilson 95 % confidence intervals. Paired drug-response data (baseline / carbachol / washout; Figure 7G–I) were analysed by repeated-measures one-way ANOVA for GiWi-lvCM and SHF-P2–rvCM, and by a mixed-effects model (REML) for SHF-P3–aCM to accommodate one missing washout value, in each case followed by Šídák’s multiple-comparisons test. All analyses were performed in GraphPad Prism v11. A two-tailed adjusted p < 0.05 was considered statistically significant.

### Micropatterned PDMS fabrication for optical mapping

Micropatterns were etched, using UV photolithography, onto a silicon wafer with the according dimensions: lanes with 5 µm in width, 5 µm in pitch, and 8 µm in depth. This silicon mold was cleaned with isopropanol, baked at 150 °C for 2hrs, and coated with PFOCTS (Sigma), a volatile, silanization agent, in a vacuum chamber overnight that would render the surface anti-adhesive. SYLGARD™ 527 Silicone Dielectric Gel (Dow) was used to create soft, ∼10kPa, PDMS foundation for the patterns to be transferred onto. Part A and B was mixed in 1:1 in weight, degassed, and then poured onto the silicon mold. Then, the resin was cured at 60 °C for 6 hrs. Cured PDMS was then peeled off the mold while saturated in isopropanol to prevent further adhesion. Using an 8 mm biopsy tool, they were punched into individual micropatterned PDMS chips and placed onto a 15 mm glass coverslip to allow easier manipulation. They were further autoclaved for sterilization, which had stiffened the chips to 18 kPa. Polydopamine (Sigma) coating was utilized as previously published by Sharma et al. to ensure proper attachments of the hPSC-CMs ^4^. Then, Cultrex (R&D Systems) coating was applied before drop-plating a suspension of hPSC-CMs onto the surface.

### Purification of hPSC-CMs and plating onto MP-PDMS chips

The day 20 differentiated hPSC-CMs from the GiWi, SHF-P2, and SHF-P3 protocols were thawed in EB20 media supplemented with 10 µM Blebbistatin and plated onto Synthemax (Corning) coated 6 well plates at 1.7∼2.3 million cells/well density. After 48 hours, the media were changed to their corresponding differentiation media and treated with Nutlin-3a (20 µM), an MDM2 inhibitor, for purification ^5^. Nutlin-3a was removed after 4 days, and the CMs were allowed to recover for 10 days. Purified hPSC-CMs were dissociated using TrypLE™ Select (Gibco) and drop-plated onto MP-PDMS chips at 55,000∼65,000 density in EB20 media supplemented with 10 µM Blebbistatin for monolayer formation. After 48 hrs, LaSR media ^6^ was used to culture all three-protocol derived hPSC-CMs before optical mapping assessments. The timeline of thawing, plating, purification and recovering is shown in the figure below.

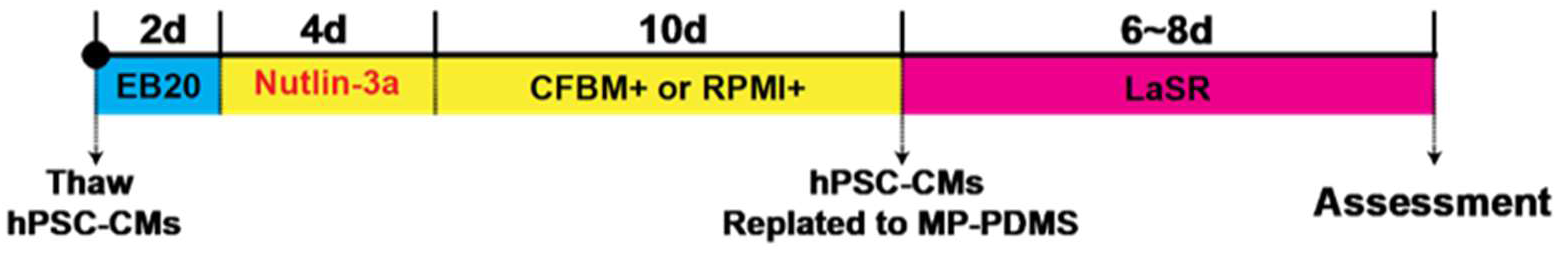

### Optical mapping

Monolayer of hPSC-CMs derived from each protocols (GiWi, SHF-P2, SHF-P3) plated onto MD-PDMS chips were stained with 5 µg/mL Calbryte™ 520 AM (AAT Bioquest) in Tyrode solution for 15 minutes at 37 °C and washed for 15 minutes. The optical mapping system comprised of a perfusion chamber with Tyrode solution flowing laminarly and maintained at 35∼37 °C. Light generated via the excitation LED (X-Cite XYLIS, Excelitas Technologies, CA, USA) was filtered through a dichroic mirror (475/35 nm; Thorlabs, NJ, USA) and emission light from the sample was collected via a long-pass filter (520/35 nm; Thorlabs, NJ, USA), which were captured by a MiCAM Ultima-L CMOS camera (SciMedia, CA, USA) at a 256 x 256 pixels (11.08 mm x 11.08 mm) resolution that temporally recorded calcium transients at 400 frames/sec. These fluorescence signals were processed with BrainVision. (BV) analyzer software (SciMedia, CA, USA). Samples were point stimulated near the center of the monolayer, hovering above yet without directly touching the cells, at varying frequencies (1, 1.5, 2 Hz) to capture parameters such as calcium transient duration (CaTD), upstroke, amplitude, and conduction velocity. Raw transient data are exported as csv files from BV analyzer and analyzed via a custom script on MatLab, while propagation velocity and its representative activation maps are calculated and curated using the BV analyzer.

### Statistics

Sample size varies and was determined based on anticipated scatter of the data, but formal power calculations were not done given the lack of previous comparable datasets from which to estimate variance. Data are presented as mean ± standard error of the mean (SEM). Technical and biological replicates are indicated for each dataset. For datasets with normal distributions, statistical significance was determined by Student’s t-test for two groups or one-way ANOVA for multiple groups with post-hoc test using Tukey methods. Statistical analysis was performed using Origin 2020, P < 0.05 was considered statistically significant.

---

**Supplementary Table S1.**
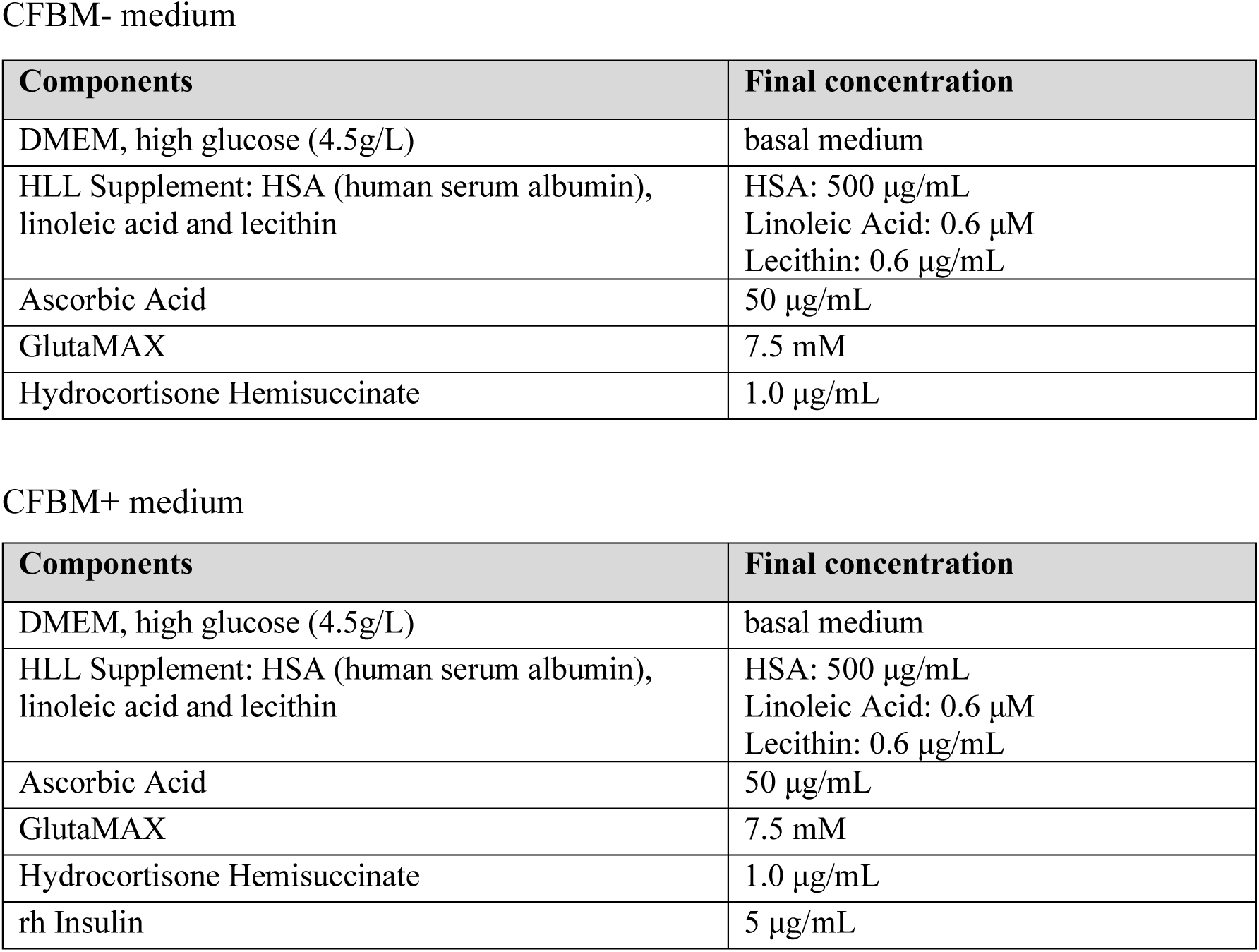
Formulation of CFBM media CFBM-media.

| **Components** | **Final concentration** |
|---|---|
| CFBM- medium |  |
| DMEM, high glucose (4.5g/L) | basal medium |
| HLL Supplement: HSA (human serum albumin), linoleic acid and lecithin | HSA: 500 µg/mL<br>Linoleic Acid: 0.6 µM<br>Lecithin: 0.6 µg/mL |
| Ascorbic Acid | 50 µg/mL |
| GlutaMAX | 7.5 mM |
| Hydrocortisone Hemisuccinate | 1.0 µg/mL |
| CFBM+ medium |  |
| **Components** | **Final concentration** |
| DMEM, high glucose (4.5g/L) | basal medium |
| HLL Supplement: HSA (human serum albumin), linoleic acid and lecithin | HSA: 500 µg/mL<br>Linoleic Acid: 0.6 µM<br>Lecithin: 0.6 µg/mL |
| Ascorbic Acid | 50 µg/mL |
| GlutaMAX | 7.5 mM |
| Hydrocortisone Hemisuccinate | 1.0 µg/mL |
| rh Insulin | 5 µg/mL |

**Supplementary Table S2.**
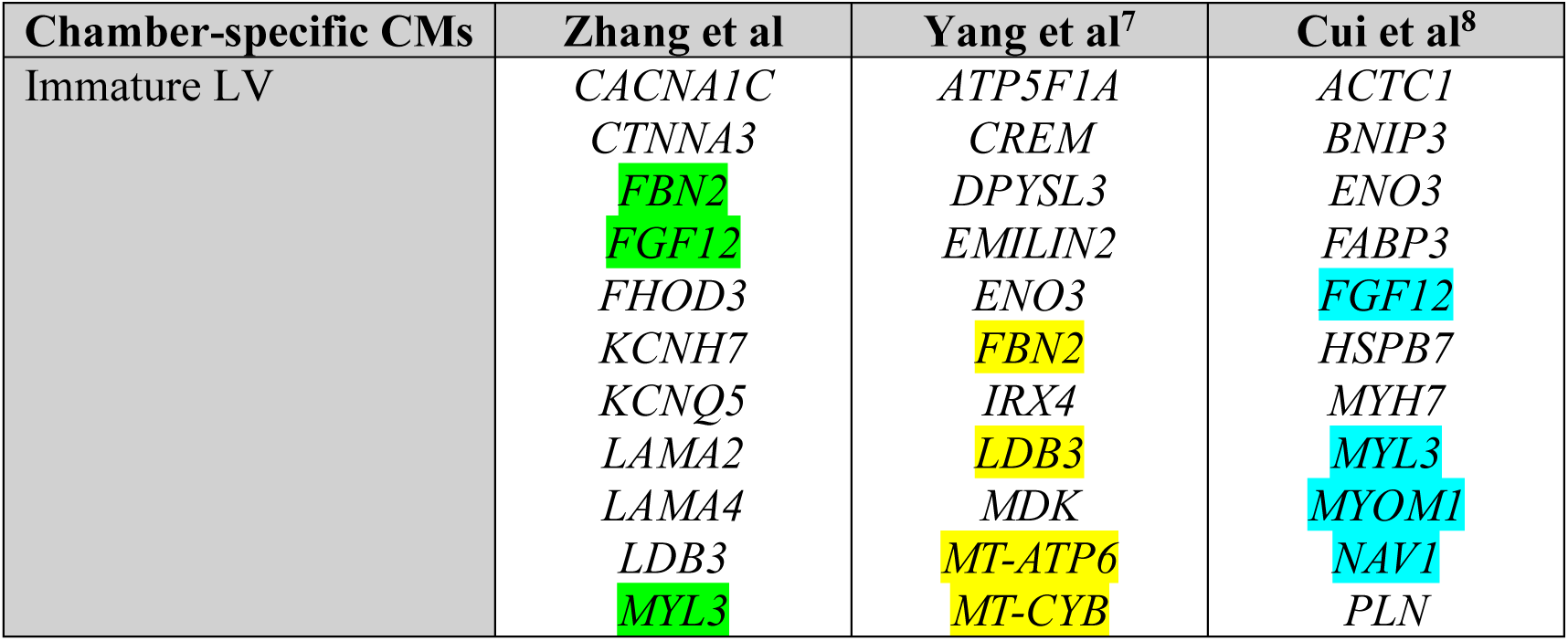
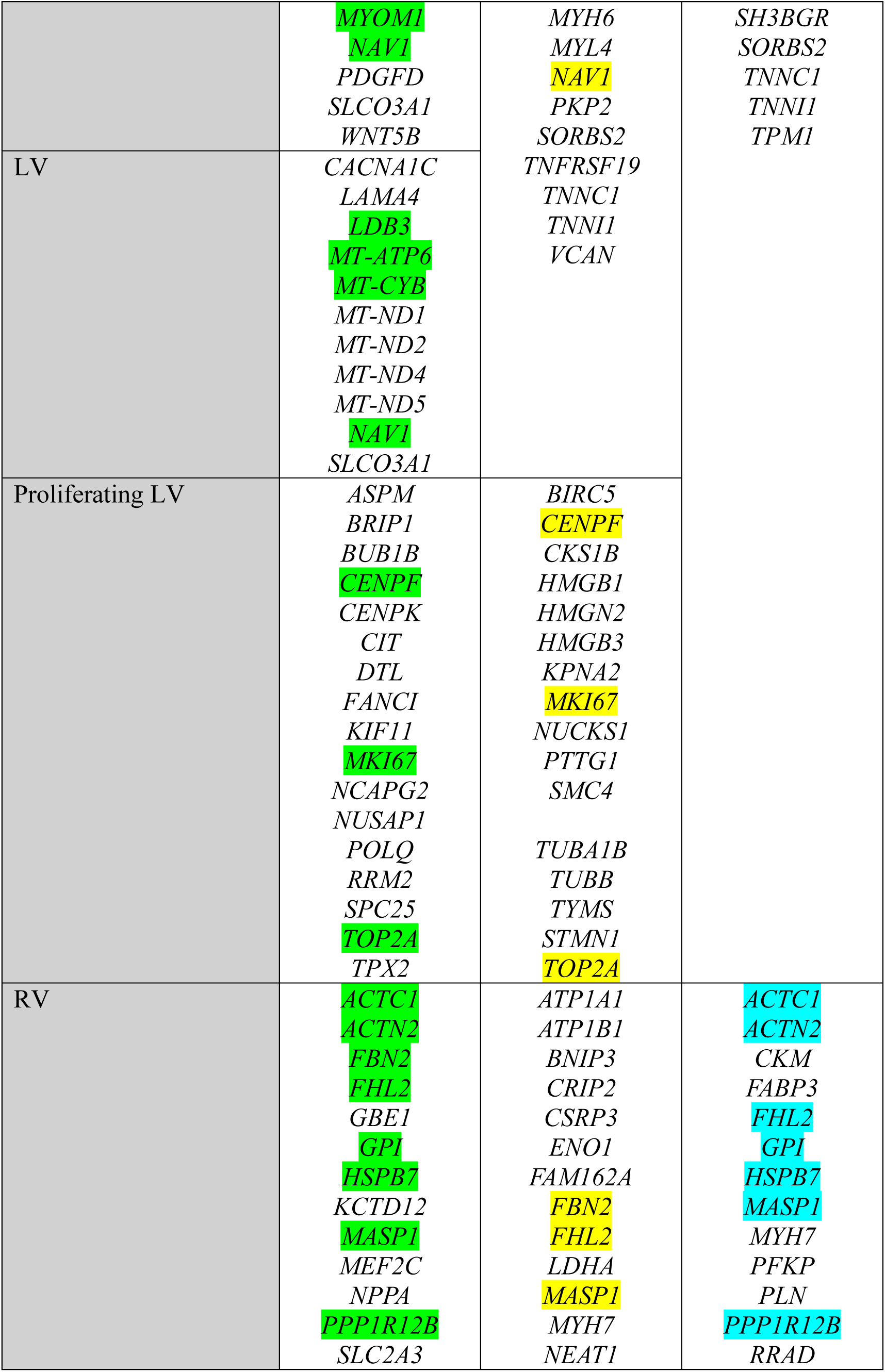
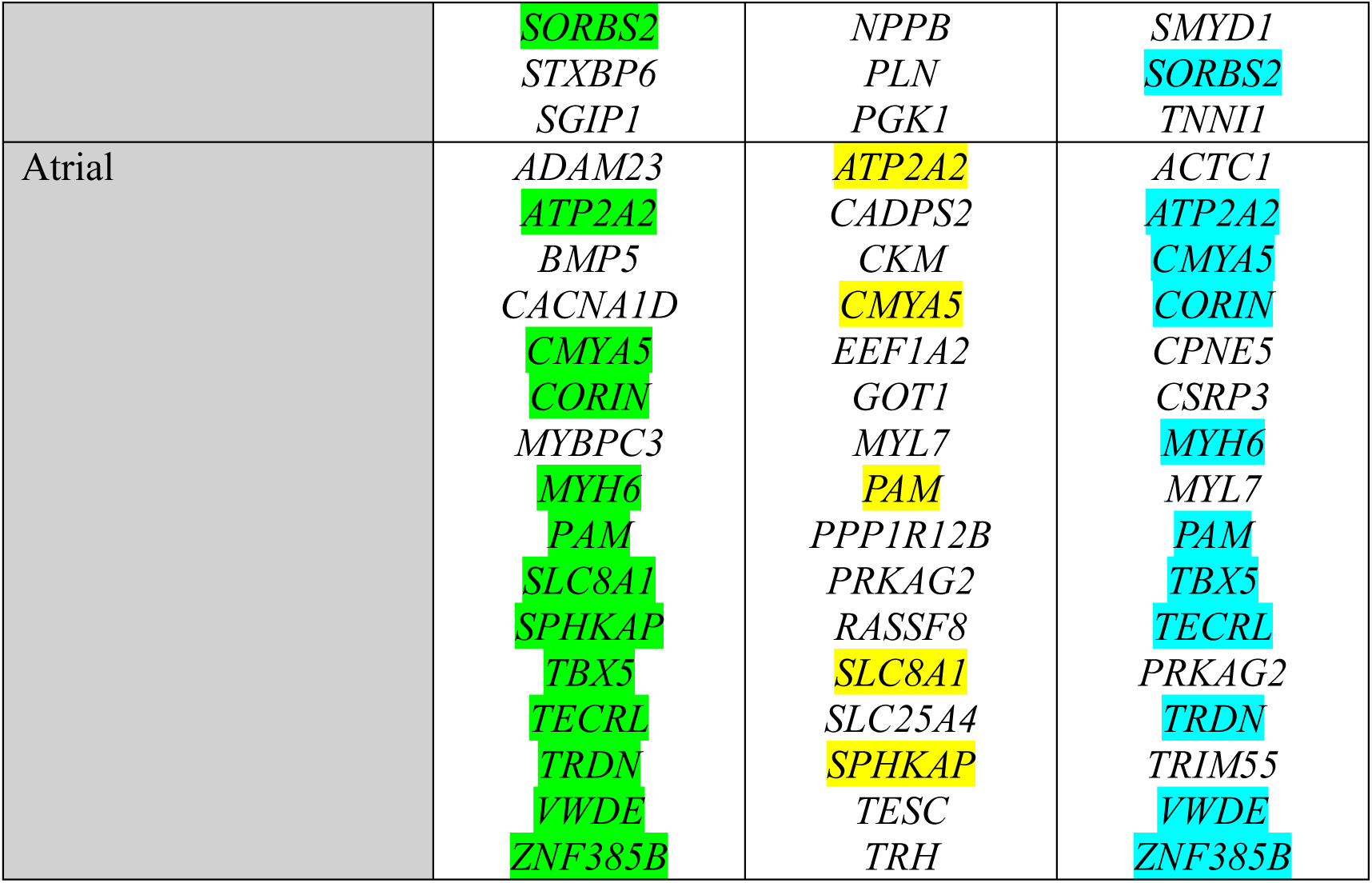
The shared, most significantly upregulated genes in the chamber-specific CMs selected from our scRNA-seq dataset (Zhang et al) and literature as shown (overlapped DEGs are highlighted).

| **Chamber-specific CMs** | **Zhang et al** | **Yang et al<sup>7</sup>** | **Cui et al<sup>8</sup>** |
|---|---|---|---|
| Immature LV | *CACNA1C*<br>*CTNNA3*<br>*FBN2*<br>*FGF12*<br>*FHOD3*<br>*KCNH7*<br>*KCNQ5*<br>*LAMA2*<br>*LAMA4*<br>*LDB3*<br>*MYL3*<br>*MYOM1*<br>*NAV1*<br>*PDGFD*<br>*SLCO3A1*<br>*WNT5B* | *ATP5F1A*<br>*CREM*<br>*DPYSL3*<br>*EMILIN2*<br>*ENO3*<br>*FBN2*<br>*IRX4*<br>*LDB3*<br>*MDK*<br>*MT-ATP6*<br>*MT-CYB*<br>*MYH6*<br>*MYL4*<br>*NAV1*<br>*PKP2*<br>*SORBS2*<br>*TNFRSF19*<br>*TNNC1*<br>*TNNI1*<br>*VCAN* | *ACTC1*<br>*BNIP3*<br>*ENO3*<br>*FABP3*<br>*FGF12*<br>*HSPB7*<br>*MYH7*<br>*MYL3*<br>*MYOM1*<br>*NAV1*<br>*PLN*<br>*SH3BGR*<br>*SORBS2*<br>*TNNC1*<br>*TNNI1*<br>*TPM1* |
| LV | *CACNA1C*<br>*LAMA4*<br>*LDB3*<br>*MT-ATP6*<br>*MT-CYB*<br>*MT-ND1*<br>*MT-ND2*<br>*MT-ND4*<br>*MT-ND5*<br>*NAV1*<br>*SLCO3A1* |  |  |
| Proliferating LV | *ASPM*<br>*BRIP1*<br>*BUB1B*<br>*CENPF*<br>*CENPK*<br>*CIT*<br>*DTL*<br>*FANCI*<br>*KIF11*<br>*MKI67*<br>*NCAPG2*<br>*NUSAP1*<br>*POLQ*<br>*RRM2*<br>*SPC25*<br>*TOP2A*<br>*TPX2* | *BIRC5*<br>*CENPF*<br>*CKS1B*<br>*HMGB1*<br>*HMGN2*<br>*HMGB3*<br>*KPNA2*<br>*MKI67*<br>*NUCKS1*<br>*PTTG1*<br>*SMC4*<br>*TUBA1B*<br>*TUBB*<br>*TYMS*<br>*STMN1*<br>*TOP2A* |  |
| RV | *ACTC1*<br>*ACTN2*<br>*FBN2*<br>*FHL2*<br>*GBE1*<br>*GPI*<br>*HSPB7*<br>*KCTD12*<br>*MASP1*<br>*MEF2C*<br>*NPPA*<br>*PPP1R12B*<br>*SLC2A3*<br>*SORBS2*<br>*STXBP6*<br>*SGIP1* | *ATP1A1*<br>*ATP1B1*<br>*BNIP3*<br>*CRIP2*<br>*CSRP3*<br>*ENO1*<br>*FAM162A*<br>*FBN2*<br>*FHL2*<br>*LDHA*<br>*MASP1*<br>*MYH7*<br>*NEAT1*<br>*NPPB*<br>*PLN*<br>*PGK1* | *ACTC1*<br>*ACTN2*<br>*CKM*<br>*FABP3*<br>*FHL2*<br>*GPI*<br>*HSPB7*<br>*MASP1*<br>*MYH7*<br>*PFKP*<br>*PLN*<br>*PPP1R12B*<br>*RRAD*<br>*SMYD1*<br>*SORBS2*<br>*TNNI1* |
| Atrial | *ADAM23*<br>*ATP2A2*<br>*BMP5*<br>*CACNA1D*<br>*CMYA5*<br>*CORIN*<br>*MYBPC3*<br>*MYH6*<br>*PAM*<br>*SLC8A1*<br>*SPHKAP*<br>*TBX5*<br>*TECRL*<br>*TRDN*<br>*VWDE*<br>*ZNF385B* | *ATP2A2*<br>*CADPS2*<br>*CKM*<br>*CMYA5*<br>*EEF1A2*<br>*GOT1*<br>*MYL7*<br>*PAM*<br>*PPP1R12B*<br>*PRKAG2*<br>*RASSF8*<br>*SLC8A1*<br>*SLC25A4*<br>*SPHKAP*<br>*TESC*<br>*TRH* | *ACTC1*<br>*ATP2A2*<br>*CMYA5*<br>*CORIN*<br>*CPNE5*<br>*CSRP3*<br>*MYH6*<br>*MYL7*<br>*PAM*<br>*TBX5*<br>*TECRL*<br>*PRKAG2*<br>*TRDN*<br>*TRIM55*<br>*VWDE*<br>*ZNF385B* |

**Supplementary Table S3.** Primary antibodies used in flow cytometry (FC) and immunocytochemistry (ICC).

|  |  |  |  |  |
|---|---|---|---|---|
| Human CXCR4 APC-conjugated Antibody | R&D (FAT173a) | Mouse | IgG<sub>2b</sub> | 10ul/1x10<sup>6</sup> cells (FC) |
| PE Mouse anti-Islet-1<br>Clone Q11-465 | BD 562547 | Mouse | IgG<sub>1</sub> | 5µL/1x10<sup>6</sup> cells (FC) |
| NKX2.5 (E1Y8H) Rabbit mAb | Cell Signaling 8792S | Rabbit mAb | IgG | 1:2400 dilution (FC, ICC) |
| Troponin T, Cardiac Isoform Ab-1 (Clone 13-11) | ThermoFisher Scientific (MA5-12960) | Mouse | IgG<sub>1</sub> | 1:200 dilution (FC, ICC) |
| Anti-ACTN2 Antibody<br>(Sarcomeric Alpha Actinin Antibody, monoclonal, EA-53) | Sigma-Aldrich A7811 | Mouse | IgG<sub>1</sub> | 1:100 dilution (ICC) |
| MLC-2A antibody | Synaptic Systems 311011 | Mouse | IgG<sub>2b</sub> | 1:400 dilution (ICC) |
| Myosin Light Chain 2/MLC-2V Polyclonal antibody | Proteintech 10906-1-AP | Rabbit | IgG | 1:200 dilution (ICC) |

**Supplementary Table S4.** Primers for quantitative RT-PCR.

| **Genes** | **TaqMan® Gene Expression Assay ID** |
|---|---|
| *GAPDH* | Hs99999905_m1 |
| *HAND1* | Hs02330376_s1 |
| *TBX5* | Hs01052563_m1 |
| *HCN4* | Hs00975492_m1 |
| *NKX2-5* | Hs00231763_m1 |
| *ISL1* | Hs00158126_m1 |
| *TBX1* | Hs00271949_m1 |
| *TBX20* | Hs00396596_m1 |
| *TNNT2* | Hs00165960_m1 |
| *TNNI3* | Hs00165957_m1 |
| *MYL2* | Hs00166405_m1 |
| *MYH6* | Hs01101425_m1 |
| *MYH7* | Hs01110632_m1 |
| *PLN* | Hs00160179_m1 |
| *IRX4* | Hs00212560_m1 |
| *NPPA* | Hs00383230_g1 |
| *NR2F2* | Hs00819630_m1 |
| *TBX20* | Hs00396596_m1 |
| *HEY2* | Hs01012057_m1 |
| *IRX1* | Hs00411782_m1 |
| *IRX2* | Hs01383002_m1 |
| *OCT4* | Hs01895061_u1 |
| *VIM* | Hs00185584_m1 |

